# High-throughput engineering and modification of non-ribosomal peptide synthetases based on Golden Gate assembly

**DOI:** 10.1101/2025.04.23.650154

**Authors:** Adrian Podolski, Timon A. Lindeboom, Leonard Präve, Janik Kranz, Daniel Schindler, Helge B. Bode

## Abstract

Non-ribosomal peptide synthetases (NRPS) are multimodular enzymes that produce complex peptides with diverse biological activities, potentially being used as clinical drugs. However, the pharmaceutical applications of such natural peptides often require further derivatisation and modification of the peptide backbone, mainly performed by chemical synthesis. A sustainable alternative resembles the *in vivo* engineering of NRPS to change and modify the enzyme properties rationally and, thus, the produced products. The novel NRPS engineering concept, the eXchange Unit Thiolation domain (XUT), allows the efficient modular assembly of different natural NRPS fragments to form hybrid NRPS that produce defined peptides. In this study, we describe a Golden Gate assembly (GGA) method for efficient high-throughput generation of novel and engineered NRPS libraries utilising the XUT concept. This method was applied to generate over 100 novel NRPS with the possibility of changing starter, elongation, and termination modules, respectively. Additionally, we applied this method for targeted modification of the xenoamicin biosynthetic gene cluster (BGC) XabABCD from *Xenorhabdus doucetiae*, resulting in the generation of 25 novel xenoamicin derivatives.

**Graphical Abstract:** A Golden Gate assembly (GGA) method was developed for the efficient assembly of natural and engineered non-ribosomal peptide synthetases (NRPS). This method has enabled the creation of NRPS libraries to generate novel peptides in high-throughput as well as the targeted derivatisation of natural products (NP).

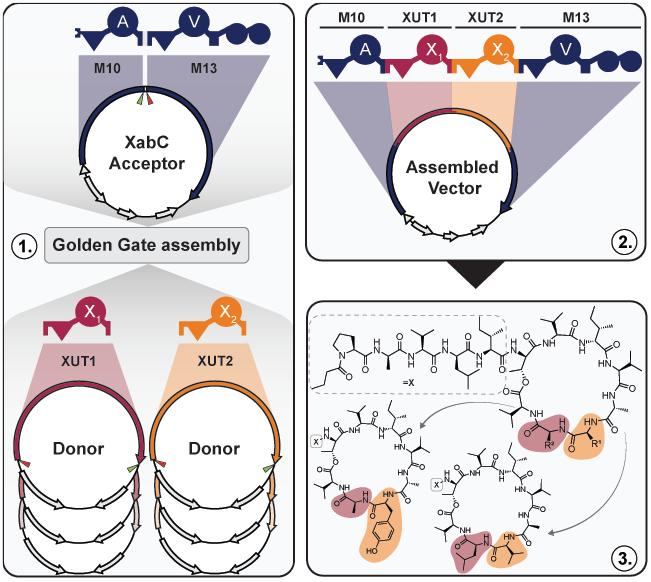

## Introduction

Microbial secondary metabolites or natural products (NPs) are structurally complex molecules with a diverse range of activities and functions. They play an important role in human health as they provide clinically used drugs or precursors for such drugs, including antibiotics or anticancer treatments^[1–3]^. NPs sometimes require optimisation of their selectivity, stability and/or (oral) availability. While chemical synthesis or medicinal chemistry methods are well established to optimise these parameters^[1,4,5]^, they can be challenging and expensive. Conversely, advances in molecular biology, especially synthetic biology, can facilitate such modifications^[6]^. Many clinically used NPs or NPs with promising bioactivities are derived from non-ribosomal peptide synthetases (NRPS) or hybrids between NRPS and polyketide synthases (PKS), which are large, multifunctional enzymes with a characteristic modular nature^[2]^. This modularity offers an ideal entry point for modification, engineering, and subsequent compound derivatisation, even at the backbone level.

Each NRPS module is responsible for selecting, activating, and binding a specific amino acid (AA) to the NRPS and connecting the covalently bound AAs, resulting in a growing peptide chain from the N- to the C-terminal part of the NRPS. NRPS modules can be divided into initiation, multiple elongation, and termination modules. Each elongation module contains a condensation (C) domain that catalyses the peptide bond formation between the upstream and downstream AA. An adenylation (A) domain is responsible for selecting and activating a specific AA, using ATP as a co-substrate. The thiolation (T) domain is posttranslationally activated by a 4’-phosphopantetheinyl transferase and covalently binds the AA to the NRPS. The initiation or starter modules are responsible for the initiation of peptide synthesis and frequently contain a C_Starter_ domain that facilitates the attachment of fatty acids to the N-terminus. Termination modules mainly contain a thioesterase (TE), which is positioned after the T domain and catalyses the release of the peptide as a linear or cyclic (depsi) peptide^[2]^. Several other domains add to the structural complexity of non-ribosomal peptides through additional cyclisation, epimerisation, oxidation, hydrolysis and reduction. Additionally, epimerisation domains can occur as hybrid epimerisation/condensation domains (C/E), performing both functions.

Since modification of the overall modular structure or domain specificity was the goal shortly after their biochemical principles were elucidated, several methods have been developed during the last 30 years. Beginning with the exchange of individual AA in the substrate binding pocket of the A domains, the exchange of partial or complete A domains to change the incorporated AA and the complete exchange of multiple domains and modules ^[7–9]^. While several of these methods function well in specific NRPS systems but show low production titers of the desired peptide when applied to other systems, we and others[10,11] tried to establish more broadly applicable strategies such as the exchange units (XU)^[12]^ and eXchange Unit Condensation domain concepts (XUC)^[13]^ applying fusion sites within the A-C didomain linker or within the pseudo-dimeric structure of the C domain. The most recent eXchange Unit approach, the eXchange Unit Thiolation domain (XUT)^[14]^, identified fusion sites directly before the T domain, described as XUT^I^ or a conserved motif FFxxGGxS in loop 1 and the N-terminus of helix 2 within the T domain, described as XUT^III^ and XUT^IV[14,15]^.

Despite the notable advances in the novel XUT method, allowing the assembly of NRPS and even PKS parts from various origins (since the T domain is the only domain shared between NRPS and PKS)^[14,16]^, cloning and heterologous expression of natural or engineered NRPS is still time-consuming and challenging^[17–20]^. Methods such as transformation-associated recombination (TAR) cloning^[21,22]^ or Gibson assembly^[12–14,23]^ have been frequently used for the assembly of NRPS systems. While these methods are precise and efficient in assembling multiple NRPS-encoding fragments into plasmids, they are time-consuming, with fragments that are often not reusable, making it difficult for high-throughput applications. Unlike previous engineering methods, the XUT concept uses a conserved motif within the T domain, allowing the use of alternative cloning approaches.

Golden Gate assembly (GGA) has become a standard cloning method and provides a baseline for other strategies such as MoClo and GoldenBraid 2.0^[22–24]^. GGA utilises Type IIS restriction enzymes, generating DNA fragments with a three- or four-nucleotide overhang that are joined by a T4 ligase in a one-pot reaction^[24–26]^. As the enzyme cleaving site is adjacent to the recognition site, sequence-independent and thus definable sticky ends can be created, which allow the ordered assembly of such fragments.

Here, we propose a GGA-based approach that facilitates high-throughput cloning for engineered NRPS systems and the generation of modular NRPS libraries. This method combines GGA with the XUT approach, harnessing the two conserved glycines from the XUT^IV^ fusion site for predefined overhangs, thereby enabling a scarless assembly. To demonstrate the functionality of this method and its high-throughput potential for modular-based NRPS libraries, we present the general construction of an acceptor/donor system, as well as its integration into a model system and subsequent derivatisation at the native xenoamicin-producing NRPS.

## Results and discussion

### NRPS Golden Gate assembly design and workflow

The construction of many NRPS encoding plasmids, in previous works, relies on Gibson assembly^[12–14,16,23,27]^. While this method efficiently utilises specific overhangs to ensure successful assembly, it lacks high-throughput potential due to the necessity of designing specific and often unique overhangs for each plasmid assembly. In contrast, a restriction enzyme approach, such as the Golden Gate assembly, obviates the need for such unique overhangs. The XUT^IV^ fusion side, located within the conserved motif “FFxxGGxS” in loop 1 and the N-terminus of helix 2 of the T domain^[14,15]^, provides conserved AAs to create universal overhangs needed for the GGA. Using both conserved glycines in this motif provides enough codons to create sixteen scarless overhangs (Fig. S1), which can be used for acceptor and donor plasmid design, with the aim of combining starter, elongation and termination modules as XUTs.

In initial experiments, acceptor plasmids were constructed that contain a starter and a termination module. For the incorporation of elongation modules, the acceptor plasmid contains two BsaI restriction sites within the XUT^IV^ fusion site, separated by a short spacer sequence (Fig. S2), all being part of the L-arabinose inducible expression plasmid pACYC. BsaI restriction creates the specific overhangs A and B (Fig. S1 and S2) for incorporating one or more elongation units (Fig. 1A). Similarly, donor plasmids contain an elongation module and the BsaI restriction sites at both ends of the XUT sequence (Fig. S2), cloned into the high-copy plasmid pSEVA681^[28,29]^. Additionally, since the GGA is a restriction enzyme-based approach, all native BsaI sites must be removed from all acceptors and donors via the introduction of silent mutations that do not change the encoding enzyme sequence. During the GGA reaction, the acceptor is linearised, while the XUT fragment from the donor is obtained from BsaI digestion. Subsequent ligation results in the insertion of the XUT gene fragment into the acceptor plasmid, thereby creating a single gene encoding for a three modular NRPS (**Fehler! Verweisquelle konnte nicht gefunden werden**.A). The assembled plasmids were then transformed into the producing strain *Escherichia coli* DH10B::*mtaA*^[18]^, which were cultivated at production conditions with the desired inducer(s). The resulting products were analysed using high-performance liquid chromatography mass spectrometry (HPLC/MS) and can be applied to bioassays after extraction (Fig. 1B).

**Figure 1.**
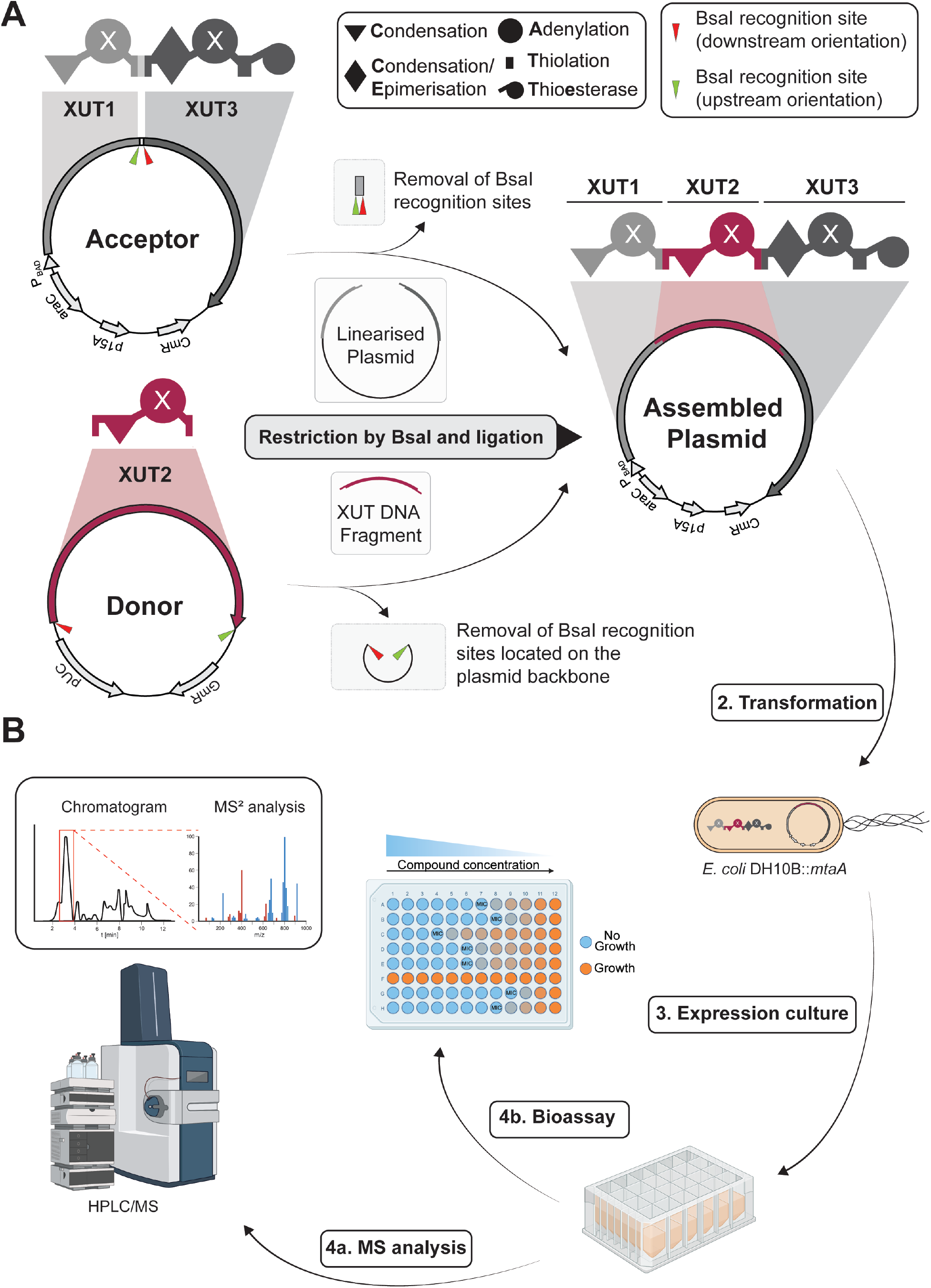
Schematic representation of the Golden Gate assembly (GGA) acceptor and donor plasmids for XUT-based NRPS engineering and the high-throughput workflow. **A**) Schematic representation of an acceptor and donor plasmid with indications of the NRPS-encoding regions highlighted with the respective schematic NRPS illustrated. Both the acceptor and the donor plasmid contain BsaI recognition sites (red and green arrows) to create two different overhangs. The acceptor plasmid harbours a starter module (XUT1; light grey) and a termination module (XUT3; dark grey). The donor plasmid contains a single elongation module (XUT2). Restriction results in the linearisation of the acceptor plasmid while the donor plasmid’s XUT DNA fragment (Bordeaux) is extracted. The final assembled plasmid containing the assembled NRPS system is created upon ligation of both fragments. **B**) After the GGA, the plasmids are transformed into *E. coli* DH10B::*mtaA* for protein expression and peptide production, followed by analysis using HPLC/MS and extract generation for bioassays (Fig. 2B was created with BioRender).

### Elongation module library

Our GGA-based strategy modifies the central motif with a few silent mutations inside the central motif (Fig. S1). In order to test the influence of this minor modification on the peptide production, a set of five NRPS systems designated NRPS-A1-D1 to -D3 and NRPS-A2_D1 to -D2 were cloned with GGA and Gibson assembly. To this end, two acceptor plasmids, NRPS-A1 and NRPS-A2, and three donor plasmids, D1, D2, and D3, were created. Both acceptor plasmids contain the NRPS starter module from the xenolindicin synthethase (XldS) from *X. indica*^[18]^ that mainly incorporates a C_14_ fatty acid at the N-terminus, followed by a glutamine, providing reasonable ionisation and retention times even for smaller peptides in our HPLC/MS system. The termination modules differ in both systems, whereas NRPS_A1 contains the termination module from the szentiamide synthetase (SzeS) from *X. szentirmaii*^[30]^, incorporating a tryptophan for better detection (absorption/excitation [λ_EX_] at ∼280 nm). The second acceptor, NRPS_A2, was designed with the termination module from GameXPeptide synthetase (GxpS) from *Photorhabdus laumondii* TTO1^[31]^, containing a Condensation/Epimerisation (C/E) domain compared to NRPS_A1 and incorporating a leucine. The modules for the first donor subset were chosen from well-studied BGCs, incorporating AAs that are easy to detect in the downstream analysis.

Both versions of each of the five constructs cloned by Gibson assembly (NRPS-1 – NRPS-5) and GGA (NRPS-A1_D1 to -D3 & NRPS-A2_D1 to -D2) were transformed and expressed in *E. coli* DH10B::*mtaA* for peptide production. After analysis using HPLC/MS, almost identical production profiles were obtained regarding both approaches, establishing the viability of GGA for NRPS engineering and the creation of libraries (Tables S7 and S8, Fig. S3).

To advance the GGA concept for the assembly of an NRPS library, the number of donors was increased to six (D1-D6), representing three modules containing C domains and three with C/E domains and the overhangs A and B (Fig. 2A and Fig. S1). The HPLC/MS analysis revealed three different results for the obtained extracts from the expression of the different NRPS systems: (i) The expected peptide was detected, as shown for NRPS-A2_D6 producing C14-Q*a*L **13** (D-amino acids are shown with small letters and in italics; Fig. S4). (ii) Only other derivatives or peptides resulting from module skipping were detected, as shown for NRPS-A2_D4, producing the truncated peptide C14-*q*L **11** (Fig. S5). (iii) No peptide was detected at all. Most of the NRPS-A2_DX systems show the production of the expected peptide, while for NRPS-A1_DX, only NRPS-A1_D3 produced the expected peptide (Fig. 2A and B). In addition, the applicability of the GGA-based NRPS engineering for the generation of a fully randomised NRPS library was tested with the donor set D1-D6 in combination with NRPS-A1. When an equal amount of all donor fragments were used for GGA, an almost equal distribution of possible NRPS variants was detected after transformation into *E. coli* (Fig. 2C).

**Figure 2.**
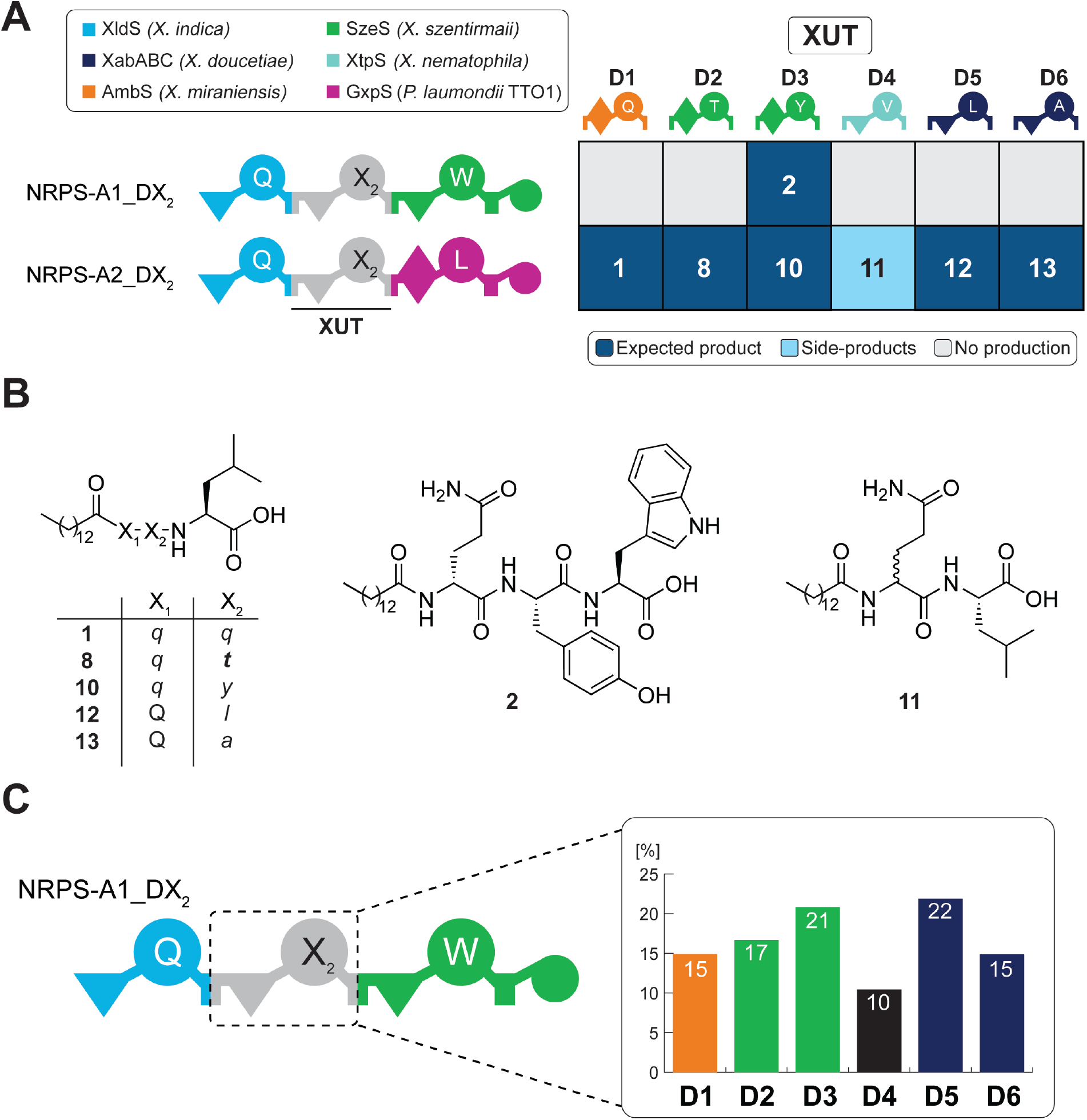
Schematic representation of the results obtained for the initial GGA NRPS module donor and acceptor subset. **A**) Schematic representation of NRPS encoded by the acceptor plasmids and donor plasmids with the overhangs A/B. The NRPS name and origin of the modules are depicted in a legend. A domain specificities are indicated by the one-letter AA code. Representation of the production results from the HPLC/MS analysis of the twelve *NRPS* constructs expressed in *E. coli* DH10B::*mtaA*, where dark blue shows the presence of the expected peptide, light blue shows the presence of side product(s) and grey shows no production. Numbers within tiles refer to the produced peptides (see B). **B**) Chemical structures of the produced peptide sequences from the library are shown in A. AAs are shown in the one-letter code (X_1_ for glutamine and X_2_/X_3_ for the incorporated AAs). Small-italic letters indicate D-amino acids, and small-bold-italic letters represent D-*allo*-amino acids. **C**) Donor incorporation frequency determined by sequencing the one-tube GGA approach for the donors D1-D6 in NRPS-A1 in a randomised library approach. For NRPS domain explanations, see Fig. 1.

In order to test if multiple modules can be incorporated at different positions of the NRPS, the subset of donors (Fig. 2A) containing the overhangs A and B were cloned as novel donors comprising the overhangs A/C (D7-D12) and C/B (D16-21). This approach facilitates the implementation of two donor fragments, thus creating 36 distinct NRPS systems, which were analysed as previously described (Fig. 3A, black box). HPLC/MS analysis results show that 83% of the construct produced functional NRPS, with 61% of them indeed producing the expected peptide (Fig. 3A & Fig. 4). Expression of some NRPS, like NRPS-A2_D7_D21, produced not only the expected peptide **17** but also truncated versions such as **13** and **11**, originating again from module skipping (Fig. S6). As an example of a functional NRPS not producing the expected peptide, NRPS-A2_D9_D16 only produced the truncated peptides **1** and **11** (Fig. S7). Unfortunately, the tyrosine incorporating module from D18, mainly resulted in non-functional NPRS (Fig. 3A). Furthermore, the valine incorporating D10 module also mainly yielded NRPS systems where the expected peptide could not be detected, consistent with the earlier findings from the individual module library (Fig. 2A and 3A). Therefore, additional valine and tyrosine modules were used and added as donors with A/C (D13-D15) and C/B (D22-D24) overhangs. This resulted in the construction of 81 different NRPS constructs in total, showing an increase of functional NRPS as well as the detection of the expected peptide sequences (Fig. 3A).

**Figure 3.**
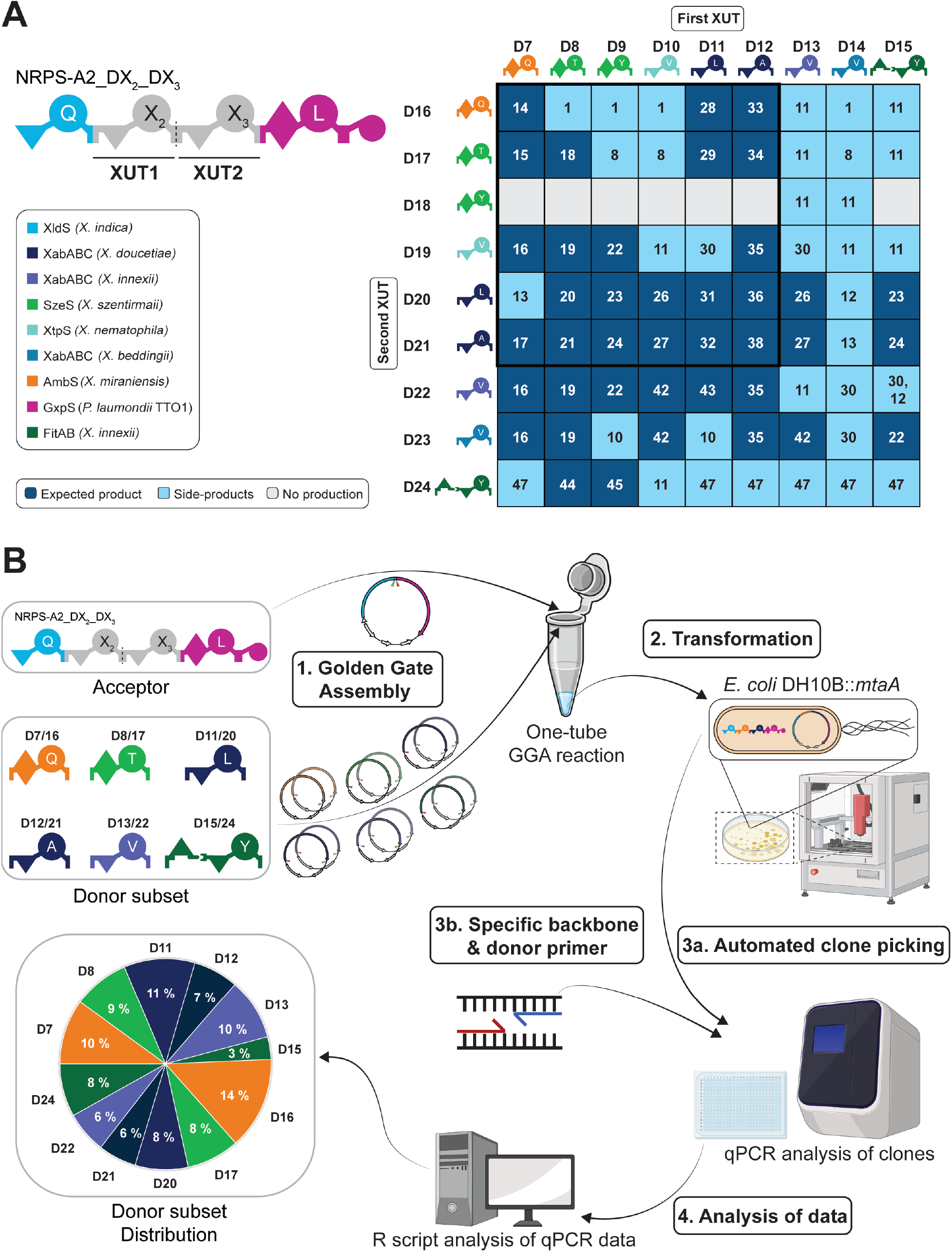
Overview of the lipo tetrapeptide library and the semi-automatic workflow of the qPCR-based NRPS-encoding plasmid validation. **A**) Peptide production as determined by HPLC/MS analysis of the 81 NRPS constructs of the tetra-modular library, where XUT1 and XUT2 have been varied by usage of donor plasmid D7-D15 and D16-D24, respectively. The black box highlights the first set of 36 constructed using D7-D12 and D16-D21. Dark blue, light blue and grey tiles represent the detection of the expected peptides (see Fig. 4), peptide derivatives (see Table S7) and no peptide production, respectively. Numbers within tiles refer to the produced peptides (see Fig. 4 & Table S7). The origin of the different modules is indicated by colour. **B**) Workflow of the semi-automated qPCR-based NRPS plasmid validation for the distribution analysis of the modified donor subset. For NRPS domain explanations, see Fig. 1 (Fig. 3B was created with BioRender).

**Figure 4.**
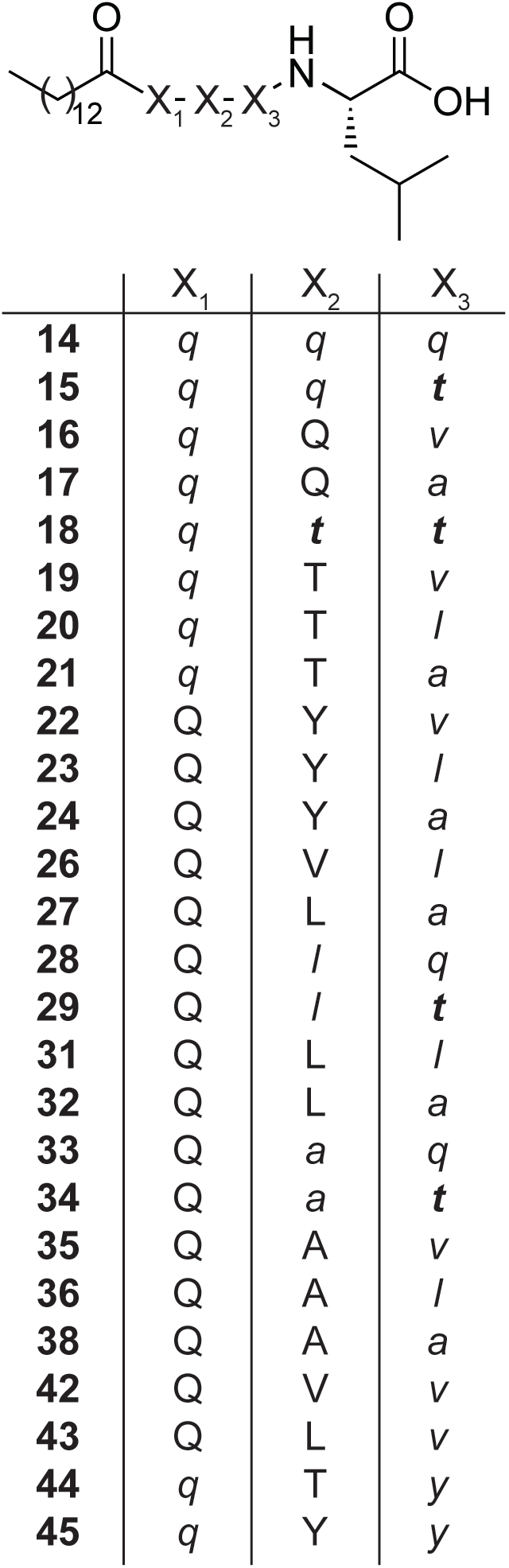
Chemical structures of all identified expected peptides produced by the NRPS systems shown in Figure 3A. The AA “X_1_” (Gln, Q) can have either L- and D-configuration, dependent on the C or C/E domain preceding the A domain of the inserted XUT. X_2_ and X_3_ describe the AAs in the first and second incorporated XUT. D-amino acids are shown in small and italic letters, and D-*allo*-Thr is shown in small, italic and bold letters.

All results regarding the produced peptides are summarised in Table S7, showing each construct with the detected peptides. Generally, all functional NRPS produced the peptides C14-QL and C14-*q*L **11** (Fig. S5-S7), which only contained the building blocks from the acceptor plasmid. Therefore, **11** was only mentioned explicitly in Fig. 3A when no other derivatives were detected. Furthermore, in some NRPS systems, two peaks with the same *m*/*z* ratio were detected, indicating incomplete epimerisation of the C/E domain of the terminal module as a result of the engineered NRPS (Fig. S5 & S7).

Similar to the single-module library, the distribution of the incorporated modules was again tested to confirm equal distribution within a library approach. The new valine (D13 and D22) and tyrosine (D15 and D24) modules were used in place of the lower-performing ones (D9 and D18; D10 and D19) (Fig. 3B). In order to simplify the analysis process independent of full plasmid or possible donor sequencing, an end-point qPCR-based analysis was developed (SI Part B) to investigate the incorporated fragment and its position (see supplementary information D). For this analysis, 288 random clones were selected and tested for their plasmid composition (Fig. 3B). While D16 is slightly overrepresented at 14% and D15 is underrepresented at 3.5%, most of the donors used show a distribution of 8-9%. This shows that this approach can be used to efficiently generate randomised libraries, while specific colonies can be analysed, as shown above.

### Starter module library

Following the success of utilising GGA to incorporate elongation modules, the method should be expanded to starter modules, allowing the attachment of different N-termini to the produced peptide. The starter donor was designed to be analogous to the elongation donors, containing the A overhang to ensure compatibility with the elongation module library. The upstream overhang was designated as the start codon “ATG”. It is also noteworthy that, given the requirement of four bases, the last nucleotide before the start codon (xATG) was utilised to prevent alterations to the subsequent AAs of the expressed NRPS. Consequently, the overhang S (“starter”; CATG) and A were employed to incorporate starter modules from donors, exhibiting a design compatible with the previously described elongation donors. Regarding the new acceptor NRPS-A3, one of the best-producing constructs from the elongation library, designated as NRPS-A2_D6, was used (Fig. 5). The starter module was removed, and BsaI restriction sites were introduced in a manner similar to that used for NRPS-A1 and NRPS-A2. The D25 starter (C14-Q) was then used as the positive control. The additional starter modules originate from the xenoamicine-producing NRPS (XabABC) (D26; C4-P) from *X. doucetiae*^[32]^, the ambactin-producing NRPS (AmbS) (D27; S) from *X. indica* having a non-functional C_starter_ domain^[18]^, and a previously uncharacterised starter from an unknown NRPS-encoding gene cluster from the *X. ishibashii* LC536431 (Fig. 5). The extracts of the four resulting NRPS systems NRPS-A3_D25 to -D28 were analysed using HPLC/MS, showing the production of the expected products (Fig. 5B). Both NRPS-A3_D27 and _D28 produced peptides (**50, 51**) without any acyl group at the N-terminus, suggesting the C_starter_ domain of LC536431 is also non-functional and only responsible for the incorporation of leucine. The success of using starter modules for the GGA approach increases the combinatorial possibilities as well as the ability to modify the N-terminus of peptides.

**Figure 5.**
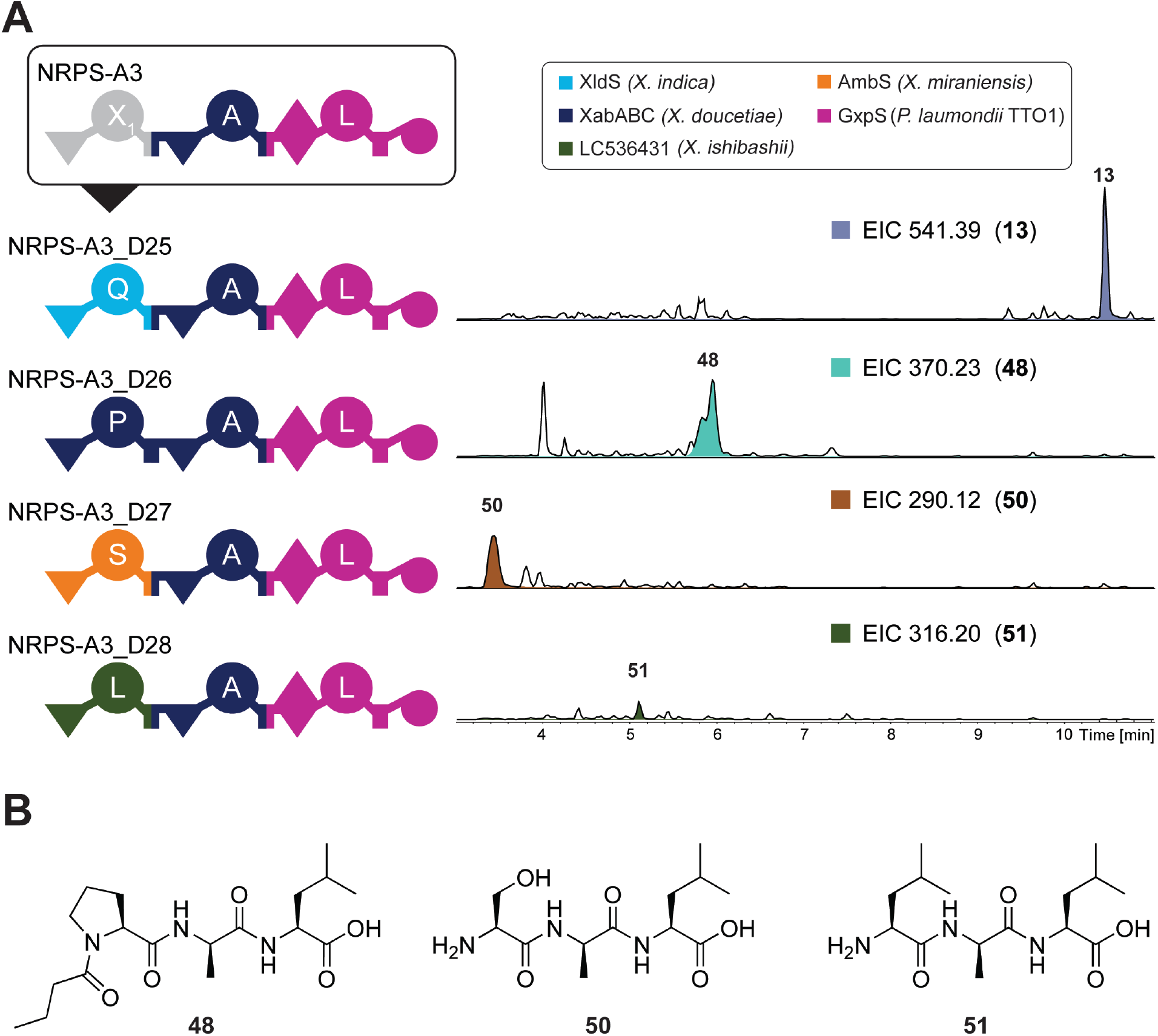
Schematic representation of the starter module system for GGA-based engineered NRPS. **A**) Schematic representation of NRPS-A3 and NRPS-A3_D25 to _D28 with the colour code of the original NRPS. The A domain specificities are indicated by the one-letter AA code. The final plasmids were assembled via GGA using the acceptor plasmid encoding NRPS-A3, lacking a starter module, and the donors D25-D28. HPLC/MS data of compounds **13, 48, 50**, and **51** produced in *E. coli* DH10B::*mtaA* expressing NRPS-A3_D25-D28. Base Peak Chromatogram (BPC, black line) and the Extracted Ion Chromatogram (EIC) of **13** (*m/z* [M+H]^+^ = 541.39), **48** (*m/z* [M+H]^+^ = 370.23), **50** (*m/z* [M+H]^+^ = 290.12) and **51** (*m/z* [M+H]^+^ = 316.20) are shown. **B**) Chemical structures of the compounds **48, 50** and **51**. For NRPS domain explanations, see Fig. 1.

### Construction of a xenoamicin library using the GGA approach

In order to test GGA for the derivatisation of a natural NRPS system, we aimed at the modification of the NRPS XabABC^[32]^ (Fig. 6A), producing xenoamicins A-C (**52**-**54**) as main products in the heterologous host *E. coli* DH10B::*mtaA*. Xenoamicins are 13-amino acid long lipo-depsipeptides with an acylated linear N-terminal and a cyclic C-terminal part, created by an intramolecular cyclisation from the threonine-8 hydroxyl group to the C-terminal carboxyl group of valine-13. The primary objective was to change the AA composition within the ring structure via modification of XabC, which incorporates the last four AAs (Fig. 6A). Additionally, XabD, an aspartate decarboxylase, is encoded downstream of XabC for the required synthesis of β-alanine (β-A). For simplicity reasons, the genes for XabA and XabB were cloned on a separate pCOLA plasmid for co-expression. The acceptor NRPS-A4 contains the first and last module from XabC, lacking the modules for β-alanine and proline, which can be reintroduced as donor building blocks. BsaI restriction sites, similar to NRPS-A1 and NRPS-A2, were added, creating A/B overhangs. In addition to the modified NRPS, the aspartate decarboxylase XabD was cloned on the acceptor plasmid to provide β-alanine.

**Figure 6.**
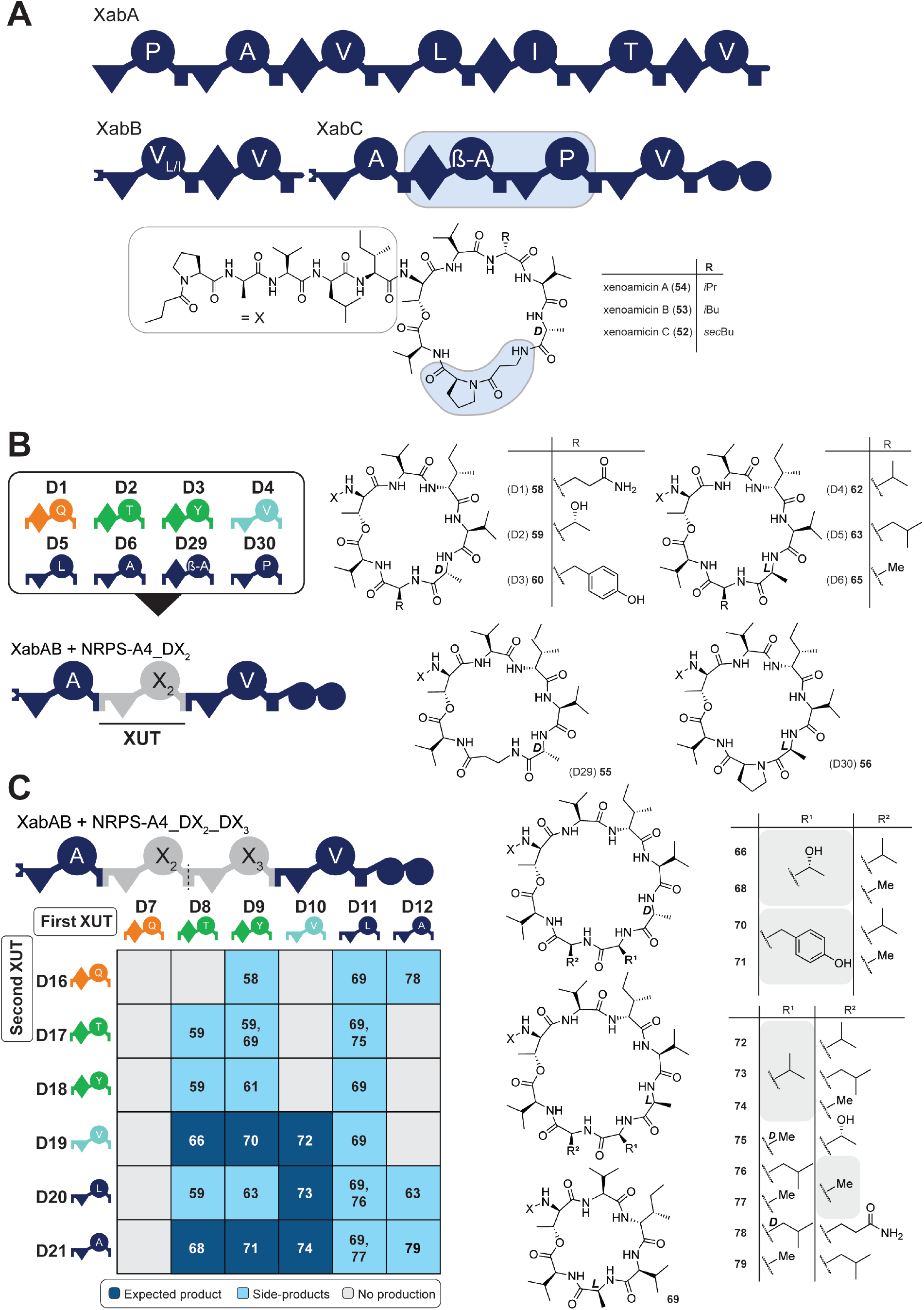
Overview of the xenoamicin library results using replacements of one or two XUTs. **A**) Overview of the XabABC NRPS architecture and the chemical structure of its main products, xenoamicin A-C (**52**-**54**). The exchanged modules and AAs are marked in light blue within the NPRS and the chemical structure, respectively. **B**) Incorporation of single modules from the first donor subset (D1-D6), D29 and D30, all resulting in the production of the expected peptide. The linear N-terminal region is abbreviated as “X”. **C**) Summary of the HPLC/MS analysis results from the extracts of 36 NRPS constructs of the xenoamicin library containing two exchanged XUTs. Dark blue, light blue and grey tiles represent the detection of expected peptides, peptide derivatives, and no peptide production. Numbers within tiles refer to the compounds (Table S9). For the explanation of the NRPS domain origin of the inserted XUTs according to their colour, see Fig. 3. For NRPS domain explanations, see Fig. 1.

In the first experiments, donors D1-D6 (Fig. 2A), as well as the β-alanine (D29) and proline donor (D30), were incorporated into NRPS-A4_DX. The HPLC/MS analysis of the obtained culture extracts shows that all eight NRPS are functional, producing the expected peptides (**55**-**65**) (Fig. 6B & Table S9). Since the first XUT might carry a C/E or a C domain, the configuration of Ala at position 10 might be D (as in the original xenoamicins) or L, respectively.

Furthermore, the native NRPS produces the two derivatives with the same *m*/*z*, xenoamicin C (**52**) and xenoamicin B (**53**), having an identical mass but differ in the incorporation of isoleucine or leucine at position eight (Fig. S9). From previous work^[32]^, we postulate that the peak eluting later is the isoleucine-containing main derivative xenoamicin C (**52** Fig. S9); therefore, Ile-8 containing structures are presented for simplicity (Table S9).

In addition to all constructs being functional, the ring size of the depsipeptide was reduced since two modules from the original NRPS were exchanged against one donor module. Additionally, along with the expected depsipeptides, linear peptides were also detected (Fig. S10).

When two modules using D7-D12 and D16-D21 were incorporated together, peptides with the original number of AAs should be obtained. As a positive control, the parent NRPS-A4_D31_D34, using D31 and D34, was generated first, showing comparable production titers to those of the native XabABC under these conditions (Fig. S9). From the 36 generated *E. coli* strains, eight showed the expected peptide structures, while 16 showed skipping of one or both of the newly incorporated modules, respectively, and 13 did not show any peptide production. Of the overall 23 peptides produced, 17 represent new derivatives of xenoamicin, highlighting the power of the GGA for NRPS engineering. However, for the incorporation of multiple modules, their compatibility might play a more important role than in the insertion of single modules (Fig. 6C).

## Conclusion

NRPS are among the largest enzymes known, and their size and modularity are both a difficulty and a chance for manipulation in order to obtain new-to-nature peptides^[2,33,34]^. While the size of the proteins and their associate gene sizes (∼3 kb per module) can be addressed via the introduction of natural^[35,36]^ or artificial docking domains^[20,37,38]^, which enables simplified cloning of smaller genes on suitable expression plasmids, the recently described XUT approach allows an efficient exchange, insertion or deletion of NRPS parts within a single given NRPS protein^[14]^.

In this work, we could successfully establish a new GGA workflow to leverage the cloning of artificial hybrid NRPS genes via XUT engineering. Compared to the previously applied Gibson assembly, the new approach allows the rapid assembly and verification of defined artificial NRPS, as well as entire defined and randomised artificial NRPS libraries (Figs. 2 and 3). Once generated, donor plasmids can be used for various XUT-based engineering applications like variation of single or multiple sites within non-ribosomal peptides, as demonstrated for the xenoamicin in this work (Fig. 6). With respect to the overall outcome, our approach is similar to protein engineering campaigns^[39]^ where individual AA positions can easily be exchanged or randomised based on DNA-sequence variation. However, the large variability of available NRPS-”encoded” AA building blocks^[2,40,41]^ offers access to a much greater chemical space than the 20 proteinogenic AAs. The flexible application of the GGA-based NRPS engineering comes with greater sustainability in terms of workload and consumables and has the potential to derivatise multiple non-ribosomal peptides in high-throughput to identify novel bioactive peptides.

The increased throughput of the presented workflow also allows the acquisition of larger datasets that contain different NRPS hybrids from various genera of bacteria with functional and non-functional NRPS assemblies. Analysis by machine learning algorithms might result in the identification of rules for acceptor-donor or donor-donor functionality and/or peptide production success, which could be dependent on so far unknown interactions of domains, linkers or protein interfaces.

In conclusion, the established GGA-based cloning for XUT engineering represents a significant improvement in developing and applying NRPS engineering in high-throughput.

## Author Contributions

A.P. performed all experiments except for the qPCR-based NRPS plasmid validation experiments. The development of the qPCR method was done by T.A.L., D.S. and A.P. The qPCR related experiments were done by A.P. and T.A.L. The custom R script was written by T.A.L. H.B.B. supervised the work and the paper was written by A.P. with contributions from all authors.

## Acknowledgements

Work in the Bode lab was supported by an ERC Advanced Grant (835108) and the Max Planck Society. The work in the Schindler lab is supported by the Max Planck Society in the framework of the MaxGENESYS project. This work was supported by the European Union (NextGenerationEU) via the European Regional Development Fund (ERDF) by the state Hesse of Germany within the project “Biotechnological production of reactive peptides from waste streams as lead structures for drug development” (D.S. & H.B.B.). The authors are grateful to Kenan Bozhüyük and all former and present group members of the NRPS engineering team for fruitful discussions.

## Data availability statement

All data and materials can be found within the manuscript, supporting information or can be requested from the corresponding author.

