## Supplementary data, tables and figures for "High-throughput engineering and modification of non-ribosomal peptide synthetases based on Golden Gate assembly"

### A. Materials and Methods

#### 1. Cultivation of strains

All *E. coli* DH10B::*mtaA* were cultured on liquid or solid low-salt LB medium (pH 7.5, 10 g/L tryptone, 5 g/L yeast extract and 5 g/L NaCl). As a selection marker, either chloramphenicol (34 µg/mL), kanamycin (50 µg/mL) or gentamicin (20 µg/mL) was added. Solid medium was prepared with 1% agar (w/v). The cells were cultivated at 37 °C. All *Xenorhabdus* strains were cultivated on solid low-salt LB medium (pH 7.5, 10 g/L tryptone, 5 g/L yeast extract and 5 g/L NaCl, 1% agar (w/v)). The cells were cultivated at 28 °C.

#### 2. Cloning of biosynthetic gene clusters and NRPS acceptors and donors

Genomic DNA (gDNA) was isolated via Monarch® Genomic DNA Purification Kit (NEB) from the respective strains depicted in Table S1 for usage as a template for PCR amplification. Alternatively, plasmid DNA was used as a template, especially for the re-amplification of donor plasmid fragments with different overhangs. In this work, Q5® High-Fidelity DNA Polymerase (NEB) or Phusion® Hot Start Flex DNA Polymerase (NEB) was utilised to create all DNA fragments. Primer pair sequences and products are summarised in Table S4. The PCR amplified fragments were digested with DpnI (NEB) (manufacturer instructions) and purified by gel extraction from 1% (w/v) agarose gels using the Monarch® DNA Gel Extraction Kit (NEB).

The acceptor and donor plasmids were cloned using Gibson assembly from the NEBuilder® HiFi DNA Assembly Cloning Kit (NEB). Additionally, native BsaI restriction sites are removed by point mutagenesis. The final constructs were assembled using Golden Gate Assembly from NEBridge® Golden Gate Assembly Kit (NEB) (reaction according to manufacturer's instructions) with the acceptor and donor plasmids, using the protocol for multiple fragments ((5 min 37 °C → 5 min 16 °C) x 30 cycles followed by 5 min 60 °C). Gibson and Golden Gate assembled plasmids were transformed by heat shock into chemical-competent *E. coli* DH10B::*mtaA*. Plasmids were isolated using the Monarch® Plasmid Miniprep Kit (NEB).

### 3. Heterologous expression of NRPS constructs

After plasmid transformation via heat shock into chemical competent *E. coli* DH10B::*mtaA*, cells were cultivated overnight in LB medium containing the necessary antibiotics. 24-deep well plates with 4 mL XPP medium[1] (each well) containing the required antibiotics and 0.02% L-arabinose (w/v) were inoculated with 1% overnight grown culture. The cells were cultivated at 22 °C for 72 hours at 200 rpm.

### 4. Culture extraction and HPLC/MS analysis

The cultures were extracted by mixing the culture in a 1:1 ratio with methanol, followed by a 1 hour incubation at room temperature while shaking. The deep-well plates were centrifuged for 30 min at 22 °C and 4400 x g. Extracts in reaction tubes were centrifuged for 20 min at 22 °C and 17000 x g.

Cleared supernatant was further used for high-performance liquid chromatography mass spectrometry (HPLC/MS) and high-performance liquid chromatography high-resolution mass spectrometry (HPLC/HRMS) analysis. For HPLC/MS, the analysis was performed using Agilent Infinity II coupled to an AmaZon speed ETD spectrometer (Bruker) with an ACQUITY UPLC BEH C18 column (130 Å, 2.1 mm × 100 mm, 1.7-µm particle size, Waters) at a flow rate of 0.4 mL/min (5–95% acetonitrile/water with 0.1% formic acid, vol/vol, 16 min, and an electrospray ionisation (ESI) source set to positive ionisation mode. To obtain high-resolution masses, HPLC/HRMS analysis was performed using performed using Bruker Elute coupled to an timsTOF fleX spectrometer (Bruker) with an ACQUITY UPLC BEH C18 column (130 Å, 2.1 mm × 100 mm, 1.7-µm particle size, Waters) at a flow rate of 0.4 mL/min (5–95% acetonitrile/water with 0.1% formic acid, vol/vol, 16 min, and a VIP-HESI (ESI) source set to positive ionisation mode. The MS data was analysed using DataAnalysis 6.1 (Bruker). Additionally, the peptide masses and sequences were calculated using Pep-Calc API[2].

### 5. Golden Gate overhangs

The described overhangs in this work are all the 16 possible overhangs for this fusion site using the codons of both glycines (Fig. S1). However, not all of them show perfect compatibility with each other, leading to potential mismatches during the GGA. Systems such as NEBridge Ligase Fidelity Viewer[3–5] offer excellent analysis capabilities for viewing fidelities and identifying possible mismatches, allowing for adjustments to the used overhangs accordingly.

### 6. qPCR analysis for NRPS constructs

Candidates were picked from a 150-mm 2% LB agar Petri dish using the Singer PIXL colony picker onto a 96 SBS-format agar plate. Following an overnight growth period at 37 °C, the colonies were transferred to 50 µL of sterile water. qPCR primers were designed using the Primer3 software[6]. Subsequently, 25 nL of 10 µM premixed primers and 100 nL of the resuspended colonies were transferred to 384 qPCR plates (Sarstedt, 72.1984.202) using the Echo525 acoustic liquid dispenser (Labcyte), and analysis was conducted on the same day. 1 µL of 1X Luna Universal qPCR Master Mix (NEB) was dispensed using a nanoliter dispenser (Cobra, Art Robbins Instruments). Subsequently, the plates were subjected to a brief centrifugation step before being sealed with an optical clear permanent seal (Agilent, 24212-001) using the Plate Loc Thermal Microplate Sealer (Agilent) at 180 °C for 1 s. The Applied Biosystems QuantStudio 5 was then utilised for SYBR Green detection of the prepared plates. Samples were first subjected to a preincubation step at 50 °C (1.6 °C/s) for 2 minutes, followed by a 1-minute incubation at 95 °C (2.57 °C/s). Subsequently, a PCR reaction was performed, comprising 30 cycles each at 95 °C (2.57 °C/s) for 1 s and 67 °C (2 °C/s) for 1 min, with a single acquisition. A melting curve was acquired from 97 °C (0.1 °C/s) with continuous acquisition. The resulting data was exported as an Excel file and subsequently analysed using a custom R script (see **A.7**).

### 7. Custom R script for the qPCR analysis

Found in a separate SI

### B. Supplementary method and results: Results of the end-point qPCR-based NRPS plasmid validation used for the tetra-modular NRPS library

During the validation of the libraries, Sanger sequencing confirmed the incorporation of one module during GGA but is not suited for sequencing two or more modules. This is due to the fact that NRPS XUTs exceed the size of traditional Sanger sequencing reads (~800 bp) and would, therefore, require multiple repetitions. Alternatively, Nanopore sequencing enables whole plasmid sequencing based on its long-read technology.

However, these methods require the cultivation of each strain of the library, as well as the extraction and purification of DNA (i.e. the plasmid). Since known DNA sequences were used as a predefined set of donors, we developed an end-point quantitative polymerase chain reaction (qPCR) method[7] to validate the GGA-generated NRPS libraries, which utilises *E. coli* colonies as templates and eliminates the need for extensive liquid cultivation, DNA extraction, and purification. Furthermore, in comparison to regular PCR, qPCR eliminates the need for gel electrophoresis and subsequently allows for the minimisation of reactions to 1 µL, saving materials.

For qPCR, two specific primers were designed for each donor, one for the starter and one for the termination module on the acceptor plasmid (Fig. S8 and Table S5). The forward primers are located at the end of the A domain, whereas the reverse primers are located within the T-C linker or at the beginning of the C domain. This results in multiple primer combinations for each possible XUT position within the GG-assembled NRPS constructs. For the detection of the correct XUT and its position, each primer combination was used for each template. Optimally, only two qPCR reactions show a positive result, creating the expected amplicon (~450-600 bp). The primers were first tested on purified plasmid DNA to record the melting temperature ($T_m$) and the size of each amplicon. Following successful results, the system was applied to *E. coli* cells (see **A.6**), where qPCR amplifications can be easily traced back to the original colony.

The tetra-modular library was tested by examining 288 colonies (Fig. 3B) (287 of which contained the library and one of which included an empty plasmid). The $C_t$ cut-off was placed at 18 cycles, and the minimum $T_m$ at 80 °C. This analysis determined that five of the colonies were false negatives, as they failed to produce two signals. Furthermore, nine showed signals for two separate plasmids. Re-evaluation of the original colony agar plates revealed that during colony-picking, double colonies were selected and therefore picked. Of the examined colonies, 96 were cultivated, the plasmids purified and sent for plasmid sequencing. The sequencing results were consistent with those from the qPCR, including the nine candidates that showed signals for two plasmids, confirming two plasmids were present. With false negatives and the double-picked colonies, the success rate of this method was over 95%. This showed great efficiency in utilising minimal resources and huge time benefits, while maintaining a good success rate for the analysis of the GG-assembled NRPS constructs.

### C. Supplemental Tables

**Table S1.** Strains used in this work.

| Strain | Genotype/NRPS | Reference |
|---|---|---|
| *E. coli* DH10B::mtaA | F_mcrA (*mrr-hsd*RMS- *mcr*BC), 80*lac*ZΔ, M15, Δ*lac*X74 *rec*A1 *end*A1 *ara*D 139Δ(*ara*, *leu*)7697 *gal*U *gal*Kλ *rps*L (*Strr*) *nup*G and mtaA from pCK_*mtaA*Δ*entD* / - | [8] |
| *P. laumondii* TTO1 | WT (*gxpS*[9]) | DSMZ |
| *X. indica* DSM 17382 | WT (*xldS*[8]) | DSMZ |
| *X. miraniensis* DSM 17902 | WT (ambS[8]) | DSMZ |
| *X. szentirmaii* DSM 16338 | WT (*szeS*[10]) | DSMZ |
| *X. nematophila* ATCC 19061 | WT (*xtpS*[11]) | DSMZ |
| *X. doucetiae* DSM 17909 | WT (*xabABCD*[12]) | DSMZ |
| *X. innexi* DSM 16336 | WT (*xabABC*, *fitAB*[13]) | DSMZ |
| *X. beddingii* DSM 4764 | WT (*xabABC*) | DSMZ |
| *X. ishibashii* DSM 22670 | WT (Locus tag: LC536431) | DSMZ |

**Table S2.** NRPS, corresponding plasmids and genotypes used in this work.

| NRPS- | Plasmids | Genotype | Reference |
|---|---|---|---|
| | pCOLA_ara/tacI | ori ColA, $kan^R$, *araC*-$P_{BAD}$ and *tacI* | [14] |
| | pACYC_ara/araE _tacI | ori p15A, $cm^R$, *araC*-$P_{BAD}$, *tacI* and *araE* | [15] |
| | pSEVA681 | ori pUC, oriT, $gm^R$ | [16,17] |
| | pAP128 | ori ColA, $kan^R$, *araC*-$P_{BAD}$ *xabAB* and *tacI* | This work |
| **A1** | pAP38 | ori p15A, $cm^R$, *araC*-$P_{BAD}$ *xldS*_C1A1T$1_{1/2}$_2xBsaI_*szeS*_T$5_{1/2}$C6A6 T6Te *tacI* and *araE* | This work |
| **A2** | pAP37 | ori p15A, $cm^R$, *araC*-$P_{BAD}$ *xldS*_C1A1T$1_{1/2}$_2xBsaI_*gxpS*_T$4_{1/2}$C/E5 A5T5Te *tacI* and *araE* | This work |

| NRPS- | Plasmids | Genotype | Reference |
|---|---|---|---|
| **A3** | pAP106 | ori p15A, $cm^R$, *araC-$P_{BAD}$* 2xBsaI_*xabA*_T1$_{1/2}$AC2A2T2$_{1/2}$_*gxpS*_T4$_{1/2}$C/E5A5T5Te *tacI* and *araE* | This work |
| **A4** | pAP119 | ori p15A, $cm^R$, *araC-$P_{BAD}$* *xabC*_C1A1T1$_{1/2}$_2xBsaI_*xabC*_T3$_{1/2}$C4A 4T4TeTe *tacI* and *araE* | This work |
| **1** | pAP32 | ori p15A, $cm^R$, *araC-$P_{BAD}$* *xldS*_C1A1T1$_{1/2}$_*szeS*_T1$_{1/2}$C/E2A2T2$_{1/2}$_ *szeS*_T5$_{1/2}$C6A6T6Te *tacI* and *araE* | This work |
| **2** | pAP33 | ori p15A, $cm^R$, *araC-$P_{BAD}$* *xldS*_C1A1T1$_{1/2}$_*ambS*_T1$_{1/2}$C/E2A2T2$_{1/2}$_ *szeS*_T5$_{1/2}$C6A6T6Te *tacI* and *araE* | This work |
| **3** | pAP34 | ori p15A, $cm^R$, *araC-$P_{BAD}$* *xldS*_C1A1T1$_{1/2}$_*szeS*_T1$_{1/2}$C/E2A2T2$_{1/2}$_ *gxpS*_T4$_{1/2}$C/E5A5T5Te *tacI* and *araE* | This work |
| **4** | pAP35 | ori p15A, $cm^R$, *araC-$P_{BAD}$* *xldS*_C1A1T1$_{1/2}$_*ambS*_T1$_{1/2}$C/E2A2T2$_{1/2}$_ *gxpS*_T4$_{1/2}$C/E5A5T5Te *tacI* and *araE* | This work |
| **5** | pAP36 | ori p15A, $cm^R$, *araC-$P_{BAD}$* *xldS*_C1A1T1$_{1/2}$_*szeS*_T4$_{1/2}$C/E5A5T5$_{1/2}$_ *szeS*_T5$_{1/2}$C6A6T6Te *tacI* and *araE* | This work |
| **5_mod** | pAP36b | ori p15A, $cm^R$, *araC-$P_{BAD}$* *xldS*_C1A1T1$_{1/2}$_*szeS*_T4$_{1/2}$C/E5A5T5$_{1/2}$_ *szeS*_T5$_{1/2}$C6A6T6Te, G2093G (GGA>GGT), G2094G (GGA>GGT) *tacI* and *araE* | This work |
| **6** | pAP111 | ori p15A, $cm^R$, *araC-$P_{BAD}$* *xabCD tacI* and *araE* | This work |
| **7** | pAP112 | ori p15A, $cm^R$, *araC-$P_{BAD}$* *xabCD*ΔBsaI *tacI* and *araE* | This work |
| **A2_D1** | pAP37_39 | ori p15A, $cm^R$, *araC-$P_{BAD}$* *xldS*_C1A1T1$_{1/2}$_*ambS*_T1$_{1/2}$C/E2A2T2$_{1/2}$_ *gxpS*_T4$_{1/2}$C/E5A5T5Te *tacI* and *araE* | This work |
| **A2_D2** | pAP37_40 | ori p15A, $cm^R$, *araC-$P_{BAD}$* *xldS*_C1A1T1$_{1/2}$_*szeS*_T1$_{1/2}$C/E2A2T2$_{1/2}$_ *gxpS*_T4$_{1/2}$C/E5A5T5Te *tacI* and *araE* | This work |
| **A2_D3** | pAP37_41 | ori p15A, $cm^R$, *araC-$P_{BAD}$* *xldS*_C1A1T1$_{1/2}$_*szeS*_T4$_{1/2}$C/E5A5T5$_{1/2}$_ *gxpS*_T4$_{1/2}$C/E5A5T5Te *tacI* and *araE* | This work |
| **A2_D4** | pAP37_42 | ori p15A, $cm^R$, *araC-$P_{BAD}$* *xldS*_C1A1T1$_{1/2}$_*xtpS*_T2$_{1/2}$C3A3T3$_{1/2}$_*gxp S*_T4$_{1/2}$C/E5A5T5Te *tacI* and *araE* | This work |
| **A2_D5** | pAP37_43 | ori p15A, $cm^R$, *araC-$P_{BAD}$* *xldS*_C1A1T1$_{1/2}$_*xabA*_T3$_{1/2}$C4A4T4$_{1/2}$_*gx pS*_T4$_{1/2}$C/E5A5T5Te *tacI* and *araE* | This work |

| NRPS- | Plasmids | Genotype | Reference |
|---|---|---|---|
| **A2_D6** | pAP37_44 | ori p15A, $cm^R$, *araC-P*$_{BAD}$ *xldS*_C1A1T1$_{1/2}$_*xabA*_T1$_{1/2}$C2A2T2$_{1/2}$_*gxpS*_T4$_{1/2}$C/E5A5T5Te *tacI* and *araE* | This work |
| **A1_D1** | pAP38_39 | ori p15A, $cm^R$, *araC-P*$_{BAD}$ *xldS*_C1A1T1$_{1/2}$_*ambS*_T1$_{1/2}$C/E2A2T2$_{1/2}$_*szeS*_T5$_{1/2}$C6A6T6Te *tacI* and *araE* | This work |
| **A1_D2** | pAP38_40 | ori p15A, $cm^R$, *araC-P*$_{BAD}$ *xldS*_C1A1T1$_{1/2}$_*szeS*_T1$_{1/2}$C/E2A2T2$_{1/2}$_*szeS*_T5$_{1/2}$C6A6T6Te *tacI* and *araE* | This work |
| **A1_D3** | pAP38_41 | ori p15A, $cm^R$, *araC-P*$_{BAD}$ *xldS*_C1A1T1$_{1/2}$_*szeS*_T4$_{1/2}$C/E5A5T5$_{1/2}$_*szeS*_T5$_{1/2}$C6A6T6Te *tacI* and *araE* | This work |
| **A1_D4** | pAP38_42 | ori p15A, $cm^R$, *araC-P*$_{BAD}$ *xldS*_C1A1T1$_{1/2}$_*xtpS*_T2$_{1/2}$C3A3T3$_{1/2}$_*szeS*_T5$_{1/2}$C6A6T6Te *tacI* and *araE* | This work |
| **A1_D5** | pAP38_43 | ori p15A, $cm^R$, *araC-P*$_{BAD}$ *xldS*_C1A1T1$_{1/2}$_*xabA*_T3$_{1/2}$C4A4T4$_{1/2}$_*szeS*_T5$_{1/2}$C6A6T6Te *tacI* and *araE* | This work |
| **A1_D6** | pAP38_44 | ori p15A, $cm^R$, *araC-P*$_{BAD}$ *xldS*_C1A1T1$_{1/2}$_*xabA*_T1$_{1/2}$C2A2T2$_{1/2}$_*szeS*_T5$_{1/2}$C6A6T6Te *tacI* and *araE* | This work |
| **A2_D7_D16** | pAP37_45_51 | ori p15A, $cm^R$, *araC-P*$_{BAD}$ *xldS*_C1A1T1$_{1/2}$_*ambS*_T1$_{1/2}$C/E2A2T2$_{1/2}$_*ambS*_T1$_{1/2}$C/E2A2T2$_{1/2}$_*gxpS*_T4$_{1/2}$C/E5A5T5Te *tacI* and *araE* | This work |
| **A2_D7_D17** | pAP37_45_52 | ori p15A, $cm^R$, *araC-P*$_{BAD}$ *xldS*_C1A1T1$_{1/2}$_*ambS*_T1$_{1/2}$C/E2A2T2$_{1/2}$_*szeS*_T1$_{1/2}$C/E2A2T2$_{1/2}$_*gxpS*_T4$_{1/2}$C/E5A5T5Te *tacI* and *araE* | This work |
| **A2_D7_D18** | pAP37_45_53 | ori p15A, $cm^R$, *araC-P*$_{BAD}$ *xldS*_C1A1T1$_{1/2}$_*ambS*_T1$_{1/2}$C/E2A2T2$_{1/2}$_*szeS*_T4$_{1/2}$C/E5A5T5$_{1/2}$_*gxpS*_T4$_{1/2}$C/E5A5T5Te *tacI* and *araE* | This work |
| **A2_D7_D19** | pAP37_45_54 | ori p15A, $cm^R$, *araC-P*$_{BAD}$ *xldS*_C1A1T1$_{1/2}$_*ambS*_T1$_{1/2}$C/E2A2T2$_{1/2}$_*xtpS*_T2$_{1/2}$C3A3T3$_{1/2}$_*gxpS*_T4$_{1/2}$C/E5A5T5Te *tacI* and *araE* | This work |
| **A2_D7_D20** | pAP37_45_55 | ori p15A, $cm^R$, *araC-P*$_{BAD}$ *xldS*_C1A1T1$_{1/2}$_*ambS*_T1$_{1/2}$C/E2A2T2$_{1/2}$_*xabA*_T3$_{1/2}$C4A4T4$_{1/2}$_*gxpS*_T4$_{1/2}$C/E5A5T5Te *tacI* and *araE* | This work |
| **A2_D7_D21** | pAP37_45_56 | ori p15A, $cm^R$, *araC-P*$_{BAD}$ *xldS*_C1A1T1$_{1/2}$_*ambS*_T1$_{1/2}$C/E2A2T2$_{1/2}$_*xabA*_T1$_{1/2}$C2A2T2$_{1/2}$_*gxpS*_T4$_{1/2}$C/E5A5T5Te *tacI* and *araE* | This work |

| NRPS- | Plasmids | Genotype | Reference |
|---|---|---|---|
| **A2_D8_D16** | pAP37_46_51 | ori p15A, *cm*$^R$, *araC-P*$_{BAD}$ *xldS*_C1A1T1$_{1/2}$_*szeS*_T1$_{1/2}$C/E2A2T2$_{1/2}$_*ambS*_T1$_{1/2}$C/E2A2T2$_{1/2}$_*gxpS*_T4$_{1/2}$C/E5A5T5Te *tacI* and *araE* | This work |
| **A2_D8_D17** | pAP37_46_52 | ori p15A, *cm*$^R$, *araC-P*$_{BAD}$ *xldS*_C1A1T1$_{1/2}$_*szeS*_T1$_{1/2}$C/E2A2T2$_{1/2}$_*szeS*_T1$_{1/2}$C/E2A2T2$_{1/2}$_*gxpS*_T4$_{1/2}$C/E5A5T5Te *tacI* and *araE* | This work |
| **A2_D8_D18** | pAP37_46_53 | ori p15A, *cm*$^R$, *araC-P*$_{BAD}$ *xldS*_C1A1T1$_{1/2}$_*szeS*_T1$_{1/2}$C/E2A2T2$_{1/2}$_*szeS*_T4$_{1/2}$C/E5A5T5$_{1/2}$_*gxpS*_T4$_{1/2}$C/E5A5T5Te *tacI* and *araE* | This work |
| **A2_D8_D19** | pAP37_46_54 | ori p15A, *cm*$^R$, *araC-P*$_{BAD}$ *xldS*_C1A1T1$_{1/2}$_*szeS*_T1$_{1/2}$C/E2A2T2$_{1/2}$_*xtpS*_T2$_{1/2}$C3A3T3$_{1/2}$_*gxpS*_T4$_{1/2}$C/E5A5T5Te *tacI* and *araE* | This work |
| **A2_D8_D20** | pAP37_46_55 | ori p15A, *cm*$^R$, *araC-P*$_{BAD}$ *xldS*_C1A1T1$_{1/2}$_*szeS*_T1$_{1/2}$C/E2A2T2$_{1/2}$_*xabA*_T3$_{1/2}$C4A4T4$_{1/2}$_*gxpS*_T4$_{1/2}$C/E5A5T5Te *tacI* and *araE* | This work |
| **A2_D8_D21** | pAP37_46_56 | ori p15A, *cm*$^R$, *araC-P*$_{BAD}$ *xldS*_C1A1T1$_{1/2}$_*szeS*_T1$_{1/2}$C/E2A2T2$_{1/2}$_*xabA*_T1$_{1/2}$C2A2T2$_{1/2}$_*gxpS*_T4$_{1/2}$C/E5A5T5Te *tacI* and *araE* | This work |
| **A2_D9_D16** | pAP37_47_51 | ori p15A, *cm*$^R$, *araC-P*$_{BAD}$ *xldS*_C1A1T1$_{1/2}$_*szeS*_T4$_{1/2}$C/E5A5T5$_{1/2}$_*ambS*_T1$_{1/2}$C/E2A2T2$_{1/2}$_*gxpS*_T4$_{1/2}$C/E5A5T5Te *tacI* and *araE* | This work |
| **A2_D9_D17** | pAP37_47_52 | ori p15A, *cm*$^R$, *araC-P*$_{BAD}$ *xldS*_C1A1T1$_{1/2}$_*szeS*_T4$_{1/2}$C/E5A5T5$_{1/2}$_*szeS*_T1$_{1/2}$C/E2A2T2$_{1/2}$_*gxpS*_T4$_{1/2}$C/E5A5T5Te *tacI* and *araE* | This work |
| **A2_D9_D18** | pAP37_47_53 | ori p15A, *cm*$^R$, *araC-P*$_{BAD}$ *xldS*_C1A1T1$_{1/2}$_*szeS*_T4$_{1/2}$C/E5A5T5$_{1/2}$_*szeS*_T4$_{1/2}$C/E5A5T5$_{1/2}$_*gxpS*_T4$_{1/2}$C/E5A5T5Te *tacI* and *araE* | This work |
| **A2_D9_D19** | pAP37_47_54 | ori p15A, *cm*$^R$, *araC-P*$_{BAD}$ *xldS*_C1A1T1$_{1/2}$_*szeS*_T4$_{1/2}$C/E5A5T5$_{1/2}$_*xtpS*_T2$_{1/2}$C3A3T3$_{1/2}$_*gxpS*_T4$_{1/2}$C/E5A5T5Te *tacI* and *araE* | This work |
| **A2_D9_D20** | pAP37_47_55 | ori p15A, *cm*$^R$, *araC-P*$_{BAD}$ *xldS*_C1A1T1$_{1/2}$_*szeS*_T4$_{1/2}$C/E5A5T5$_{1/2}$_*xabA*_T3$_{1/2}$C4A4T4$_{1/2}$_*gxpS*_T4$_{1/2}$C/E5A5T5Te *tacI* and *araE* | This work |

| NRPS- | Plasmids | Genotype | Reference |
|---|---|---|---|
| **A2_D9_D21** | pAP37_47_56 | ori p15A, *cm*$^R$, *araC-P*$_{BAD}$ *xldS*_C1A1T1$_{1/2}$_*szeS*_T4$_{1/2}$C/E5A5T5$_{1/2}$_*xabA*_T1$_{1/2}$C2A2T2$_{1/2}$_*gxpS*_T4$_{1/2}$C/E5A5T5Te *tacI* and *araE* | This work |
| **A2_D10_D16** | pAP37_48_51 | ori p15A, *cm*$^R$, *araC-P*$_{BAD}$ *xldS*_C1A1T1$_{1/2}$_*xtpS*_T2$_{1/2}$C3A3T3$_{1/2}$_*ambS*_T1$_{1/2}$C/E2A2T2$_{1/2}$_*gxpS*_T4$_{1/2}$C/E5A5T5Te *tacI* and *araE* | This work |
| **A2_D10_D17** | pAP37_48_52 | ori p15A, *cm*$^R$, *araC-P*$_{BAD}$ *xldS*_C1A1T1$_{1/2}$_*xtpS*_T2$_{1/2}$C3A3T3$_{1/2}$_*szeS*_T1$_{1/2}$C/E2A2T2$_{1/2}$_*gxpS*_T4$_{1/2}$C/E5A5T5Te *tacI* and *araE* | This work |
| **A2_D10_D18** | pAP37_48_53 | ori p15A, *cm*$^R$, *araC-P*$_{BAD}$ *xldS*_C1A1T1$_{1/2}$_*xtpS*_T2$_{1/2}$C3A3T3$_{1/2}$_*szeS*_T4$_{1/2}$C/E5A5T5$_{1/2}$_*gxpS*_T4$_{1/2}$C/E5A5T5Te *tacI* and *araE* | This work |
| **A2_D10_D19** | pAP37_48_54 | ori p15A, *cm*$^R$, *araC-P*$_{BAD}$ *xldS*_C1A1T1$_{1/2}$_*xtpS*_T2$_{1/2}$C3A3T3$_{1/2}$_*xtpS*_T2$_{1/2}$C3A3T3$_{1/2}$_*gxpS*_T4$_{1/2}$C/E5A5T5Te *tacI* and *araE* | This work |
| **A2_D10_D20** | pAP37_48_55 | ori p15A, *cm*$^R$, *araC-P*$_{BAD}$ *xldS*_C1A1T1$_{1/2}$_*xtpS*_T2$_{1/2}$C3A3T3$_{1/2}$_*xabA*_T3$_{1/2}$C4A4T4$_{1/2}$_*gxpS*_T4$_{1/2}$C/E5A5T5Te *tacI* and *araE* | This work |
| **A2_D10_D21** | pAP37_48_56 | ori p15A, *cm*$^R$, *araC-P*$_{BAD}$ *xldS*_C1A1T1$_{1/2}$_*xtpS*_T2$_{1/2}$C3A3T3$_{1/2}$_*xabA*_T1$_{1/2}$C2A2T2$_{1/2}$_*gxpS*_T4$_{1/2}$C/E5A5T5Te *tacI* and *araE* | This work |
| **A2_D11_D16** | pAP37_49_51 | ori p15A, *cm*$^R$, *araC-P*$_{BAD}$ *xldS*_C1A1T1$_{1/2}$_*xabA*_T3$_{1/2}$C4A4T4$_{1/2}$_*ambS*_T1$_{1/2}$C/E2A2T2$_{1/2}$_*gxpS*_T4$_{1/2}$C/E5A5T5Te *tacI* and *araE* | This work |
| **A2_D11_D17** | pAP37_49_52 | ori p15A, *cm*$^R$, *araC-P*$_{BAD}$ *xldS*_C1A1T1$_{1/2}$_*xabA*_T3$_{1/2}$C4A4T4$_{1/2}$_*szeS*_T1$_{1/2}$C/E2A2T2$_{1/2}$_*gxpS*_T4$_{1/2}$C/E5A5T5Te *tacI* and *araE* | This work |
| **A2_D11_D18** | pAP37_49_53 | ori p15A, *cm*$^R$, *araC-P*$_{BAD}$ *xldS*_C1A1T1$_{1/2}$_*xabA*_T3$_{1/2}$C4A4T4$_{1/2}$_*szeS*_T4$_{1/2}$C/E5A5T5$_{1/2}$_*gxpS*_T4$_{1/2}$C/E5A5T5Te *tacI* and *araE* | This work |
| **A2_D11_D19** | pAP37_49_54 | ori p15A, *cm*$^R$, *araC-P*$_{BAD}$ *xldS*_C1A1T1$_{1/2}$_*xabA*_T3$_{1/2}$C4A4T4$_{1/2}$_*xtpS*_T2$_{1/2}$C3A3T3$_{1/2}$_*gxpS*_T4$_{1/2}$C/E5A5T5Te *tacI* and *araE* | This work |

| NRPS- | Plasmids | Genotype | Reference |
|---|---|---|---|
| **A2_D11_D20** | pAP37_49_55 | ori p15A, *cm*$^R$, *araC-P*$_{BAD}$ *xldS*_C1A1T1$_{1/2}$_*xabA*_T3$_{1/2}$C4A4T4$_{1/2}$_*xabA*_T3$_{1/2}$C4A4T4$_{1/2}$_*gxpS*_T4$_{1/2}$C/E5A5T5Te *tacI* and *araE* | This work |
| **A2_D11_D21** | pAP37_49_56 | ori p15A, *cm*$^R$, *araC-P*$_{BAD}$ *xldS*_C1A1T1$_{1/2}$_*xabA*_T3$_{1/2}$C4A4T4$_{1/2}$_*xabA*_T1$_{1/2}$C2A2T2$_{1/2}$_*gxpS*_T4$_{1/2}$C/E5A5T5Te *tacI* and *araE* | This work |
| **A2_D12_D16** | pAP37_50_51 | ori p15A, *cm*$^R$, *araC-P*$_{BAD}$ *xldS*_C1A1T1$_{1/2}$_*xabA*_T1$_{1/2}$C2A2T2$_{1/2}$_*ambS*_T1$_{1/2}$C/E2A2T2$_{1/2}$_*gxpS*_T4$_{1/2}$C/E5A5T5Te *tacI* and *araE* | This work |
| **A2_D12_D17** | pAP37_50_52 | ori p15A, *cm*$^R$, *araC-P*$_{BAD}$ *xldS*_C1A1T1$_{1/2}$_*xabA*_T1$_{1/2}$C2A2T2$_{1/2}$_*szeS*_T1$_{1/2}$C/E2A2T2$_{1/2}$_*gxpS*_T4$_{1/2}$C/E5A5T5Te *tacI* and *araE* | This work |
| **A2_D12_D18** | pAP37_50_53 | ori p15A, *cm*$^R$, *araC-P*$_{BAD}$ *xldS*_C1A1T1$_{1/2}$_*xabA*_T1$_{1/2}$C2A2T2$_{1/2}$_*szeS*_T4$_{1/2}$C/E5A5T5$_{1/2}$_*gxpS*_T4$_{1/2}$C/E5A5T5Te *tacI* and *araE* | This work |
| **A2_D12_D19** | pAP37_50_54 | ori p15A, *cm*$^R$, *araC-P*$_{BAD}$ *xldS*_C1A1T1$_{1/2}$_*xabA*_T1$_{1/2}$C2A2T2$_{1/2}$_*xtpS*_T2$_{1/2}$C3A3T3$_{1/2}$_*gxpS*_T4$_{1/2}$C/E5A5T5Te *tacI* and *araE* | This work |
| **A2_D12_D20** | pAP37_50_55 | ori p15A, *cm*$^R$, *araC-P*$_{BAD}$ *xldS*_C1A1T1$_{1/2}$_*xabA*_T1$_{1/2}$C2A2T2$_{1/2}$_*xabA*_T3$_{1/2}$C4A4T4$_{1/2}$_*gxpS*_T4$_{1/2}$C/E5A5T5Te *tacI* and *araE* | This work |
| **A2_D12_D21** | pAP37_50_56 | ori p15A, *cm*$^R$, *araC-P*$_{BAD}$ *xldS*_C1A1T1$_{1/2}$_*xabA*_T1$_{1/2}$C2A2T2$_{1/2}$_*xabA*_T1$_{1/2}$C2A2T2$_{1/2}$_*gxpS*_T4$_{1/2}$C/E5A5T5Te *tacI* and *araE* | This work |
| **A2_D13_D16** | pAP37_95_51 | ori p15A, *cm*$^R$, *araC-P*$_{BAD}$ *xldS*_C1A1T1$_{1/2}$_*xabB*_T7$_{1/2}$C8A8T8$_{1/2}$_*ambS*_T1$_{1/2}$C/E2A2T2$_{1/2}$_*gxpS*_T4$_{1/2}$C/E5A5T5Te *tacI* and *araE* | This work |
| **A2_D13_D17** | pAP37_95_52 | ori p15A, *cm*$^R$, *araC-P*$_{BAD}$ *xldS*_C1A1T1$_{1/2}$_*xabB*_T7$_{1/2}$C8A8T8$_{1/2}$_*szeS*_T1$_{1/2}$C/E2A2T2$_{1/2}$_*gxpS*_T4$_{1/2}$C/E5A5T5Te *tacI* and *araE* | This work |
| **A2_D13_D18** | pAP37_95_53 | ori p15A, *cm*$^R$, *araC-P*$_{BAD}$ *xldS*_C1A1T1$_{1/2}$_*xabB*_T7$_{1/2}$C8A8T8$_{1/2}$_*szeS*_T4$_{1/2}$C/E5A5T5$_{1/2}$_*gxpS*_T4$_{1/2}$C/E5A5T5Te *tacI* and *araE* | This work |

| NRPS- | Plasmids | Genotype | Reference |
|---|---|---|---|
| **A2_D13_D19** | pAP37_95_54 | ori p15A, *cm*$^R$, *araC-P*$_{BAD}$ *xldS*_C1A1T1$_{1/2}$_*xabB*_T7$_{1/2}$C8A8T8$_{1/2}$_*xtpS*_T2$_{1/2}$C3A3T3$_{1/2}$_*gxpS*_T4$_{1/2}$C/E5A5T5Te *tacI* and *araE* | This work |
| **A2_D13_D20** | pAP37_95_55 | ori p15A, *cm*$^R$, *araC-P*$_{BAD}$ *xldS*_C1A1T1$_{1/2}$_*xabB*_T7$_{1/2}$C8A8T8$_{1/2}$_*xabA*_T3$_{1/2}$C4A4T4$_{1/2}$_*gxpS*_T4$_{1/2}$C/E5A5T5Te *tacI* and *araE* | This work |
| **A2_D13_D21** | pAP37_95_56 | ori p15A, *cm*$^R$, *araC-P*$_{BAD}$ *xldS*_C1A1T1$_{1/2}$_*xabB*_T7$_{1/2}$C8A8T8$_{1/2}$_*xabA*_T1$_{1/2}$C2A2T2$_{1/2}$_*gxpS*_T4$_{1/2}$C/E5A5T5Te *tacI* and *araE* | This work |
| **A2_D14_D16** | pAP37_96_51 | ori p15A, *cm*$^R$, *araC-P*$_{BAD}$ *xldS*_C1A1T1$_{1/2}$_*xabA*_T4$_{1/2}$C5A5T5$_{1/2}$_*ambS*_T1$_{1/2}$C/E2A2T2$_{1/2}$_*gxpS*_T4$_{1/2}$C/E5A5T5Te *tacI* and *araE* | This work |
| **A2_D14_D17** | pAP37_96_52 | ori p15A, *cm*$^R$, *araC-P*$_{BAD}$ *xldS*_C1A1T1$_{1/2}$_*xabA*_T4$_{1/2}$C5A5T5$_{1/2}$_*szeS*_T1$_{1/2}$C/E2A2T2$_{1/2}$_*gxpS*_T4$_{1/2}$C/E5A5T5Te *tacI* and *araE* | This work |
| **A2_D14_D18** | pAP37_96_53 | ori p15A, *cm*$^R$, *araC-P*$_{BAD}$ *xldS*_C1A1T1$_{1/2}$_*xabA*_T4$_{1/2}$C5A5T5$_{1/2}$_*szeS*_T4$_{1/2}$C/E5A5T5$_{1/2}$_*gxpS*_T4$_{1/2}$C/E5A5T5Te *tacI* and *araE* | This work |
| **A2_D14_D19** | pAP37_96_54 | ori p15A, *cm*$^R$, *araC-P*$_{BAD}$ *xldS*_C1A1T1$_{1/2}$_*xabA*_T4$_{1/2}$C5A5T5$_{1/2}$_*xtpS*_T2$_{1/2}$C3A3T3$_{1/2}$_*gxpS*_T4$_{1/2}$C/E5A5T5Te *tacI* and *araE* | This work |
| **A2_D14_D20** | pAP37_96_55 | ori p15A, *cm*$^R$, *araC-P*$_{BAD}$ *xldS*_C1A1T1$_{1/2}$_*xabA*_T4$_{1/2}$C5A5T5$_{1/2}$_*xabA*_T3$_{1/2}$C4A4T4$_{1/2}$_*gxpS*_T4$_{1/2}$C/E5A5T5Te *tacI* and *araE* | This work |
| **A2_D14_D21** | pAP37_96_56 | ori p15A, *cm*$^R$, *araC-P*$_{BAD}$ *xldS*_C1A1T1$_{1/2}$_*xabA*_T4$_{1/2}$C5A5T5$_{1/2}$_*xabA*_T1$_{1/2}$C2A2T2$_{1/2}$_*gxpS*_T4$_{1/2}$C/E5A5T5Te *tacI* and *araE* | This work |
| **A2_D7_D22** | pAP37_45_98 | ori p15A, *cm*$^R$, *araC-P*$_{BAD}$ *xldS*_C1A1T1$_{1/2}$_*ambS*_T1$_{1/2}$C/E2A2T2$_{1/2}$_*xabB*_T7$_{1/2}$C8A8T8$_{1/2}$_*gxpS*_T4$_{1/2}$C/E5A5T5Te *tacI* and *araE* | This work |
| **A2_D8_D22** | pAP37_46_98 | ori p15A, *cm*$^R$, *araC-P*$_{BAD}$ *xldS*_C1A1T1$_{1/2}$_*szeS*_T1$_{1/2}$C/E2A2T2$_{1/2}$_*xabB*_T7$_{1/2}$C8A8T8$_{1/2}$_*gxpS*_T4$_{1/2}$C/E5A5T5Te *tacI* and *araE* | This work |

| NRPS- | Plasmids | Genotype | Reference |
|---|---|---|---|
| **A2_D9_D22** | pAP37_47_98 | ori p15A, *cm*$^R$, *araC-P*$_{BAD}$ *xldS*_C1A1T1$_{1/2}$_*szeS*_T4$_{1/2}$C/E5A5T5$_{1/2}$_*xabB*_T7$_{1/2}$C8A8T8$_{1/2}$_*gxpS*_T4$_{1/2}$C/E5A5T5Te *tacI* and *araE* | This work |
| **A2_D10_D22** | pAP37_48_98 | ori p15A, *cm*$^R$, *araC-P*$_{BAD}$ *xldS*_C1A1T1$_{1/2}$_*xtpS*_T2$_{1/2}$C3A3T3$_{1/2}$_*xabB*_T7$_{1/2}$C8A8T8$_{1/2}$_*gxpS*_T4$_{1/2}$C/E5A5T5Te *tacI* and *araE* | This work |
| **A2_D11_D22** | pAP37_49_98 | ori p15A, *cm*$^R$, *araC-P*$_{BAD}$ *xldS*_C1A1T1$_{1/2}$_*xabA*_T3$_{1/2}$C4A4T4$_{1/2}$_*xabB*_T7$_{1/2}$C8A8T8$_{1/2}$_*gxpS*_T4$_{1/2}$C/E5A5T5Te *tacI* and *araE* | This work |
| **A2_D12_D22** | pAP37_50_98 | ori p15A, *cm*$^R$, *araC-P*$_{BAD}$ *xldS*_C1A1T1$_{1/2}$_*xabA*_T1$_{1/2}$C2A2T2$_{1/2}$_*xabB*_T7$_{1/2}$C8A8T8$_{1/2}$_*gxpS*_T4$_{1/2}$C/E5A5T5Te *tacI* and *araE* | This work |
| **A2_D7_D23** | pAP37_45_99 | ori p15A, *cm*$^R$, *araC-P*$_{BAD}$ *xldS*_C1A1T1$_{1/2}$_*ambS*_T1$_{1/2}$C/E2A2T2$_{1/2}$_*xabA*_T4$_{1/2}$C5A5T5$_{1/2}$_*gxpS*_T4$_{1/2}$C/E5A5T5Te *tacI* and *araE* | This work |
| **A2_D8_D23** | pAP37_46_99 | ori p15A, *cm*$^R$, *araC-P*$_{BAD}$ *xldS*_C1A1T1$_{1/2}$_*szeS*_T1$_{1/2}$C/E2A2T2$_{1/2}$_*xabA*_T4$_{1/2}$C5A5T5$_{1/2}$_*gxpS*_T4$_{1/2}$C/E5A5T5Te *tacI* and *araE* | This work |
| **A2_D9_D23** | pAP37_47_99 | ori p15A, *cm*$^R$, *araC-P*$_{BAD}$ *xldS*_C1A1T1$_{1/2}$_*szeS*_T4$_{1/2}$C/E5A5T5$_{1/2}$_*xabA*_T4$_{1/2}$C5A5T5$_{1/2}$_*gxpS*_T4$_{1/2}$C/E5A5T5Te *tacI* and *araE* | This work |
| **A2_D10_D23** | pAP37_48_99 | ori p15A, *cm*$^R$, *araC-P*$_{BAD}$ *xldS*_C1A1T1$_{1/2}$_*xtpS*_T2$_{1/2}$C3A3T3$_{1/2}$_*xabA*_T4$_{1/2}$C5A5T5$_{1/2}$_*gxpS*_T4$_{1/2}$C/E5A5T5Te *tacI* and *araE* | This work |
| **A2_D11_D23** | pAP37_49_99 | ori p15A, *cm*$^R$, *araC-P*$_{BAD}$ *xldS*_C1A1T1$_{1/2}$_*xabA*_T3$_{1/2}$C4A4T4$_{1/2}$_*xabA*_T4$_{1/2}$C5A5T5$_{1/2}$_*gxpS*_T4$_{1/2}$C/E5A5T5Te *tacI* and *araE* | This work |
| **A2_D12_D23** | pAP37_50_99 | ori p15A, *cm*$^R$, *araC-P*$_{BAD}$ *xldS*_C1A1T1$_{1/2}$_*xabA*_T1$_{1/2}$C2A2T2$_{1/2}$_*xabA*_T4$_{1/2}$C5A5T5$_{1/2}$_*gxpS*_T4$_{1/2}$C/E5A5T5Te *tacI* and *araE* | This work |
| **A2_D13_D22** | pAP37_95_98 | ori p15A, *cm*$^R$, *araC-P*$_{BAD}$ *xldS*_C1A1T1$_{1/2}$_*xabB*_T7$_{1/2}$C8A8T8$_{1/2}$_*xabB_T7*$_{1/2}$*C8A8T8*$_{1/2}$_*gxpS*_T4$_{1/2}$C/E5A5T5Te *tacI* and *araE* | This work |

| NRPS- | Plasmids | Genotype | Reference |
|---|---|---|---|
| **A2_D13_D23** | pAP37_95_99 | ori p15A, *cm*$^R$, *araC-P*$_{BAD}$ *xldS*_C1A1T1$_{1/2}$_*xabB*_T7$_{1/2}$C8A8T8$_{1/2}$_*xabA*_T4$_{1/2}$C5A5T5$_{1/2}$_*gxpS*_T4$_{1/2}$C/E5A5T5Te *tacI* and *araE* | This work |
| **A2_D14_D22** | pAP37_96_98 | ori p15A, *cm*$^R$, *araC-P*$_{BAD}$ *xldS*_C1A1T1$_{1/2}$_*xabA_T4*$_{1/2}$*C5A5T5*$_{1/2}$_*xabB_T7*$_{1/2}$*C8A8T8*$_{1/2}$_*gxpS*_T4$_{1/2}$C/E5A5T5Te *tacI* and *araE* | This work |
| **A2_D14_D23** | pAP37_96_99 | ori p15A, *cm*$^R$, *araC-P*$_{BAD}$ *xldS*_C1A1T1$_{1/2}$_*xabA_T4*$_{1/2}$*C5A5T5*$_{1/2}$_*xabA*_T4$_{1/2}$C5A5T5$_{1/2}$_*gxpS*_T4$_{1/2}$C/E5A5T5Te *tacI* and *araE* | This work |
| **A2_D15_D16** | pAP37_89_51 | ori p15A, *cm*$^R$, *araC-P*$_{BAD}$ *xldS*_C1A1T1$_{1/2}$_*fitAB*_T3$_{1/2}$EC5A5T5$_{1/2}$_*ambS*_T1$_{1/2}$C/E2A2T2$_{1/2}$_*gxpS*_T4$_{1/2}$C/E5A5T5Te *tacI* and *araE* | This work |
| **A2_D15_D17** | pAP37_89_52 | ori p15A, *cm*$^R$, *araC-P*$_{BAD}$ *xldS*_C1A1T1$_{1/2}$_*fitAB*_T3$_{1/2}$EC5A5T5$_{1/2}$_*szeS*_T1$_{1/2}$C/E2A2T2$_{1/2}$_*gxpS*_T4$_{1/2}$C/E5A5T5Te *tacI* and *araE* | This work |
| **A2_D15_D18** | pAP37_89_53 | ori p15A, *cm*$^R$, *araC-P*$_{BAD}$ *xldS*_C1A1T1$_{1/2}$_*fitAB*_T3$_{1/2}$EC5A5T5$_{1/2}$_*szeS*_T4$_{1/2}$C/E5A5T5$_{1/2}$_*gxpS*_T4$_{1/2}$C/E5A5T5Te *tacI* and *araE* | This work |
| **A2_D15_D19** | pAP37_89_54 | ori p15A, *cm*$^R$, *araC-P*$_{BAD}$ *xldS*_C1A1T1$_{1/2}$_*fitAB*_T3$_{1/2}$EC5A5T5$_{1/2}$_*xtpS*_T2$_{1/2}$C3A3T3$_{1/2}$_*gxpS*_T4$_{1/2}$C/E5A5T5Te *tacI* and *araE* | This work |
| **A2_D15_D20** | pAP37_89_55 | ori p15A, *cm*$^R$, *araC-P*$_{BAD}$ *xldS*_C1A1T1$_{1/2}$_*fitAB*_T3$_{1/2}$EC5A5T5$_{1/2}$_*xabA*_T3$_{1/2}$C4A4T4$_{1/2}$_*gxpS*_T4$_{1/2}$C/E5A5T5Te *tacI* and *araE* | This work |
| **A2_D15_D21** | pAP37_89_56 | ori p15A, *cm*$^R$, *araC-P*$_{BAD}$ *xldS*_C1A1T1$_{1/2}$_*fitAB*_T3$_{1/2}$EC5A5T5$_{1/2}$_*xabA*_T1$_{1/2}$C2A2T2$_{1/2}$_*gxpS*_T4$_{1/2}$C/E5A5T5Te *tacI* and *araE* | This work |
| **A2_D7_D24** | pAP37_45_92 | ori p15A, *cm*$^R$, *araC-P*$_{BAD}$ *xldS*_C1A1T1$_{1/2}$_*ambS*_T1$_{1/2}$C/E2A2T2$_{1/2}$_*fitAB*_T3$_{1/2}$EC5A5T5$_{1/2}$_*gxpS*_T4$_{1/2}$C/E5A5T5Te *tacI* and *araE* | This work |
| **A2_D8_D24** | pAP37_46_92 | ori p15A, *cm*$^R$, *araC-P*$_{BAD}$ *xldS*_C1A1T1$_{1/2}$_*szeS*_T1$_{1/2}$C/E2A2T2$_{1/2}$_*fitAB*_T3$_{1/2}$EC5A5T5$_{1/2}$_*gxpS*_T4$_{1/2}$C/E5A5T5Te *tacI* and *araE* | This work |

| NRPS- | Plasmids | Genotype | Reference |
|---|---|---|---|
| **A2_D9_D24** | pAP37_47_92 | ori p15A, *cm*$^R$, *araC-P*$_{BAD}$ *xldS*_C1A1T1$_{1/2}$_*szeS*_T4$_{1/2}$C/E5A5T5$_{1/2}$_*fitAB*_T3$_{1/2}$EC5A5T5$_{1/2}$_*gxpS*_T4$_{1/2}$C/E5A5T5Te *tacI* and *araE* | This work |
| **A2_D10_D24** | pAP37_48_92 | ori p15A, *cm*$^R$, *araC-P*$_{BAD}$ *xldS*_C1A1T1$_{1/2}$_*xtpS*_T2$_{1/2}$C3A3T3$_{1/2}$_*fitAB*_T3$_{1/2}$EC5A5T5$_{1/2}$_*gxpS*_T4$_{1/2}$C/E5A5T5Te *tacI* and *araE* | This work |
| **A2_D11_D24** | pAP37_49_92 | ori p15A, *cm*$^R$, *araC-P*$_{BAD}$ *xldS*_C1A1T1$_{1/2}$_*xabA*_T3$_{1/2}$C4A4T4$_{1/2}$_*fitAB*_T3$_{1/2}$EC5A5T5$_{1/2}$_*gxpS*_T4$_{1/2}$C/E5A5T5Te *tacI* and *araE* | This work |
| **A2_D12_D24** | pAP37_50_92 | ori p15A, *cm*$^R$, *araC-P*$_{BAD}$ *xldS*_C1A1T1$_{1/2}$_*xabA*_T1$_{1/2}$C2A2T2$_{1/2}$_*fitAB*_T3$_{1/2}$EC5A5T5$_{1/2}$_*gxpS*_T4$_{1/2}$C/E5A5T5Te *tacI* and *araE* | This work |
| **A2_D15_D24** | pAP37_89_92 | ori p15A, *cm*$^R$, *araC-P*$_{BAD}$ *xldS*_C1A1T1$_{1/2}$_*fitAB*_T3$_{1/2}$EC5A5T5$_{1/2}$_*fitAB*_T3$_{1/2}$EC5A5T5$_{1/2}$_*gxpS*_T4$_{1/2}$C/E5A5T5Te *tacI* and *araE* | This work |
| **A2_D15_D22** | pAP37_89_98 | ori p15A, *cm*$^R$, *araC-P*$_{BAD}$ *xldS*_C1A1T1$_{1/2}$_*fitAB*_T3$_{1/2}$EC5A5T5$_{1/2}$_*xabB*_T7$_{1/2}$C8A8T8$_{1/2}$_*gxpS*_T4$_{1/2}$C/E5A5T5Te *tacI* and *araE* | This work |
| **A2_D15_D23** | pAP37_89_99 | ori p15A, *cm*$^R$, *araC-P*$_{BAD}$ *xldS*_C1A1T1$_{1/2}$_*fitAB*_T3$_{1/2}$EC5A5T5$_{1/2}$_*xabA*_T4$_{1/2}$C5A5T5$_{1/2}$_*gxpS*_T4$_{1/2}$C/E5A5T5Te *tacI* and *araE* | This work |
| **A2_D13_D24** | pAP37_95_92 | ori p15A, *cm*$^R$, *araC-P*$_{BAD}$ *xldS*_C1A1T1$_{1/2}$_*xabB*_T7$_{1/2}$C8A8T8$_{1/2}$_*fitAB*_T3$_{1/2}$EC5A5T5$_{1/2}$_*gxpS*_T4$_{1/2}$C/E5A5T5Te *tacI* and *araE* | This work |
| **A2_D14_D24** | pAP37_96_92 | ori p15A, *cm*$^R$, *araC-P*$_{BAD}$ *xldS*_C1A1T1$_{1/2}$_*xabA*_T4$_{1/2}$C5A5T5$_{1/2}$_*fitAB*_T3$_{1/2}$EC5A5T5$_{1/2}$_*gxpS*_T4$_{1/2}$C/E5A5T5Te *tacI* and *araE* | This work |
| **A3_D25** | pAP106_107 | ori p15A, *cm*$^R$, *araC-P*$_{BAD}$ *xldS*_C1A1T1$_{1/2}$_*xabA*_T1$_{1/2}$C2A2T2$_{1/2}$_*gxpS*_T4$_{1/2}$C/E5A5T5Te *tacI* and *araE* | This work |
| **A3_D26** | pAP106_108 | ori p15A, *cm*$^R$, *araC-P*$_{BAD}$ *xabA*_C1A1T1$_{1/2}$_*xabA*_T1$_{1/2}$C2A2T2$_{1/2}$_*gxpS*_T4$_{1/2}$C/E5A5T5Te *tacI* and *araE* | This work |
| **A3_D27** | pAP106_109 | ori p15A, *cm*$^R$, *araC-P*$_{BAD}$ *ambS*_C1A1T1$_{1/2}$_*xabA*_T1$_{1/2}$C2A2T2$_{1/2}$_*gxpS*_T4$_{1/2}$C/E5A5T5Te *tacI* and *araE* | This work |

| NRPS- | Plasmids | Genotype | Reference |
|---|---|---|---|
| **A3_D28** | pAP106_110 | ori p15A, *cm*$^R$, *araC-P*$_{BAD}$ XISHV2_07515_C1A1T1$_{1/2}$_*xabA*_T1$_{1/2}$C2A2T2$_{1/2}$_*gxpS*_T4$_{1/2}$C/E5A5T5Te *tacI* and *araE* | This work |
| **A4_D29** | pAP119_113 | ori p15A, *cm*$^R$, *araC-P*$_{BAD}$ *xabC*_C1A1T1$_{1/2}$_*xabC*_T1$_{1/2}$C/E2A2T2$_{1/2}$_*xabC*_T3$_{1/2}$C4A4T4TeTe *tacI* and *araE* | This work |
| **A4_D30** | pAP119_114 | ori p15A, *cm*$^R$, *araC-P*$_{BAD}$ *xabC*_C1A1T1$_{1/2}$_*xabC*_T2$_{1/2}$C3A3T3$_{1/2}$_*xabC*_T3$_{1/2}$C4A4T4TeTe *tacI* and *araE* | This work |
| **A4_D31_D33** | pAP119_115_117 | ori p15A, *cm*$^R$, *araC-P*$_{BAD}$ *xabC*_C1A1T1$_{1/2}$_*xabC*_T1$_{1/2}$C/E2A2T2$_{1/2}$_*xabC*_T11$_{1/2}$C/E2A2T2$_{1/2}$_*xabC*_T3$_{1/2}$C4A4T4TeTe *tacI* and *araE* | This work |
| **A4_D31_D34** | pAP119_115_118 | ori p15A, *cm*$^R$, *araC-P*$_{BAD}$ *xabC*_C1A1T1$_{1/2}$_*xabC*_T1$_{1/2}$C/E2A2T2$_{1/2}$_*xabC*_T2$_{1/2}$C3A3T3$_{1/2}$_*xabC*_T3$_{1/2}$C4A4T4TeTe *tacI* and *araE* | This work |
| **A4_D32_D33** | pAP119_116_117 | ori p15A, *cm*$^R$, *araC-P*$_{BAD}$ *xabC*_C1A1T1$_{1/2}$_*xabC*_T2$_{1/2}$C/E3A3T3$_{1/2}$_*xabC*_T1$_{1/2}$C/E2A2T2$_{1/2}$_*xabC*_T3$_{1/2}$C4A4T4TeTe *tacI* and *araE* | This work |
| **A4_D32_D34** | pAP119_116_118 | ori p15A, *cm*$^R$, *araC-P*$_{BAD}$ *xabC*_C1A1T1$_{1/2}$_*xabC*_T2$_{1/2}$C3A3T3$_{1/2}$_*xabC*_T2$_{1/2}$C3A3T3$_{1/2}$_*xabC*_T3$_{1/2}$C4A4T4TeTe *tacI* and *araE* | This work |
| **A4_D1** | pAP119_39 | ori p15A, *cm*$^R$, *araC-P*$_{BAD}$ *xabC*_C1A1T1$_{1/2}$_ambS_T1$_{1/2}$C/E2A2T2$_{1/2}$_*xabC*_T3$_{1/2}$C4A4T4TeTe *tacI* and *araE* | This work |
| **A4_D2** | pAP119_40 | ori p15A, *cm*$^R$, *araC-P*$_{BAD}$ *xabC*_C1A1T1$_{1/2}$_szeS_T1$_{1/2}$C/E2A2T2$_{1/2}$_*xabC*_T3$_{1/2}$C4A4T4TeTe *tacI* and *araE* | This work |
| **A4_D3** | pAP119_41 | ori p15A, *cm*$^R$, *araC-P*$_{BAD}$ *xabC*_C1A1T1$_{1/2}$_szeS_T4$_{1/2}$C/E5A5T5$_{1/2}$_*xabC*_T3$_{1/2}$C4A4T4TeTe *tacI* and *araE* | This work |
| **A4_D4** | pAP119_42 | ori p15A, *cm*$^R$, *araC-P*$_{BAD}$ *xabC*_C1A1T1$_{1/2}$_xtpS_T2$_{1/2}$C3A3T3$_{1/2}$_*xabC*_T3$_{1/2}$C4A4T4TeTe *tacI* and *araE* | This work |
| **A4_D5** | pAP119_43 | ori p15A, *cm*$^R$, *araC-P*$_{BAD}$ *xabC*_C1A1T1$_{1/2}$_xabA_T3$_{1/2}$C4A4T4$_{1/2}$_gxpS_*xabC*_T3$_{1/2}$C4A4T4TeTe *tacI* and *araE* | This work |
| **A4_D6** | pAP119_44 | ori p15A, *cm*$^R$, *araC-P*$_{BAD}$ *xabC*_C1A1T1$_{1/2}$_xabA_T1$_{1/2}$C2A2T2$_{1/2}$_gxpS_*xabC*_T3$_{1/2}$C4A4T4TeTe *tacI* and *araE* | This work |

| NRPS- | Plasmids | Genotype | Reference |
|---|---|---|---|
| **A4_D7_D16** | pAP119_45_51 | ori p15A, *cm*$^R$, *araC-P*$_{BAD}$ *xab*C_C1A1T1$_{1/2}$_*ambS*_T1$_{1/2}$C/E2A2T2$_{1/2}$ _*ambS*_T1$_{1/2}$C/E2A2T2$_{1/2}$_*xabC*_T3$_{1/2}$C4A 4T4TeTe *tacI* and *araE* | This work |
| **A4_D7_D17** | pAP119_45_52 | ori p15A, *cm*$^R$, *araC-P*$_{BAD}$ *xabC*_C1A1T1$_{1/2}$_*ambS*_T1$_{1/2}$C/E2A2T2$_{1/2}$ _*szeS*_T1$_{1/2}$C/E2A2T2$_{1/2}$_*xabC*_T3$_{1/2}$C4A4 T4TeTe *tacI* and *araE* | This work |
| **A4_D7_D18** | pAP119_45_53 | ori p15A, *cm*$^R$, *araC-P*$_{BAD}$ *xabC*_C1A1T1$_{1/2}$_*ambS*_T1$_{1/2}$C/E2A2T2$_{1/2}$ _*szeS*_T4$_{1/2}$C/E5A5T5$_{1/2}$_*xabC*_T3$_{1/2}$C4A4 T4TeTe *tacI* and *araE* | This work |
| **A4_D7_D19** | pAP119_45_54 | ori p15A, *cm*$^R$, *araC-P*$_{BAD}$ *xabC*_C1A1T1$_{1/2}$_*ambS*_T1$_{1/2}$C/E2A2T2$_{1/2}$ _*xtpS*_T2$_{1/2}$C3A3T3$_{1/2}$_*xabC*_T3$_{1/2}$C4A4T4 TeTe *tacI* and *araE* | This work |
| **A4_D7_D20** | pAP119_45_55 | ori p15A, *cm*$^R$, *araC-P*$_{BAD}$ *xabC*_C1A1T1$_{1/2}$_*ambS*_T1$_{1/2}$C/E2A2T2$_{1/2}$ _*xabA*_T3$_{1/2}$C4A4T4$_{1/2}$_*xabC*_T3$_{1/2}$C4A4T 4TeTe *tacI* and *araE* | This work |
| **A4_D7_D21** | pAP119_45_56 | ori p15A, *cm*$^R$, *araC-P*$_{BAD}$ *xabC*_C1A1T1$_{1/2}$_*ambS*_T1$_{1/2}$C/E2A2T2$_{1/2}$ _*xabA*_T1$_{1/2}$C2A2T2$_{1/2}$_*xabC*_T3$_{1/2}$C4A4T 4TeTe *tacI* and *araE* | This work |
| **A4_D8_D16** | pAP119_46_51 | ori p15A, *cm*$^R$, *araC-P*$_{BAD}$ *xabC*_C1A1T1$_{1/2}$_*szeS*_T1$_{1/2}$C/E2A2T2$_{1/2}$_ *ambS*_T1$_{1/2}$C/E2A2T2$_{1/2}$_*xabC*_T3$_{1/2}$C4A4 T4TeTe *tacI* and *araE* | This work |
| **A4_D8_D17** | pAP119_46_52 | ori p15A, *cm*$^R$, *araC-P*$_{BAD}$ *xabC*_C1A1T1$_{1/2}$_*szeS*_T1$_{1/2}$C/E2A2T2$_{1/2}$_ *szeS*_T1$_{1/2}$C/E2A2T2$_{1/2}$_*xabC*_T3$_{1/2}$C4A4T 4TeTe *tacI* and *araE* | This work |
| **A4_D8_D18** | pAP119_46_53 | ori p15A, *cm*$^R$, *araC-P*$_{BAD}$ *xabC*_C1A1T1$_{1/2}$_*szeS*_T1$_{1/2}$C/E2A2T2$_{1/2}$_ *szeS*_T4$_{1/2}$C/E5A5T5$_{1/2}$_*xabC*_T3$_{1/2}$C4A4T 4TeTe *tacI* and *araE* | This work |
| **A4_D8_D19** | pAP119_46_54 | ori p15A, *cm*$^R$, *araC-P*$_{BAD}$ *xldS*_C1A1T1$_{1/2}$_*szeS*_T1$_{1/2}$C/E2A2T2$_{1/2}$_ *xtpS*_T2$_{1/2}$C3A3T3$_{1/2}$_xabC_T31/2C4A4T4 TeTe *tacI* and *araE* | This work |
| **A4_D8_D20** | pAP119_46_55 | ori p15A, *cm*$^R$, *araC-P*$_{BAD}$ *xabC*_C1A1T1$_{1/2}$_*szeS*_T1$_{1/2}$C/E2A2T2$_{1/2}$_ *xabA*_T3$_{1/2}$C4A4T4$_{1/2}$_*xabC*_T3$_{1/2}$C4A4T4 TeTe *tacI* and *araE* | This work |

| NRPS- | Plasmids | Genotype | Reference |
|---|---|---|---|
| **A4_D8_D21** | pAP119_46_56 | ori p15A, *cm*$^R$, *araC-P*$_{BAD}$ *xabC*_C1A1T1$_{1/2}$_*szeS*_T1$_{1/2}$C/E2A2T2$_{1/2}$_ *xabA*_T1$_{1/2}$C2A2T2$_{1/2}$_*xabC*_T3$_{1/2}$C4A4T4TeTe *tacI* and *araE* | This work |
| **A4_D9_D16** | pAP119_47_51 | ori p15A, *cm*$^R$, *araC-P*$_{BAD}$ *xabC_C1A1T11/2_szeS*_T4$_{1/2}$C/E5A5T5$_{1/2}$ _*ambS*_T1$_{1/2}$C/E2A2T2$_{1/2}$_xabC_T31/2C4A4T4TeTe *tacI* and *araE* | This work |
| **A4_D9_D17** | pAP119_47_52 | ori p15A, *cm*$^R$, *araC-P*$_{BAD}$ *xabC*_C1A1T1$_{1/2}$_*szeS*_T4$_{1/2}$C/E5A5T5$_{1/2}$_ *szeS*_T1$_{1/2}$C/E2A2T2$_{1/2}$_*xabC*_T3$_{1/2}$C4A4T4TeTe *tacI* and *araE* | This work |
| **A4_D9_D18** | pAP119_47_53 | ori p15A, *cm*$^R$, *araC-P*$_{BAD}$ *xabC*_C1A1T1$_{1/2}$_*szeS*_T4$_{1/2}$C/E5A5T5$_{1/2}$_ *szeS*_T4$_{1/2}$C/E5A5T5$_{1/2}$_*xabC*_T3$_{1/2}$C4A4T4TeTe *tacI* and *araE* | This work |
| **A4_D9_D19** | pAP119_47_54 | ori p15A, *cm*$^R$, *araC-P*$_{BAD}$ *xabC*_C1A1T1$_{1/2}$_*szeS*_T4$_{1/2}$C/E5A5T5$_{1/2}$_ *xtpS*_T2$_{1/2}$C3A3T3$_{1/2}$_*xabC*_T3$_{1/2}$C4A4T4TeTe *tacI* and *araE* | This work |
| **A4_D9_D20** | pAP119_47_55 | ori p15A, *cm*$^R$, *araC-P*$_{BAD}$ *xabC*_C1A1T1$_{1/2}$_*szeS*_T4$_{1/2}$C/E5A5T5$_{1/2}$_ *xabA*_T3$_{1/2}$C4A4T4$_{1/2}$_*xabC*_T3$_{1/2}$C4A4T4TeTe *tacI* and *araE* | This work |
| **A4_D9_D21** | pAP119_47_56 | ori p15A, *cm*$^R$, *araC-P*$_{BAD}$ *xabC*_C1A1T1$_{1/2}$_*szeS*_T4$_{1/2}$C/E5A5T5$_{1/2}$_ *xabA*_T1$_{1/2}$C2A2T2$_{1/2}$_*xabC*_T3$_{1/2}$C4A4T4TeTe *tacI* and *araE* | This work |
| **A4_D10_D16** | pAP119_48_51 | ori p15A, *cm*$^R$, *araC-P*$_{BAD}$ *xabC*_C1A1T1$_{1/2}$_*xtpS*_T2$_{1/2}$C3A3T3$_{1/2}$_*ambS*_T1$_{1/2}$C/E2A2T2$_{1/2}$_*xabC*_T3$_{1/2}$C4A4T4TeTe *tacI* and *araE* | This work |
| **A4_D10_D17** | pAP119_48_52 | ori p15A, *cm*$^R$, *araC-P*$_{BAD}$ *xabC*_C1A1T1$_{1/2}$_*xtpS*_T2$_{1/2}$C3A3T3$_{1/2}$_*szeS*_T1$_{1/2}$C/E2A2T2$_{1/2}$_*xabC*_T3$_{1/2}$C4A4T4TeTe *tacI* and *araE* | This work |
| **A4_D10_D18** | pAP119_48_53 | ori p15A, cmR, araC-PBAD *xabC*_C1A1T1$_{1/2}$_xtpS_T21/2C3A3T31/2_ szeS_T41/2C/E5A5T51/2_*xabC*_T3$_{1/2}$C4A4T4TeTe tacI and araE | This work |
| **A4_D10_D19** | pAP119_48_54 | ori p15A, *cm*$^R$, *araC-P*$_{BAD}$ *xabC*_C1A1T1$_{1/2}$_*xtpS*_T2$_{1/2}$C3A3T3$_{1/2}$_*xtpS*_T2$_{1/2}$C3A3T3$_{1/2}$_*xabC*_T3$_{1/2}$C4A4T4TeTe *tacI* and *araE* | This work |

| NRPS- | Plasmids | Genotype | Reference |
|---|---|---|---|
| **A4_D10_D20** | pAP119_48_55 | ori p15A, $cm^R$, *araC-P*$_{BAD}$ *xabC*_C1A1T1$_{1/2}$_*xtpS*_T2$_{1/2}$C3A3T3$_{1/2}$_*xabA*_T3$_{1/2}$C4A4T4$_{1/2}$_*xabC*_T3$_{1/2}$C4A4T4TeTe *tacI* and *araE* | This work |
| **A4_D10_D21** | pAP119_48_56 | ori p15A, $cm^R$, *araC-P*$_{BAD}$ *xabC*_C1A1T1$_{1/2}$_*xtpS*_T2$_{1/2}$C3A3T3$_{1/2}$_*xabA*_T1$_{1/2}$C2A2T2$_{1/2}$_*xabC*_T3$_{1/2}$C4A4T4TeTe *tacI* and *araE* | This work |
| **A4_D11_D16** | pAP119_49_51 | ori p15A, $cm^R$, *araC-P*$_{BAD}$ *xabC*_C1A1T1$_{1/2}$_*xabA*_T3$_{1/2}$C4A4T4$_{1/2}$_*ambS*_T1$_{1/2}$C/E2A2T2$_{1/2}$_*xabC*_T3$_{1/2}$C4A4T4TeTe *tacI* and *araE* | This work |
| **A4_D11_D17** | pAP119_49_52 | ori p15A, $cm^R$, *araC-P*$_{BAD}$ *xabC*_C1A1T1$_{1/2}$_*xabA*_T3$_{1/2}$C4A4T4$_{1/2}$_*szeS*_T1$_{1/2}$C/E2A2T2$_{1/2}$_*xabC*_T3$_{1/2}$C4A4T4TeTe *tacI* and *araE* | This work |
| **A4_D11_D18** | pAP119_49_53 | ori p15A, $cm^R$, *araC-P*$_{BAD}$ *xabC*_C1A1T1$_{1/2}$_*xabA*_T3$_{1/2}$C4A4T4$_{1/2}$_*szeS*_T4$_{1/2}$C/E5A5T5$_{1/2}$_*xabC*_T3$_{1/2}$C4A4T4TeTe *tacI* and *araE* | This work |
| **A4_D11_D19** | pAP119_49_54 | ori p15A, $cm^R$, *araC-P*$_{BAD}$ *xabC*_C1A1T1$_{1/2}$_*xabA*_T3$_{1/2}$C4A4T4$_{1/2}$_*xtpS*_T2$_{1/2}$C3A3T3$_{1/2}$_*xabC*_T3$_{1/2}$C4A4T4TeTe *tacI* and *araE* | This work |
| **A4_D11_D20** | pAP119_49_55 | ori p15A, $cm^R$, *araC-P*$_{BAD}$ *xabC*_C1A1T1$_{1/2}$_*xabA*_T3$_{1/2}$C4A4T4$_{1/2}$_*xabA*_T3$_{1/2}$C4A4T4$_{1/2}$_*xabC*_T3$_{1/2}$C4A4T4TeTe *tacI* and *araE* | This work |
| **A4_D11_D21** | pAP119_49_56 | ori p15A, $cm^R$, *araC-P*$_{BAD}$ *xabC*_C1A1T1$_{1/2}$_*xabA*_T3$_{1/2}$C4A4T4$_{1/2}$_*xabA*_T1$_{1/2}$C2A2T2$_{1/2}$_*xabC*_T3$_{1/2}$C4A4T4TeTe *tacI* and *araE* | This work |
| **A4_D12_D16** | pAP119_50_51 | ori p15A, $cm^R$, *araC-P*$_{BAD}$ *xabC*_C1A1T1$_{1/2}$_*xabA*_T1$_{1/2}$C2A2T2$_{1/2}$_*ambS*_T1$_{1/2}$C/E2A2T2$_{1/2}$_*xabC*_T3$_{1/2}$C4A4T4TeTe *tacI* and *araE* | This work |
| **A4_D12_D17** | pAP119_50_52 | ori p15A, $cm^R$, *araC-P*$_{BAD}$ *xabC*_C1A1T1$_{1/2}$_*xabA*_T1$_{1/2}$C2A2T2$_{1/2}$_*szeS*_T1$_{1/2}$C/E2A2T2$_{1/2}$_*xabC*_T3$_{1/2}$C4A4T4TeTe *tacI* and *araE* | This work |
| **A4_D12_D18** | pAP119_50_53 | ori p15A, $cm^R$, *araC-P*$_{BAD}$ *xabC_C1A1T1*$_{1/2}$_*xabA*_T1$_{1/2}$C2A2T2$_{1/2}$_*szeS*_T4$_{1/2}$C/E5A5T5$_{1/2}$_*xabC*_T3$_{1/2}$C4A4T4TeTe *tacI* and *araE* | This work |

| NRPS- | Plasmids | Genotype | Reference |
|---|---|---|---|
| **A4_D12_D19** | pAP119_50_54 | ori p15A, *cm*$^R$, *araC-P*$_{BAD}$ *xabC*_C1A1T1$_{1/2}$_*xabA*_T1$_{1/2}$C2A2T2$_{1/2}$_*xtpS*_T2$_{1/2}$C3A3T3$_{1/2}$_*xabC*_T3$_{1/2}$C4A4T4TeTe *tacI* and *araE* | This work |
| **A4_D12_D20** | pAP119_50_55 | ori p15A, *cm*$^R$, *araC-P*$_{BAD}$ *xabC*_C1A1T1$_{1/2}$_*xabA*_T1$_{1/2}$C2A2T2$_{1/2}$_*xabA*_T3$_{1/2}$C4A4T4$_{1/2}$_*xabC*_T3$_{1/2}$C4A4T4TeTe *tacI* and *araE* | This work |
| **A4_D12_D21** | pAP119_50_56 | ori p15A, *cm*$^R$, *araC-P*$_{BAD}$ *xabC*_C1A1T1$_{1/2}$_*xabA*_T1$_{1/2}$C2A2T2$_{1/2}$_*xabA*_T1$_{1/2}$C2A2T2$_{1/2}$_*xabC*_T3$_{1/2}$C4A4T4TeTe *tacI* and *araE* | This work |

**Table S3.** List of all NRPS donors created in this work, with their respective GGA overhangs, strain origin, NRPS and module number, amino acids and corresponding plasmid.

| Donor | Overhang | Origin strain | NRPS & Module Nr. | Amino acid | Plasmid |
|---|---|---|---|---|---|
| **D1** | A/B | *X. miraniensis* DSM 17902 | AmbS M2 | Glu | pAP39 |
| **D2** | A/B | *X. szentirmaii* DSM 16338 | SzeS M2 | Thr | pAP40 |
| **D3** | A/B | *X. szentirmaii* DSM 16338 | SzeS M5 | Tyr | pAP41 |
| **D4** | A/B | *X. nematophila* ATCC 19061 | XtpS M3 | Val | pAP42 |
| **D5** | A/B | *X. doucetiae* DSM 17909 | XabABC M4 | Leu | pAP43 |
| **D6** | A/B | *X. doucetiae* DSM 17909 | XabABC M2 | Ala | pAP44 |
| **D7** | A/C | *X. miraniensis* DSM 17902 | AmbS M2 | Glu | pAP45 |
| **D8** | A/C | *X. szentirmaii* DSM 16338 | SzeS M2 | Thr | pAP46 |
| **D9** | A/C | *X. szentirmaii* DSM 16338 | SzeS M5 | Tyr | pAP47 |
| **D10** | A/C | *X. nematophila* ATCC 19061 | XtpS M3 | Val | pAP48 |
| **D11** | A/C | *X. doucetiae* DSM 17909 | XabABC M4 | Leu | pAP49 |
| **D12** | A/C | *X. doucetiae* DSM 17909 | XabABC M2 | Ala | pAP50 |
| **D13** | A/C | *X. innexi* DSM 16336 | XabABC M8 | Val | pAP95 |

| Donor | Overhang | Origin strain | NRPS & Module Nr. | Amino acid | Plasmid |
|---|---|---|---|---|---|
| **D14** | A/C | *X. beddingii* DSM 4764 | XabABC M5 | Val | pAP96 |
| **D15** | A/C | *X. innexi* DSM 16336 | FitAB M4 | Tyr | pAP89 |
| **D16** | C/B | *X. miraniensis* DSM 17902 | AmbS M2 | Glu | pAP51 |
| **D17** | C/B | *X. szentirmaii* DSM 16338 | SzeS M2 | Thr | pAP52 |
| **D18** | C/B | *X. szentirmaii* DSM 16338 | SzeS M5 | Tyr | pAP53 |
| **D19** | C/B | *X. nematophila* ATCC 19061 | XtpS M3 | Val | pAP54 |
| **D20** | C/B | *X. doucetiae* DSM 17909 | XabABC M4 | Leu | pAP55 |
| **D21** | C/B | *X. doucetiae* DSM 17909 | XabABC M2 | Ala | pAP56 |
| **D22** | C/B | *X. innexi* DSM 16336 | XabABC M8 | Val | pAP98 |
| **D23** | C/B | *X. beddingii* DSM 4764 | XabABC M5 | Val | pAP99 |
| **D24** | C/B | *X. innexi* DSM 16336 | FitAB M4 | Tyr | pAP92 |
| **D25** | S/A | *X. indica* DSM 17382 | XldS M1 | FA-Glu | pAP107 |
| **D26** | S/A | *X. doucetiae* DSM 17909 | XabABC M1 | FA-Pro | pAP108 |
| **D27** | S/A | *X. miraniensis* DSM 17902 | AmbS M1 | Ser | pAP109 |
| **D28** | S/A | *X. ishibashii* DSM 22670 | LC536431 M1 | Leu | pAP110 |
| **D29** | A/B | *X. doucetiae* DSM 17909 | XabABC M11 | β-Ala | pAP113 |
| **D30** | A/B | *X. doucetiae* DSM 17909 | XabABC M12 | Pro | pAP114 |
| **D31** | A/C | *X. doucetiae* DSM 17909 | XabABC M11 | β-Ala | pAP115 |
| **D32** | A/C | *X. doucetiae* DSM 17909 | XabABC M12 | Pro | pAP116 |
| **D33** | C/B | *X. doucetiae* DSM 17909 | XabABC M11 | β-Ala | pAP117 |
| **D34** | C/B | *X. doucetiae* DSM 17909 | XabABC M12 | Pro | pAP118 |

**Table S4.** Primer and template used in this work to generate the indicated plasmids. The sizes of the PCR products are shown below the template.

| Plasmids | Oligo-nucleotide | Sequence (5' → 3') | Template<br>Product size in bp |
|---|---|---|---|
| **pAP32** | AP63 | TTTTTGGGCTAACAGGAGGAATTCCATG AATATGACACGTAACCATACAT | *X. innexi* gDNA<br>3091 |
| | AP64.2 | TCCTCCAAGCTCAAAGAAATGATCG | |
| | AP98 | CGATCATTTCTTTGAGCTTGGAGGACAC TCCTTGTTGGCGGTAAAG | *X. szentirmaii* gDNA<br>3343 |
| | AP107 | CCGTACCGCCAGCAACGAATGACCACC CAGGGTAAAGAAGTTATCATGC | |
| | AP105 | TGGTCATTCGTTGCTGGCGG | *X. szentirmaii* gDNA<br>4199 |
| | AP106 | GGTGGCAGCAGCCTAGGTTAATTAATTA AATATTATTTACTATATTGTATTCCTCTGT ACC | |
| | AP61 | TTAATTAACCTAGGCTGCTGCCACCGCT GA | pCK_0434<br>3693 |
| | AP62 | GGAATTCCTCCTGTTAGCCCAAAAAAAC GG | |
| **pAP33** | AP63 | TTTTTGGGCTAACAGGAGGAATTCCATG AATATGACACGTAACCATACAT | *X. innexi* gDNA<br>3091 |
| | AP64.2 | TCCTCCAAGCTCAAAGAAATGATCG | |
| | AP100 | CGATCATTTCTTTGAGCTTGGAGGACAC TCGCTATTGATCGTGCAG | *X. miraniensis* gDNA<br>3271 |
| | AP104 | CCGTACCGCCAGCAACGAATGACCACC CAAGGCAAAGAAGCTG | |
| | AP105 | TGGTCATTCGTTGCTGGCGG | *X. szentirmaii* gDNA<br>4199 |
| | AP106 | GGTGGCAGCAGCCTAGGTTAATTAATTA AATATTATTTACTATATTGTATTCCTCTGT ACC | |
| | AP61 | TTAATTAACCTAGGCTGCTGCCACCGCT GA | pCK_0434<br>3693 |
| | AP62 | GGAATTCCTCCTGTTAGCCCAAAAAAAC GG | |
| **pAP34** | AP63 | TTTTTGGGCTAACAGGAGGAATTCCATG AATATGACACGTAACCATACAT | *X. innexi* gDNA<br>3091 |
| | AP64.2 | TCCTCCAAGCTCAAAGAAATGATCG | |
| | AP98 | CGATCATTTCTTTGAGCTTGGAGGACAC TCCTTGTTGGCGGTAAAG | *X. szentirmaii* gDNA<br>3343 |
| | AP107 | CCGTACCGCAAGCAACGAATGACCACC CAGGGTAAAGAAGTTATCATGC | |
| | AP108 | TGGTCATTCGTTGCTTGCGGTACG | *P. laumondii* TTO1 gDNA<br>4232 |
| | AP109 | GGTGGCAGCAGCCTAGGTTAATTAATCA CAGCGCCTCCGC | |

| Plasmids | Oligo-nucleotide | Sequence (5’ → 3’) | Template<br>Product size in bp |
|---|---|---|---|
| | AP61 | TTAATTAACCTAGGCTGCTGCCACCGCTGA | pCK_0434<br>3693 |
| | AP62 | GGAATTCCTCCTGTTAGCCCAAAAAAACGG | |
| **pAP35** | AP63 | TTTTTGGGCTAACAGGAGGAATTCCATGAATATGACACGTAACCATACAT | *X. innexi* gDNA<br>3091 |
| | AP64.2 | TCCTCCAAGCTCAAAGAAATGATCG | |
| | AP100 | CGATCATTTCTTTGAGCTTGGAGGACACTCGCTATTGATCGTGCAG | *X. miraniensis* gDNA<br>3271 |
| | AP110 | CCGTACCGCAAGCAACGAATGACCACCCAAGGCAAAGAAGCTG | |
| | AP108 | TGGTCATTCGTTGCTTGCGGTACG | *P. laumondii* TTO1 gDNA<br>4232 |
| | AP109 | GGTGGCAGCAGCCTAGGTTAATTAATCACAGCGCCTCCGC | |
| | AP61 | TTAATTAACCTAGGCTGCTGCCACCGCTGA | pCK_0434<br>3693 |
| | AP62 | GGAATTCCTCCTGTTAGCCCAAAAAAACGG | |
| **pAP36** | AP63 | TTTTTGGGCTAACAGGAGGAATTCCATGAATATGACACGTAACCATACAT | *X. innexi* gDNA<br>3091 |
| | AP64.2 | TCCTCCAAGCTCAAAGAAATGATCG | |
| | AP111 | CGATCATTTCTTTGAGCTTGGAGGACATTCCTTGATGGCGGT | *X. szentirmaii* gDNA<br>7433 |
| | AP106 | GGTGGCAGCAGCCTAGGTTAATTAATTAAATATTATTTACTATATTGTATTCCTCTGTACC | |
| | AP61 | TTAATTAACCTAGGCTGCTGCCACCGCTGA | pCK_0434<br>3693 |
| | AP62 | GGAATTCCTCCTGTTAGCCCAAAAAAACGG | |
| **pAP36b** | AP283 | GCCCTGGGTGGTCATTCGTT | pAP36<br>5168 |
| | AP46 | ATGGAGAAAAAAATCACTGGATATACCACC | |
| | AP45 | GGTGGTATATCCAGTGATTTTTTTCTCC | pAP36<br>9024 |
| | AP284 | AACGAATGACCACCCAGGGC | |
| **pAP37** | AP63 | TTTTTGGGCTAACAGGAGGAATTCCATGAATATGACACGTAACCATACAT | *X. indica* gDNA<br>3120 |
| | AP118 | ACCATGAGACCTTGCATTCGTTGGTCTCGTCCTCCAAGCTCAAAGAAATGATCG | |
| | AP115 | AGGACGAGACCAACGAATGCAAGGTCTCATGGTCATTCGTTGCTTGCGG | *P. laumondii* TTO1 gDNA<br>4261 |
| | AP109 | GGTGGCAGCAGCCTAGGTTAATTAATCACAGCGCCTCCGC | |

| Plasmids | Oligo-nucleotide | Sequence (5' → 3') | Template<br>Product size in bp |
|---|---|---|---|
| | AP61 | TTAATTAACCTAGGCTGCTGCCACCGCTGA | pCK_0434<br>3693 |
| | AP62 | GGAATTCCTCCTGTTAGCCCAAAAAAACGG | |
| **pAP38** | AP63 | TTTTTGGGCTAACAGGAGGAATTCCATGAATATGACACGTAACCATACAT | *X. indica* gDNA<br>3120 |
| | AP118 | ACCATGAGACCTTGCATTCGTTGGTCTCGTCCTCCAAGCTCAAAGAAATGATCG | |
| | AP117 | AGGACGAGACCAACGAATGCAAGGTCTCATGGTCATTCGTTGCTGGCG | *X. indica* gDNA<br>4228 |
| | AP106 | GGTGGCAGCAGCCTAGGTTAATTAATTAAATATTATTTACTATATTGTATTCCTCTGTACC | |
| | AP61 | TTAATTAACCTAGGCTGCTGCCACCGCTGA | pCK_0434<br>3693 |
| | AP62 | GGAATTCCTCCTGTTAGCCCAAAAAAACGG | |
| **pAP39** | AP121 | CACACAGGAAAGAAGGTCTCAAGGACACTCGCTATTGATCGTGCAGT | *X. miraniensis* gDNA<br>3271 |
| | AP122 | CAGTCACGACCTTTGGTCTCTACCACCCAAGGCAAAGAAGCTGTC | |
| | AP119 | TGGTAGAGACCAAAGGTCGTGACTGGGAAAACCC | pSEVA681<br>2351 |
| | AP120 | TCCTTGAGACCTTCTTTCCTGTGTGAAATTGTTATCCGCT | |
| **pAP40** | AP123 | CACACAGGAAAGAAGGTCTCAAGGACACTCCTTGTTGGCGGTAAAG | *X. szentirmaii* gDNA<br>3343 |
| | AP124 | CAGTCACGACCTTTGGTCTCTACCACCCAGGGTAAAGAAGTTATCATGC | |
| | AP119 | TGGTAGAGACCAAAGGTCGTGACTGGGAAAACCC | pSEVA681<br>2351 |
| | AP120 | TCCTTGAGACCTTCTTTCCTGTGTGAAATTGTTATCCGCT | |
| **pAP41** | AP125 | CACACAGGAAAGAAGGTCTCAAGGACATTCCTTGATGGCGGT | *X. szentirmaii* gDNA<br>3259 |
| | AP126 | CAGTCACGACCTTTGGTCTCTACCACCCAGGGCAAAGAAATTATC | |
| | AP119 | TGGTAGAGACCAAAGGTCGTGACTGGGAAAACCC | pSEVA681<br>2351 |
| | AP120 | TCCTTGAGACCTTCTTTCCTGTGTGAAATTGTTATCCGCT | |
| **pAP42** | AP128 | CACACAGGAAAGAAGGTCTCAAGGACATTCTCTGTTAGCGGTACG | *X. nematophila* gDNA<br>3229 |

| Plasmids | Oligo-nucleotide | Sequence (5’ → 3’) | Template<br>Product size in bp |
|---|---|---|---|
| | AP129 | CAGTCACGACCTTTGGTCTCTACCACCC AAAGCAAAGAAACTGTCA | |
| | AP119 | TGGTAGAGACCAAAGGTCGTGACTGGG AAAACCC | pSEVA681<br>2351 |
| | AP120 | TCCTTGAGACCTTCTTTCCTGTGTGAAAT TGTTATCCGCT | |
| **pAP43** | AP130 | CACACAGGAAAGAAGGTCTCAAGGACAT TCATTGCTTGCTGTCCA | *X. doucetiae* gDNA<br>3265 |
| | AP131 | CAGTCACGACCTTTGGTCTCTACCACCC AATTCAAAGAAATGATCATGG | |
| | AP119 | TGGTAGAGACCAAAGGTCGTGACTGGG AAAACCC | pSEVA681<br>2351 |
| | AP120 | TCCTTGAGACCTTCTTTCCTGTGTGAAAT TGTTATCCGCT | |
| **pAP44** | AP132 | CACACAGGAAAGAAGGTCTCAAGGACA CTCACTGCTGGCTGTC | *X. doucetiae* gDNA<br>3217 |
| | AP133 | CAGTCACGACCTTTGGTCTCTACCACCA AGCTCAAAGAAATGGTCAT | |
| | AP119 | TGGTAGAGACCAAAGGTCGTGACTGGG AAAACCC | pSEVA681<br>2351 |
| | AP120 | TCCTTGAGACCTTCTTTCCTGTGTGAAAT TGTTATCCGCT | |
| **pAP45** | AP121 | CACACAGGAAAGAAGGTCTCAAGGACA CTCGCTATTGATCGTGCAGT | pAP39<br>3262 |
| | AP154 | CCTTTGGTCTCTGCCGCCCAAGGCAAA G | |
| | AP150 | CGGCAGAGACCAAAGGTCGTGACTG | pSEVA681<br>2351 |
| | AP149 | TCCTTGAGACCTTCTTTCCTGTGTG | |
| **pAP46** | AP123 | CACACAGGAAAGAAGGTCTCAAGGACA CTCCTTGTTGGCGGTAAAG | pAP40<br>3332 |
| | AP162 | TTTGGTCTCTGCCGCCCAGGGTAA | |
| | AP150 | CGGCAGAGACCAAAGGTCGTGACTG | pSEVA681<br>2351 |
| | AP149 | TCCTTGAGACCTTCTTTCCTGTGTG | |
| **pAP47** | AP125 | CACACAGGAAAGAAGGTCTCAAGGACAT TCCTTGATGGCGGT | pAP41<br>3248 |
| | AP164 | TTTGGTCTCTGCCGCCCAGGGCAA | |
| | AP150 | CGGCAGAGACCAAAGGTCGTGACTG | pSEVA681<br>2351 |
| | AP149 | TCCTTGAGACCTTCTTTCCTGTGTG | |
| **pAP48** | AP128 | CACACAGGAAAGAAGGTCTCAAGGACAT TCTCTGTTAGCGGTACG | pAP42<br>3218 |
| | AP166 | TTTGGTCTCTGCCGCCCAAAGCAA | |
| | AP150 | CGGCAGAGACCAAAGGTCGTGACTG | pSEVA681<br>2351 |
| | AP149 | TCCTTGAGACCTTCTTTCCTGTGTG | |

| Plasmids | Oligo-nucleotide | Sequence (5' → 3') | Template<br>Product size in bp |
|---|---|---|---|
| **pAP49** | AP130 | CACACAGGAAAGAAGGTCTCAAGGACAT TCATTGCTTGCTGTCCA | pAP43<br>3254 |
| | AP168 | TTTGGTCTCTGCCGCCGAGTTCAA | |
| | AP150 | CGGCAGAGACCAAAGGTCGTGACTG | pSEVA681<br>2351 |
| | AP149 | TCCTTGAGACCTTCTTTCCTGTGTG | |
| **pAP50** | AP132 | CACACAGGAAAGAAGGTCTCAAGGACA CTCACTGCTGGCTGTC | pAP44<br>3217 |
| | AP170 | TTTGGTCTCTGCCGCCAAGCTCAA | |
| | AP150 | CGGCAGAGACCAAAGGTCGTGACTG | pSEVA681<br>2351 |
| | AP149 | TCCTTGAGACCTTCTTTCCTGTGTG | |
| **pAP51** | AP155 | AAGAAGGTCTCACGGCCACTCGCTATTG | pAP39<br>3262 |
| | AP122 | CAGTCACGACCTTTGGTCTCTACCACCC AAGGCAAAGAAGCTGTC | |
| | AP151 | TGGTAGAGACCAAAGGTCGTGACTG | pSEVA681<br>2351 |
| | AP156 | CAATAGCGAGTGGCCGTGAGACCTTCTT | |
| **pAP52** | AP138 | CACACAGGAAAGAAGGTCTCACGGC | pAP40<br>3343 |
| | AP152 | CAGTCACGACCTTTGGTCTCTACCA | |
| | AP151 | TGGTAGAGACCAAAGGTCGTGACTG | pAP40<br>2351 |
| | AP139 | GCCGTGAGACCTTCTTTCCTGTGTG | |
| **pAP53** | AP173 | GAAGGTCTCACGGCCATTCCTTGA | pAP41<br>3248 |
| | AP152 | CAGTCACGACCTTTGGTCTCTACCA | |
| | AP151 | TGGTAGAGACCAAAGGTCGTGACTG | pAP41<br>2361 |
| | AP174 | TCAAGGAATGGCCGTGAGACCTTC | |
| **pAP54** | AP175 | GAAGGTCTCACGGCCATTCTCTGT | pAP42<br>3218 |
| | AP152 | CAGTCACGACCTTTGGTCTCTACCA | |
| | AP151 | TGGTAGAGACCAAAGGTCGTGACTG | pAP42<br>2361 |
| | AP176 | ACAGAGAATGGCCGTGAGACCTTC | |
| **pAP55** | AP177 | GAAGGTCTCACGGCCACTCATTGC | pAP43<br>3254 |
| | AP152 | CAGTCACGACCTTTGGTCTCTACCA | |
| | AP151 | TGGTAGAGACCAAAGGTCGTGACTG | pAP43<br>2361 |
| | AP178 | GCAATGAGTGGCCGTGAGACCTTC | |
| **pAP56** | AP179 | GAAGGTCTCACGGCCACTCACTGC | pAP44<br>3206 |
| | AP152 | CAGTCACGACCTTTGGTCTCTACCA | |
| | AP151 | TGGTAGAGACCAAAGGTCGTGACTG | pAP44<br>2361 |
| | AP180 | GCAGTGAGTGGCCGTGAGACCTTC | |
| **pAP89** | AP226 | CACACAGGAAAGAAGGTCTCAAGGAGA CTCCATTACCAGTATCCAATTAGT | *X. innexi* gDNA<br>819 |
| | AP229 | GGGGTCTTGAGTCCACCCGATTG | |
| | AP228 | CAATCGGGTGGACTCAAGACCCC | |

| Plasmids | Oligo-nucleotide | Sequence (5’ → 3’) | Template<br>Product size in bp |
|---|---|---|---|
| | AP227 | CAGTCACGACCTTTGGTCTCTGCCGCC GATACGGAAGAAATTATCTTCGA | *X. innexi* gDNA<br>3453 |
| | AP134 | CGGCAGAGACCAAAGGTCGTGACTG | pAP45<br>2351 |
| | AP149 | TCCTTGAGACCTTCTTTCCTGTGTG | |
| **pAP92** | AP234 | CACACAGGAAAGAAGGTCTCACGGCGA CTCCATTACCAGTATCCAATTAGT | pAP89<br>4249 |
| | AP235 | CAGTCACGACCTTTGGTCTCTACCACCG ATACGGAAGAAATTATCTTCGA | |
| | AP151 | TGGTAGAGACCAAAGGTCGTGACTG | pAP51<br>2351 |
| | AP139 | GCCGTGAGACCTTCTTTCCTGTGTG | |
| **pAP95** | AP242 | CACACAGGAAAGAAGGTCTCAAGGACA CTCATTGATGGTGGTCCG | *X. innexi* gDNA<br>3489 |
| | AP243 | CAGTCACGACCTTTGGTCTCTGCCGCCC AGTTCAAAGAAATGGTCATG | |
| | AP134 | CGGCAGAGACCAAAGGTCGTGACTG | pAP45<br>2351 |
| | AP149 | TCCTTGAGACCTTCTTTCCTGTGTG | |
| **pAP96** | AP244 | CACACAGGAAAGAAGGTCTCAAGGACAT TCCCTGCTGGCTATCC | *X. beddingii* gDNA<br>3286 |
| | AP245 | CAGTCACGACCTTTGGTCTCTGCCGCC GAGTTCAAAGAAATGGTCATG | |
| | AP134 | CGGCAGAGACCAAAGGTCGTGACTG | pAP45<br>2351 |
| | AP149 | TCCTTGAGACCTTCTTTCCTGTGTG | |
| **pAP98** | AP246 | CACACAGGAAAGAAGGTCTCACGGCCA CTCATTGATGGTGGTCCG | *X. innexi* gDNA<br>3489 |
| | AP247 | CAGTCACGACCTTTGGTCTCTACCACCC AGTTCAAAGAAATGGTCATG | |
| | AP151 | TGGTAGAGACCAAAGGTCGTGACTG | pAP51<br>2351 |
| | AP139 | GCCGTGAGACCTTCTTTCCTGTGTG | |
| **pAP99** | AP248 | CACACAGGAAAGAAGGTCTCACGGCCA TTCCCTGCTGGCTATCC | *X. beddingii* gDNA<br>3286 |
| | AP216 | CAGTCACGACCTTTGGTCTCTACCACCG AGTTCAAAGAAATGGTCATG | |
| | AP151 | TGGTAGAGACCAAAGGTCGTGACTG | pAP51<br>2351 |
| | AP139 | GCCGTGAGACCTTCTTTCCTGTGTG | |
| **pAP106** | AP268 | CGAGACCAACGAATGCAAGGTCTCAAG GACACTCACTGCTGGCTGTC | pAP37_44<br>7428 |
| | AP109 | GGTGGCAGCAGCCTAGGTTAATTAATCA CAGCGCCTCCGC | |
| | AP61 | TTAATTAACCTAGGCTGCTGCCACCGCT GA | pAP37_44<br>3721 |
| | AP269 | TGAGACCTTGCATTCGTTGGTCTCGCAT GGAATTCCTCCTGTTAGCCCAAAA | |

| Plasmids | Oligo-nucleotide | Sequence (5' → 3') | Template<br>Product size in bp |
|---|---|---|---|
| **pAP107** | AP271 | CACACAGGAAAGAAGGTCTCACATGAAT ATGACACGTAACCATACATCCTC | pAP37_44<br>3107 |
| | AP272 | GTCACGACCTTTGGTCTCTTCCTCCAAG CTCAAAGAAATGATCG | |
| | AP273 | AGGAAGAGACCAAAGGTCGTGAC | pAP39<br>2351 |
| | AP274 | CATGTGAGACCTTCTTTCCTGTGTG | |
| **pAP108** | AP275 | CACACAGGAAAGAAGGTCTCACATGCCT ATGTCATGCAATGGTATTAACAACG | *X. doucetiae* gDNA<br>3236 |
| | AP276 | GTCACGACCTTTGGTCTCTTCCTCCAAG CTCAAAGAAATGGTCATGG | |
| | AP273 | AGGAAGAGACCAAAGGTCGTGAC | pAP39<br>2351 |
| | AP274 | CATGTGAGACCTTCTTTCCTGTGTG | |
| **pAP109** | AP277 | CACACAGGAAAGAAGGTCTCACATGAAA AATGATAAGGTGATGACTCTGCCAAC | *X. miraniensis* gDNA<br>2981 |
| | AP278 | GTCACGACCTTTGGTCTCTTCCTCCGAG CGTAAAGAAGTTATCAAACC | |
| | AP273 | AGGAAGAGACCAAAGGTCGTGAC | pAP39<br>2351 |
| | AP274 | CATGTGAGACCTTCTTTCCTGTGTG | |
| **pAP110** | AP279 | ACAGGAAAGAAGGTCTCACATGCCTATG TCATGCAATAGTAGCAATAATATTAAATT TCC | *X. ishibashii* gDNA<br>3104 |
| | AP280 | GTCACGACCTTTGGTCTCTTCCTCCGAG TTCGAAGAAATGATCGTAACG | |
| | AP273 | AGGAAGAGACCAAAGGTCGTGAC | pAP39<br>2351 |
| | AP274 | CATGTGAGACCTTCTTTCCTGTGTG | |
| **pAP111** | AP285 | TTTTTGGGCTAACAGGAGGAATTCCATG ATAAGGTCAGAAGACATGAACCTC | *X. doucetiae* gDNA<br>8180 |
| | AP286 | CCGACGGAAAATCCACCTGG | |
| | AP287 | CCAGGTGGATTTTCCGTCGG | *X. doucetiae* gDNA<br>8373 |
| | AP288 | GGTGGCAGCAGCCTAGGTTAATTAATCA ACTAATCACAGCCCAAGCC | |
| | AP61 | TTAATTAACCTAGGCTGCTGCCACCGCT GA | pAP37<br>3693 |
| | AP62 | GGAATTCCTCCTGTTAGCCCAAAAAAAC GG | |
| **pAP112** | AP289 | CACGAGTTGGGACTCTTCTATCG | pAP111<br>6129 |
| | AP292 | CATTAATACAGGCCTCTCAAAGAGTG | |
| | AP291 | CACTCTTTGAGAGGCCTGTATTAATG | pAP111<br>4516 |
| | AP294 | GCCAAATCGGTTTCCGTTTCAATC | |
| | AP293 | GATTGAAACGGAAACCGATTTGGC | pAP111<br>3364 |
| | AP296 | CGGAAAGAGTCCGGCTTTCA | |
| | AP295 | TGAAAGCCGGACTCTTTCCG | |

| Plasmids | Oligo-nucleotide | Sequence (5’ → 3’) | Template Product size in bp |
|---|---|---|---|
| | AP298 | CCCTGTTCAGACACCAGAATGG | pAP111 1001 |
| | AP297 | CCATTCTGGTGTCTGAACAGGG | pAP111 5281 |
| | AP290 | CGATAGAAGAGTCCCAACTCGTG | |
| **pAP113** | AP299 | CACACAGGAAAGAAGGTCTCAAGGACATTCACTGATGATCGTCAGCCTG | *X. doucetiae* gDNA 3400 |
| | AP300 | CAGTCACGACCTTTGGTCTCTACCACCAAGTTCAAAGAAGTGATCATGGC | |
| | AP119 | TGGTAGAGACCAAAGGTCGTGACTGGGAAAACCC | pAP39 2336 |
| | AP120 | TCCTTGAGACCTTCTTTCCTGTGTGAAATTGTTATCCGCT | |
| **pAP114** | AP301 | CACACAGGAAAGAAGGTCTCAAGGACACTCCCTGCTGGCTGTC | *X. doucetiae* gDNA 121 |
| | AP292 | CATTAATACAGGCCTCTCAAAGAGTG | |
| | AP291 | CACTCTTTGAGAGGCCTGTATTAATG | *X. doucetiae* gDNA 3182 |
| | AP302 | CAGTCACGACCTTTGGTCTCTACCACCAAGTTCAAAGAAATGGTCATGGC | |
| | AP119 | TGGTAGAGACCAAAGGTCGTGACTGGGAAAACCC | pAP39 2336 |
| | AP120 | TCCTTGAGACCTTCTTTCCTGTGTGAAATTGTTATCCGCT | |
| **pAP115** | AP303 | CACACAGGAAAGAAGGTCTCAAGGACATTCACTGATGATCGTCAGCCT | pAP113 3400 |
| | AP304 | CAGTCACGACCTTTGGTCTCTGCCGCCAAGTTCAAAGAAGTGATCATGGC | |
| | AP150 | CGGCAGAGACCAAAGGTCGTGACTG | pAP113 2351 |
| | AP149 | TCCTTGAGACCTTCTTTCCTGTGTG | |
| **pAP116** | AP305 | CACACAGGAAAGAAGGTCTCAAGGACACTCCCTGCTGGCTGTC | pAP114 3277 |
| | AP306 | CAGTCACGACCTTTGGTCTCTGCCGCCAAGTTCAAAGAAGTGATCATGGC | |
| | AP150 | CGGCAGAGACCAAAGGTCGTGACTG | pAP114 2351 |
| | AP149 | TCCTTGAGACCTTCTTTCCTGTGTG | |
| **pAP117** | AP307 | CACACAGGAAAGAAGGTCTCACGGCCATTCACTGATGATCGTCAGCCT | pAP113 3400 |
| | AP308 | CAGTCACGACCTTTGGTCTCTACCACCAAGTTCAAAGAAGTGATCATGGC | |
| | AP151 | TGGTAGAGACCAAAGGTCGTGACTG | pAP113 2351 |
| | AP139 | GCCGTGAGACCTTCTTTCCTGTGTG | |
| **pAP118** | AP309 | CACACAGGAAAGAAGGTCTCACGGCCACTCCCTGCTGGCTGTC | pAP114 3277 |

| Plasmids | Oligo-nucleotide | Sequence (5' → 3') | Template Product size in bp |
|---|---|---|---|
| | AP310 | CAGTCACGACCTTTGGTCTCTACCACCAAGTTCAAAGAAATGGTCATGGC | |
| | AP151 | TGGTAGAGACCAAAGGTCGTGACTG | pAP114 2351 |
| | AP139 | GCCGTGAGACCTTCTTTCCTGTGTG | |
| **pAP119** | AP312 | AGGACGAGACCAACGAATGCAAGGTCTCATGGTCACTCGCTGCTGGCCGTC | pAP112 7662 |
| | AP46 | ATGGAGAAAAAAATCACTGGATATACCACC | |
| | AP45 | GGTGGTATATCCAGTGATTTTTTTCTCC | pAP112 6021 |
| | AP312 | ACCATGAGACCTTGCATTCGTTGGTCTCGTCCTCCAAGCTCAAAGAAATGGTCATGCC | |
| **pAP128** | AP351 | TTTTTGGGCTAACAGGAGGAATTCCATGCCTATGTCATGCAATGGTATTAAC | *X. doucetiae* gDNA 14520 |
| | AP323 | GTGGCATTGAAATCGACCAGTATTTG | |
| | AP324 | CAAATACTGGTCGATTTCAATGCCAC | *X. doucetiae* gDNA 15183 |
| | AP352 | AGCGGTGGCAGCAGCCTAGGTTAATTCATTGATGACTATCTCCGCCTAAC | |
| | AP353 | ATTAACCTAGGCTGCTGCCAC | pCOLA_ara/tacI 3108 |
| | AP68 | GGAATTCCTCCTGTTAGCCCAAAAA | |

**Table S5.** Primer used in this work for the qPCR analysis of the tera-modular NRPS library.

| Oligonucleotide | Sequence (5' → 3') | qPCR targeted template |
|---|---|---|
| SLo5558 | ACTTGCACAACATCTGGCTG | pAP37 backbone fw. |
| SLo5559 | CAGAATCCCATCCTGCAACG | pAP37 backbone rv. |
| SLo5560 | TCAACGGCAACATTTCAGGG | pAP45 rv. |
| SLo5561 | TTTGGGCCAATTCTGACAGC | pAP46 rv. |
| SLo5562 | AGGGGACAACGGGTAAATGT | pAP47 rv. |
| SLo5563 | CAGGGCTGTCGGGATATGAT | pAP48 rv. |
| SLo5564 | ACGGATATGGGCCATGAGTT | pAP49 rv. |
| SLo5565 | ATGAGAGTGGCAAGGGTTGA | pAP50 rv. |
| SLo5566 | CATCTGCGTGTTCACCTGAG | pAP51 fw. |
| SLo5567 | ACAAGGGGAGACAGAAACCA | pAP52 fw. |
| SLo5568 | TTGCCCGTCAGATTTATGCG | pAP53 fw. |
| SLo5569 | CATTAGCGGCTATCTGGTGC | pAP54 fw. |
| SLo5570 | ACTCACCATCTCGCTGAACA | pAP55 fw. |
| SLo5571 | CGTTTGGTCGCTTATCTGCA | pAP56 fw. |

| Oligonucleotide | Sequence (5' → 3') | qPCR targeted template |
|---|---|---|
| SLo5912 | TCAGTAGCTTTGCCAGACGA | pAP89 rv. |
| SLo5913 | AACAAAGACAGCTACACCGC | pAP92 fw. |
| SLo5914 | ATGAGTTCGGCCTGTTCTGT | pAP95 rv. |
| SLo5915 | TGGCTCGCGTCTATGAATCT | pAP98 fw. |

**Table S6.** Acceptor and donor plasmids used for the GGA assembled plasmids created and used in this work.

| Assembled Plasmid | Acceptor Plasmid | Donor Plasmids |
|---|---|---|
| pAP37_39 | pAP37 | pAP39 |
| pAP37_40 | pAP37 | pAP40 |
| pAP37_41 | pAP37 | pAP41 |
| pAP37_42 | pAP37 | pAP42 |
| pAP37_43 | pAP37 | pAP43 |
| pAP37_44 | pAP37 | pAP44 |
| pAP38_39 | pAP38 | pAP39 |
| pAP38_40 | pAP38 | pAP40 |
| pAP38_41 | pAP38 | pAP41 |
| pAP38_42 | pAP38 | pAP42 |
| pAP38_43 | pAP38 | pAP43 |
| pAP38_44 | pAP38 | pAP44 |
| pAP37_45_51 | pAP37 | pAP45, pAP51 |
| pAP37_45_52 | pAP37 | pAP45, pAP52 |
| pAP37_45_53 | pAP37 | pAP45, pAP53 |
| pAP37_45_54 | pAP37 | pAP45, pAP54 |
| pAP37_45_55 | pAP37 | pAP45, pAP55 |
| pAP37_45_56 | pAP37 | pAP45, pAP56 |
| pAP37_46_51 | pAP37 | pAP46, pAP51 |
| pAP37_46_52 | pAP37 | pAP46, pAP52 |
| pAP37_46_53 | pAP37 | pAP46, pAP53 |
| pAP37_46_54 | pAP37 | pAP46, pAP54 |
| pAP37_46_55 | pAP37 | pAP46, pAP55 |
| pAP37_46_56 | pAP37 | pAP46, pAP56 |
| pAP37_47_51 | pAP37 | pAP47, pAP51 |
| pAP37_47_52 | pAP37 | pAP47, pAP52 |
| pAP37_47_53 | pAP37 | pAP47, pAP53 |
| pAP37_47_54 | pAP37 | pAP47, pAP54 |
| pAP37_47_55 | pAP37 | pAP47, pAP55 |
| pAP37_47_56 | pAP37 | pAP47, pAP56 |

| Assembled Plasmid | Acceptor Plasmid | Donor Plasmids |
|---|---|---|
| pAP37_48_51 | pAP37 | pAP48, pAP51 |
| pAP37_48_52 | pAP37 | pAP48, pAP52 |
| pAP37_48_53 | pAP37 | pAP48, pAP53 |
| pAP37_48_54 | pAP37 | pAP48, pAP54 |
| pAP37_48_55 | pAP37 | pAP48, pAP55 |
| pAP37_48_56 | pAP37 | pAP48, pAP56 |
| pAP37_49_51 | pAP37 | pAP49, pAP51 |
| pAP37_49_52 | pAP37 | pAP49, pAP52 |
| pAP37_49_53 | pAP37 | pAP49, pAP53 |
| pAP37_49_54 | pAP37 | pAP49, pAP54 |
| pAP37_49_55 | pAP37 | pAP49, pAP55 |
| pAP37_49_56 | pAP37 | pAP49, pAP56 |
| pAP37_50_51 | pAP37 | pAP50, pAP51 |
| pAP37_50_52 | pAP37 | pAP50, pAP52 |
| pAP37_50_53 | pAP37 | pAP50, pAP53 |
| pAP37_50_54 | pAP37 | pAP50, pAP54 |
| pAP37_50_55 | pAP37 | pAP50, pAP55 |
| pAP37_50_56 | pAP37 | pAP50, pAP56 |
| pAP37_95_51 | pAP37 | pAP95, pAP51 |
| pAP37_95_52 | pAP37 | pAP95, pAP52 |
| pAP37_95_53 | pAP37 | pAP95, pAP53 |
| pAP37_95_54 | pAP37 | pAP95, pAP54 |
| pAP37_95_55 | pAP37 | pAP95, pAP55 |
| pAP37_95_56 | pAP37 | pAP95, pAP56 |
| pAP37_96_51 | pAP37 | pAP96, pAP51 |
| pAP37_96_52 | pAP37 | pAP96, pAP52 |
| pAP37_96_53 | pAP37 | pAP96, pAP53 |
| pAP37_96_54 | pAP37 | pAP96, pAP54 |
| pAP37_96_55 | pAP37 | pAP96, pAP55 |
| pAP37_96_56 | pAP37 | pAP96, pAP56 |
| pAP37_45_98 | pAP37 | pAP45, pAP98 |
| pAP37_46_98 | pAP37 | pAP46, pAP98 |
| pAP37_47_98 | pAP37 | pAP47, pAP98 |
| pAP37_48_98 | pAP37 | pAP48, pAP98 |
| pAP37_49_98 | pAP37 | pAP49, pAP98 |
| pAP37_50_98 | pAP37 | pAP50, pAP98 |
| pAP37_45_99 | pAP37 | pAP45, pAP99 |
| pAP37_46_99 | pAP37 | pAP46, pAP99 |
| pAP37_47_99 | pAP37 | pAP47, pAP99 |
| pAP37_48_99 | pAP37 | pAP48, pAP99 |

| Assembled Plasmid | Acceptor Plasmid | Donor Plasmids |
|---|---|---|
| pAP37_49_99 | pAP37 | pAP49, pAP99 |
| pAP37_50_99 | pAP37 | pAP50, pAP99 |
| pAP37_95_98 | pAP37 | pAP95, pAP98 |
| pAP37_95_99 | pAP37 | pAP95, pAP99 |
| pAP37_96_98 | pAP37 | pAP96, pAP98 |
| pAP37_96_99 | pAP37 | pAP96, pAP99 |
| pAP37_89_51 | pAP37 | pAP89, pAP51 |
| pAP37_89_52 | pAP37 | pAP89, pAP52 |
| pAP37_89_53 | pAP37 | pAP89, pAP53 |
| pAP37_89_54 | pAP37 | pAP89, pAP54 |
| pAP37_89_55 | pAP37 | pAP89, pAP55 |
| pAP37_89_56 | pAP37 | pAP89, pAP56 |
| pAP37_45_92 | pAP37 | pAP45, pAP92 |
| pAP37_46_92 | pAP37 | pAP46, pAP92 |
| pAP37_47_92 | pAP37 | pAP47, pAP92 |
| pAP37_48_92 | pAP37 | pAP48, pAP92 |
| pAP37_49_92 | pAP37 | pAP49, pAP92 |
| pAP37_50_92 | pAP37 | pAP50, pAP92 |
| pAP37_89_92 | pAP37 | pAP89, pAP92 |
| pAP37_89_98 | pAP37 | pAP89, pAP98 |
| pAP37_89_99 | pAP37 | pAP89, pAP99 |
| pAP37_95_92 | pAP37 | pAP95, pAP92 |
| pAP37_96_92 | pAP37 | pAP96, pAP92 |
| pAP106_107 | pAP106 | pAP107 |
| pAP106_108 | pAP106 | pAP108 |
| pAP106_109 | pAP106 | pAP109 |
| pAP106_110 | pAP106 | pAP110 |
| pAP119_113 | pAP119 | pAP113 |
| pAP119_114 | pAP119 | pAP114 |
| pAP119_115_117 | pAP119 | pAP115, pAP117 |
| pAP119_115_118 | pAP119 | pAP115, pAP118 |
| pAP119_116_117 | pAP119 | pAP116, pAP117 |
| pAP119_116_118 | pAP119 | pAP116, pAP118 |
| pAP119_39 | pAP119 | pAP39 |
| pAP119_40 | pAP119 | pAP40 |
| pAP119_41 | pAP119 | pAP41 |
| pAP119_42 | pAP119 | pAP42 |
| pAP119_43 | pAP119 | pAP43 |
| pAP119_44 | pAP119 | pAP44 |
| pAP119_45_51 | pAP119 | pAP45, pAP51 |

| Assembled Plasmid | Acceptor Plasmid | Donor Plasmids |
|---|---|---|
| pAP119_45_52 | pAP119 | pAP45, pAP52 |
| pAP119_45_53 | pAP119 | pAP45, pAP53 |
| pAP119_45_54 | pAP119 | pAP45, pAP54 |
| pAP119_45_55 | pAP119 | pAP45, pAP55 |
| pAP119_45_56 | pAP119 | pAP45, pAP56 |
| pAP119_46_51 | pAP119 | pAP46, pAP51 |
| pAP119_46_52 | pAP119 | pAP46, pAP52 |
| pAP119_46_53 | pAP119 | pAP46, pAP53 |
| pAP119_46_54 | pAP119 | pAP46, pAP54 |
| pAP119_46_55 | pAP119 | pAP46, pAP55 |
| pAP119_46_56 | pAP119 | pAP46, pAP56 |
| pAP119_47_51 | pAP119 | pAP47, pAP51 |
| pAP119_47_52 | pAP119 | pAP47, pAP52 |
| pAP119_47_53 | pAP119 | pAP47, pAP53 |
| pAP119_47_54 | pAP119 | pAP47, pAP54 |
| pAP119_47_55 | pAP119 | pAP47, pAP55 |
| pAP119_47_56 | pAP119 | pAP47, pAP56 |
| pAP119_48_51 | pAP119 | pAP48, pAP51 |
| pAP119_48_52 | pAP119 | pAP48, pAP52 |
| pAP119_48_53 | pAP119 | pAP48, pAP53 |
| pAP119_48_54 | pAP119 | pAP48, pAP54 |
| pAP119_48_55 | pAP119 | pAP48, pAP55 |
| pAP119_48_56 | pAP119 | pAP48, pAP56 |
| pAP119_49_51 | pAP119 | pAP49, pAP51 |
| pAP119_49_52 | pAP119 | pAP49, pAP52 |
| pAP119_49_53 | pAP119 | pAP49, pAP53 |
| pAP119_49_54 | pAP119 | pAP49, pAP54 |
| pAP119_49_55 | pAP119 | pAP49, pAP55 |
| pAP119_49_56 | pAP119 | pAP49, pAP56 |
| pAP119_50_51 | pAP119 | pAP50, pAP51 |
| pAP119_50_52 | pAP119 | pAP50, pAP52 |
| pAP119_50_53 | pAP119 | pAP50, pAP53 |
| pAP119_50_54 | pAP119 | pAP50, pAP54 |
| pAP119_50_55 | pAP119 | pAP50, pAP55 |
| pAP119_50_56 | pAP119 | pAP50, pAP56 |

**Table S7. List of all compounds produced by NRPS related to the hybrid system (NRPS-A1, NRPS-A2 and NRPS-A3) for the first and second donor libraries and the starter library.** Expected compounds that were not detected are marked as ND (not detected). Small italic letters indicate D and small italic bold letters indicate D-*allo*-amino acids, respectively.

| NRPS- | Compound | Peptide sequence | MS calculated $[M+H]^+$ | MS detected $[M+H]^+$ | Ion Formula | Δppm |
|---|---|---|---|---|---|---|
| **1** | | C14-*q*TW | 644.402325 | ND | | |
| **2** | | C14-*q*QW | 671.4127 | ND | | |
| **3** | | C14-*q*__*t*__L | 571.4066 | ND | | |
| **4** | **1** | C14-*qq*L | 598.4174 | 598.4166 | $C_{30}H_{56}N_5O_7$ | 1.3 |
| **5** | **2** | C14-*q*YW | 706.4174 | 706.4179 | $C_{39}H_{56}N_5O_7$ | 0.7 |
| | **3** | C12-*q*YW | 678.3861 | 678.3858 | $C_{37}H_{52}N_5O_7$ | 0.3 |
| | **4** | C14-*q*YF | 667.4066 | 667.4065 | $C_{37}H_{55}N_4O_7$ | 0.1 |
| | **5** | C12-*q*YF | 639.3752 | 639.3747 | $C_{35}H_{51}N_4O_7$ | 0.7 |
| | **6** | C14-*q*FW | 690.4225 | 690.4221 | $C_{39}H_{56}N_5O_6$ | 0.5 |
| | **7** | C14-*q*FF | 651.4116 | 651.4114 | $C_{37}H_{55}N_4O_6$ | 0.4 |
| **A2_D1** | **1** | C14-*qq*L | 598.4174 | 598.4169 | $C_{30}H_{56}N_5O_7$ | 0.9 |
| **A2_D2** | **8** | C14-*q*__*t*__L | 571.4066 | 571.4054 | $C_{29}H_{54}N_4O_7$ | 2.0 |
| | **9** | C12-*q*__*t*__L | 543.3752 | 543.3744 | $C_{27}H_{51}N_4O_7$ | 0.9 |
| **A2_D3** | **10** | C14-*qq*L | 633.4222 | 633.4206 | $C_{34}H_{57}N_4O_7$ | 2.5 |
| **A2_D4** | | C14-Q*v*L | 569.4273 | ND | | |
| | **11** | C14-*q*L | 470.3589 | 470.3587 | $C_{25}H_{48}N_3O_5$ | 0.1 |
| **A2_D5** | **12** | C14-Q*l*L | 583.4429 | 583.4429 | $C_{31}H_{59}N_4O_6$ | 0 |
| **A2_D6** | **13** | C14-Q*a*L | 541.396 | 541.3961 | $C_{28}H_{53}N_4O_6$ | 0.3 |

| NRPS- | Compound | Peptide sequence | MS calculated [M+H]$^+$ | MS detected [M+H]$^+$ | Ion Formula | Δppm |
|---|---|---|---|---|---|---|
| **A1_D1** | | C14-*q*QW | 671.4127 | ND | | |
| **A1_D2** | | C14-*q*TW | 644.4018 | ND | | |
| **A1_D3** | **2** | C14-*q*YW | 706.4174 | 706.4175 | $C_{39}H_{56}N_5O_7$ | 0.1 |
| | **3** | C12-*q*YW | 678.3861 | 678.3858 | $C_{37}H_{52}N_5O_7$ | 0.3 |
| | **4** | C14-*q*YF | 667.4066 | 667.4065 | $C_{37}H_{55}N_4O_7$ | 0.1 |
| | **5** | C12-*q*YF | 639.3752 | 639.3745 | $C_{35}H_{51}N_4O_7$ | 1.2 |
| | **6** | C14-*q*FW | 690.4225 | 690.4219 | $C_{39}H_{56}N_5O_6$ | 0.9 |
| | **7** | C14-*q*FF | 651.4116 | 651.411 | $C_{37}H_{55}N_4O_6$ | 1 |
| **A1_D4** | | C14-QVW | 642.4225 | ND | | |
| **A1_D5** | | C14-QLW | 656.4382 | ND | | |
| **A1_D6** | | C14-QAW | 614.3912 | ND | | |
| **A2_D7_D16** | **14** | C14-*qqq*L | 726.476 | 726.4761 | $C_{35}H_{64}N_7O_9$ | 0.1 |
| **A2_D7_D17** | **15** | C14-*qq**t***L | 699.4651 | 699.4642 | $C_{35}H_{64}N_7O_{10}$ | 1.2 |
| | **8** | C14-*q**t***L | 571.4066 | 571.4059 | $C_{29}H_{55}N_4O_7$ | 1.1 |
| **A2_D7_D18** | | C14-*qqy*L | 761.4808 | ND | | |
| **A2_D7_D19** | **16** | C14-*q*Q*v*L | 697.4859 | 697.4862 | $C_{35}H_{65}N_6O_8$ | 0.5 |
| **A2_D7_D20** | | C14-*q*Q*l*L | 711.5015 | ND | | |
| | **13** | C14-Q-*a*L | 541.396 | 541.3957 | $C_{28}H_{53}N_4O_6$ | 0.5 |
| **A2_D7_D21** | **17** | C14-*q*Q*a*L | 669.4546 | 669.4539 | $C_{33}H_{61}N_6O_8$ | 0.9 |
| | **13** | C14-Q*a*L | 541.396 | 541.3958 | $C_{28}H_{53}N_4O_6$ | 0.4 |
| **A2_D8_D16** | | C14-*q**t**q*L | 699.4651 | ND | | |
| | **1** | C14-*qq*L | 598.4174 | 598.4165 | $C_{30}H_{56}N_5O_7$ | 1.6 |
| **A2_D8_D17** | **18** | C14-*q**tt***L | 672.4542 | 672.4535 | $C_{33}H_{62}N_5O_9$ | 1 |
| **A2_D8_D18** | | C14-*q**t**y*L | 734.4699 | ND | | |

| NRPS- | Compound | Peptide sequence | MS calculated [M+H]+ | MS detected [M+H]+ | Ion Formula | Δppm |
|---|---|---|---|---|---|---|
| **A2_D8_D19** | **19** | C14-*q*T*v*L | 670.475 | 670.4734 | $C_{34}H_{64}N_5O_8$ | 2.3 |
| **A2_D8_D20** | **20** | C14-*q*T*l*L | 684.4906 | 684.4901 | $C_{35}H_{66}N_5O_8$ | 0.7 |
| **A2_D8_D21** | **21** | C14-*q*T*a*L | 642.4437 | 642.4428 | $C_{32}H_{60}N_5O_8$ | 1.4 |
| **A2_D9_D16** | | C14-*qyq*L | 761.4808 | ND | | |
| | **1** | C14-*qq*L | 598.4174 | 598.4161 | $C_{30}H_{56}N_5O_7$ | 1.7 |
| **A2_D9_D17** | | C14-*qy*_**t**_L | 734.4699 | ND | | |
| | **8** | C14-*q*_**t**_L | 571.4066 | 571.4061 | $C_{29}H_{54}N_4O_7$ | 0.7 |
| **A2_D9_D18** | | C14-*qyy*L | 796.4855 | ND | | |
| **A2_D9_D19** | **22** | C14-*q*Y*v*L | 732.4906 | 732.491 | $C_{39}H_{66}N_5O_8$ | 0.5 |
| **A2_D9_D20** | **23** | C14-*q*Y*l*L | 746.5063 | 746.5056 | $C_{40}H_{68}N_5O_8$ | 0.9 |
| **A2_D9_D21** | **24** | C14-*q*Y*a*L | 704.4593 | 704.459 | $C_{37}H_{62}N_5O_8$ | 0.4 |
| | **25** | C12-*q*Y*a*L | 676.428 | 676.4281 | $C_{35}H_{58}N_5O_8$ | 0.2 |
| **A2_D10_D16** | | C14-Q*vq*L | 697.4859 | ND | | |
| | **1** | C14-*qq*L | 598.4174 | 598.4172 | $C_{30}H_{56}N_5O_7$ | 0.2 |
| **A2_D10_D17** | | C14-Q*v*_**t**_L | 670.475 | ND | | |
| | **8** | C14-*q*_**t**_L | 571.4066 | 571.4051 | $C_{29}H_{55}N_4O_7$ | 2.5 |
| **A2_D10_D18** | | C14-Q-*vy*L | 732.4906 | ND | | |
| **A2_D10_D19** | | C14-Q-V*v*L | 668.4957 | ND | | |
| | **11** | C14-*q*L | 470.3589 | 470.3587 | $C_{25}H_{48}N_3O_5$ | 0.3 |
| **A2_D10_D20** | **26** | C14-QV*l*L | 682.5113 | 682.511 | $C_{36}H_{68}N_5O_7$ | 0.4 |
| | **12** | C14-Q*l*L | 583.4429 | 583.4421 | $C_{31}H_{59}N_4O_6$ | 1.4 |
| **A2_D10_D21** | **27** | C14-QVaL | 640.4644 | 640.4635 | $C_{33}H_{62}N_5O_7$ | 1.4 |
| **A2_D11_D16** | **28** | C14-Q*lq*L | 711.5015 | 711.501 | $C_{36}H_{67}N_6O_8$ | 0.6 |
| | **12** | C14-Q*l*L | 583.4429 | 583.4423 | $C_{31}H_{59}N_4O_6$ | 1.1 |

| NRPS- | Compound | Peptide sequence | MS calculated $[M+H]^+$ | MS detected $[M+H]^+$ | Ion Formula | Δppm |
|---|---|---|---|---|---|---|
| **A2_D11_D17** | **29** | C14-Q*lt*L | 684.4906 | 684.4898 | $C_{35}H_{66}N_5O_8$ | 1.1 |
| **A2_D11_D18** | | C14-Q*ly*L | 746.5063 | ND | | |
| **A2_D11_D19** | | C14-QL*v*L | 682.5113 | ND | | |
| | **30** | C14Q*v*L | 569.4273 | 569.4258 | $C_{30}H_{57}N_4O_6$ | 2.6 |
| **A2_D11_D20** | **31** | C14-QL*l*L | 696.527 | 696.5269 | $C_{37}H_{70}N_5O_7$ | 0.1 |
| **A2_D11_D21** | **32** | C14-QL*a*L | 654.48 | 654.4797 | $C_{34}H_{64}N_5O_7$ | 0.4 |
| **A2_D12_D16** | **33** | C14-Q*aq*L | 669.4546 | 669.454 | $C_{33}H_{61}N_6O_8$ | 0.8 |
| | **13** | C14-Q*a*L | 541.396 | 541.3954 | $C_{28}H_{53}N_4O_6$ | 1.1 |
| **A2_D12_D17** | **34** | C14-Q*at*L | 642.4437 | 642.4438 | $C_{32}H_{60}N_5O_8$ | 1.3 |
| **A2_D12_D18** | | C14-Q*ay*L | 704.4593 | ND | | |
| **A2_D12_D19** | **35** | C14-QA*v*L | 640.4644 | 640.464 | $C_{33}H_{62}N_5O_7$ | 0.6 |
| | **30** | C14-Q*v*L | 569.4273 | 569.4266 | $C_{30}H_{57}N_4O_6$ | 0.7 |
| **A2_D12_D20** | **36** | C14-QA*l*L | 654.48 | 654.4806 | $C_{34}H_{64}N_5O_7$ | 0.8 |
| | **37** | C12-QA*l*L | 626.4487 | 626.4485 | $C_{32}H_{60}N_5O_7$ | 0.2 |
| **A2_D12_D21** | **38** | C14-QA*a*L | 612.4331 | 612.4326 | $C_{31}H_{58}N_5O_7$ | 0.9 |
| **A2_D13_D16** | | C14-Q*vq*L | 697.4859 | ND | | |
| | **11** | C14-*q*L | 470.3589 | 470.3587 | $C_{25}H_{48}N_3O_5$ | 0.3 |
| **A2_D13_D17** | | C14-Q*vt*L | 670.475 | ND | | |
| | **11** | C14-*q*L | 470.3589 | 470.3585 | $C_{25}H_{48}N_3O_5$ | 0.7 |
| **A2_D13_D18** | | C14-Q*vy*L | 732.4906 | ND | | |
| | **11** | C14-*q*L | 470.3589 | 470.3587 | $C_{25}H_{48}N_3O_5$ | 0.3 |
| **A2_D13_D19** | | C14-QV*v*L | 668.4957 | ND | | |
| | **30** | C14-Q*v*L | 569.4273 | 569.4267 | $C_{30}H_{57}N_4O_6$ | 1.1 |
| **A2_D13_D20** | **26** | C14-QV*l*L | 682.5113 | 682.5109 | $C_{36}H_{68}N_5O_7$ | 0.6 |

| NRPS- | Compound | Peptide sequence | MS calculated $[M+H]^+$ | MS detected $[M+H]^+$ | Ion Formula | Δppm |
|---|---|---|---|---|---|---|
| | **31** | C14-QL*l*L | 696.527 | 696.5266 | $C_{37}H_{70}N_5O_7$ | 0.6 |
| **A2_D13_D21** | **27** | C14-QV*a*L | 640.4644 | 640.4645 | $C_{33}H_{62}N_5O_7$ | 0.2 |
| | **32** | C14-QL*a*L | 654.48 | 654.4797 | $C_{34}H_{64}N_5O_7$ | 0.6 |
| **A2_D14_D16** | | C14-Q*vq*L | 697.4859 | ND | | |
| | **1** | C14-*qq*L | 598.4174 | 598.417 | $C_{30}H_{56}N_5O_7$ | 0.7 |
| **A2_D14_D17** | | C14-Q*vt*L | 670.475 | ND | | |
| | **8** | C14-*qt*L | 571.4066 | 571.4059 | $C_{29}H_{55}N_4O_7$ | 1.2 |
| **A2_D14_D18** | | C14-Q*vy*L | 732.4906 | ND | | |
| | **11** | C14-*q*L | 470.3589 | 470.3586 | $C_{25}H_{48}N_3O_5$ | 0.6 |
| **A2_D14_D19** | | C14-QV*v*L | 668.4957 | ND | | |
| | **11** | C14-*v*L | 470.3589 | 470.3585 | $C_{25}H_{48}N_3O_5$ | 0.5 |
| **A2_D14_D20** | | C14-QV*l*L | 682.5113 | ND | | |
| | **12** | C14-Q*l*L | 583.4429 | 583.4429 | $C_{31}H_{59}N_4O_6$ | 0.1 |
| **A2_D14_D21** | | C14-Q-V*l*L | 640.4644 | ND | | |
| | **13** | C14-Q*l*L | 541.396 | 541.3956 | $C_{28}H_{53}N_4O_6$ | 0.7 |
| **A2_D7_D22** | **16** | C14-*q*-Q*l*L | 697.4859 | 697.4855 | $C_{35}H_{65}N_6O_8$ | 0.5 |
| | **39** | C14-*q*-Q*l*L | 711.5015 | 711.5011 | $C_{36}H_{67}N_6O_8$ | 0.6 |
| | **30** | C14-Q*v*L | 569.4273 | 569.4266 | $C_{30}H_{57}N_4O_6$ | 1.1 |
| | **12** | C14-Q*l*L | 583.4429 | 583.4418 | $C_{31}H_{59}N_4O_6$ | 1.9 |
| **A2_D8_D22** | **19** | C14-*q*T*v*L | 670.475 | 670.475 | $C_{34}H_{64}N_5O_8$ | 0.1 |
| | **20** | C14-*q*T*l*L | 684.4906 | 684.4908 | $C_{35}H_{66}N_5O_8$ | 0.3 |
| | **40** | C12-*q*T*l*L | 656.4593 | 656.4589 | $C_{33}H_{62}N_5O_8$ | 0.6 |
| | **41** | C12-*q*T*v*L | 642.4437 | 642.4432 | $C_{32}H_{60}N_5O_8$ | 0.6 |

| NRPS- | Compound | Peptide sequence | MS calculated [M+H]$^+$ | MS detected [M+H]$^+$ | Ion Formula | Δppm |
|---|---|---|---|---|---|---|
| **A2_D9_D22** | **22** | C14-*q*Y*v*L | 732.4906 | 732.4906 | $C_{39}H_{66}N_5O_8$ | 0 |
| | **23** | C14-*q*Y*l*L | 746.5063 | 746.506 | $C_{40}H_{68}N_5O_8$ | 0.3 |
| | **12** | C14-Q*l*L | 583.4429 | 583.4423 | $C_{31}H_{59}N_4O_6$ | 1.3 |
| | **10** | C14-*qy*L | 633.4222 | 633.4217 | $C_{34}H_{57}N_4O_7$ | 0.8 |
| **A2_D10_D22** | **42** | C14-QV*v*L | 668.4957 | 668.4957 | $C_{35}H_{66}N_5O_7$ | 0.1 |
| | **30** | C14-Q*v*L | 569.4273 | 569.4268 | $C_{30}H_{57}N_4O_6$ | 0.9 |
| | **12** | C14-Q*l*L | 583.4429 | 583.4428 | $C_{31}H_{59}N_4O_6$ | 0.2 |
| **A2_D11_D22** | **43** | C14-QL*v*L | 682.5113 | 682.5111 | $C_{36}H_{68}N_5O_7$ | 0.4 |
| | **31** | C14-QL*l*L | 696.527 | 696.5266 | $C_{37}H_{70}N_5O_7$ | 0.5 |
| **A2_D12_D22** | **35** | C14-QA*v*L | 640.4644 | 640.4637 | $C_{33}H_{62}N_5O_7$ | 1.1 |
| | **36** | C14-QA*l*L | 654.48 | 654.4788 | $C_{34}H_{64}N_5O_7$ | 1.9 |
| **A2_D7_D23** | **16** | C14-*q*Q*v*L | 697.4859 | 697.4851 | $C_{35}H_{65}N_6O_8$ | 1 |
| | **30** | C14-Q*v*L | 569.4273 | 569.4265 | $C_{30}H_{57}N_4O_6$ | 1.3 |
| **A2_D8_D23** | **19** | C14-*q*T*v*L | 670.475 | 670.4742 | $C_{35}H_{65}N_6O_8$ | 1.2 |
| **A2_D9_D23** | | C14-*q*Y*v*L | 732.4906 | ND | | |
| **A2_D9_D23** | **10** | C14-*qy*L | 633.4222 | 644.4216 | $C_{34}H_{57}N_4O_7$ | 0.9 |
| **A2_D10_D23** | **42** | C14-QV*v*L | 668.4957 | 668.4954 | $C_{35}H_{66}N_5O_7$ | 0.5 |
| **A2_D11_D23** | | C14-QL*v*L | 682.5113 | ND | | |
| | **10** | C14-*qy*L | 633.4222 | 633.4219 | $C_{34}H_{57}N_4O_7$ | 0.5 |
| **A2_D12_D23** | **35** | C14-QA*v*L | 640.4644 | 640.4646 | $C_{33}H_{62}N_5O_7$ | 0.4 |
| | **30** | C14-Q*v*L | 569.4273 | 569.4268 | $C_{30}H_{57}N_4O_6$ | 0.8 |
| **A2_D13_D22** | | C14-QV*v*L | 668.4957 | ND | | |
| | **11** | C14-*q*L | 470.3589 | 470.3579 | $C_{25}H_{48}N_3O_5$ | 2 |

| NRPS- | Compound | Peptide sequence | MS calculated [M+H]$^+$ | MS detected [M+H]$^+$ | Ion Formula | Δppm |
|---|---|---|---|---|---|---|
| **A2_D13_D23** | **42** | C14-QV*v*L | 668.4957 | 668.4956 | $C_{35}H_{66}N_5O_7$ | 0.1 |
| | **43** | C14-QL*v*L | 682.5113 | 682.5106 | $C_{36}H_{68}N_5O_7$ | 1 |
| **A2_D14_D22** | | C14-QV*v*L | 668.4957 | ND | | |
| | **30** | C14-Q*v*L | 569.4273 | 569.4272 | $C_{30}H_{57}N_4O_6$ | 0.1 |
| **A2_D14_D23** | | C14-QV*v*L | 668.4957 | ND | | |
| | **30** | C14-Q*v*L | 569.4273 | 569.4273 | $C_{30}H_{57}N_4O_6$ | 0 |
| **A2_D15_D16** | | C14-*qyq*L | 761.4808 | ND | NA | ND |
| | **11** | C14-*q*L | 470.3589 | 470.3584 | $C_{25}H_{48}N_3O_5$ | 1.1 |
| **A2_D15_D17** | | C14-*qyt*L | 734.4699 | ND | | |
| | **11** | C14-*q*L | 470.3589 | 470.3586 | $C_{25}H_{48}N_3O_5$ | 0.6 |
| **A2_D15_D18** | | C14-*qyy*L | 796.4855 | ND | | |
| **A2_D15_D19** | | C14-*q*Y*v*L | 732.4906 | ND | | |
| | **11** | C14-*q*L | 470.3589 | 470.3585 | $C_{25}H_{48}N_3O_5$ | 0.8 |
| **A2_D15_D20** | **23** | C14-*q*Y*l*L | 746.5063 | 746.5068 | $C_{40}H_{68}N_5O_8$ | 0.8 |
| | **12** | C14-Q*l*L | 583.4429 | 583.4428 | $C_{31}H_{59}N_4O_6$ | 0.3 |
| **A2_D15_D21** | **24** | C14-qYaL | 704.4593 | 704.4587 | $C_{37}H_{62}N_5O_8$ | 0.9 |
| | **13** | C14-Q*a*L | 541.396 | 541.3956 | $C_{28}H_{53}N_4O_6$ | 0.7 |
| **A2_D7_D24** | | C14-*qqy*L | 761.4808 | ND | | |
| | **47** | C14-Q*y*L | 633.4222 | 633.4217 | $C_{34}H_{57}N_4O_7$ | 0.7 |
| **A2_D8_D24** | **44** | C14-*q*T*y*L | 734.4699 | 734.4696 | $C_{38}H_{64}N_5O_9$ | 0.4 |
| | **10** | C14-*qy*L | 633.4222 | 633.4212 | $C_{34}H_{57}N_4O_7$ | 1.5 |
| **A2_D9_D24** | **45** | C14-*q*Y*y*L | 796.4855 | 796.4862 | $C_{43}H_{66}N_5O_9$ | 0.8 |
| | **47** | C14-Q*y*L | 633.4222 | 633.4221 | $C_{34}H_{57}N_4O_7$ | 0.2 |
| **A2_D10_D24** | | C14-Q*vy*L | 732.4906 | ND | | |

| NRPS- | Compound | Peptide sequence | MS calculated [M+H]$^+$ | MS detected [M+H]$^+$ | Ion Formula | Δppm |
|---|---|---|---|---|---|---|
| | **11** | C14-*q*L | 470.3589 | 470.3583 | $C_{25}H_{48}N_3O_5$ | 1.3 |
| **A2_D11_D24** | | C14-Q*ly*L | 746.5063 | ND | | |
| | **47** | C14-Q*y*L | 633.4222 | 633.4213 | $C_{34}H_{57}N_4O_7$ | 1.4 |
| **A2_D12_D24** | | C14-Q*ay*L | 704.4593 | ND | | |
| | **47** | C14-Q*y*L | 633.4222 | 633.4213 | $C_{34}H_{57}N_4O_7$ | 1.4 |
| **A2_D15_D24** | | C14-*qyy*L | 796.4855 | ND | | |
| | **47** | C14-Q*y*L | 633.4222 | 633.4212 | $C_{34}H_{57}N_4O_7$ | 1.6 |
| | **22** | C14-*q*Y*v*L | 732.4906 | ND | | |
| **A2_D15_D22** | **30** | C14-Q*v*L | 569.4273 | 569.4261 | $C_{30}H_{57}N_4O_6$ | 2 |
| | **12** | C14-Q*l*L | 583.4429 | 583.4423 | $C_{31}H_{59}N_4O_6$ | 1.1 |
| **A2_D15_D23** | **22** | C14-*q*Y*v*L | 732.4906 | 732.4907 | $C_{39}H_{66}N_5O_8$ | 0.1 |
| | **30** | C14-Q*v*L | 569.4273 | 569.4267 | $C_{30}H_{57}N_4O_6$ | 0.9 |
| | **46** | C14-QVL*v*L | 781.5798 | 781.5797 | $C_{41}H_{77}N_6O_8$ | 0 |
| **A2_D13_D24** | | C14-Q*vy*L | 732.4906 | ND | | |
| | **47** | C14-Q*y*L | 633.4222 | 633.422 | $C_{34}H_{57}N_4O_7$ | 0.2 |
| **A2_D14_D24** | | C14-Q*vy*L | 732.4906 | ND | | |
| | **47** | C14-Q*y*L | 633.4222 | 633.4225 | $C_{34}H_{57}N_4O_7$ | 0.4 |
| **A3_D25** | **13** | C14-Q*a*L | 541.396 | 541.3959 | $C_{28}H_{53}N_4O_6$ | 0.2 |
| **A3_D26** | **48** | C4-P*a*L | 370.2337 | 370.233 | $C_{18}H_{32}N_3O_5$ | 1.6 |
| | **49** | P*a*L | 300.1918 | 300.1916 | $C_{14}H_{26}N_3O_4$ | 0.7 |
| **A3_D27** | **50** | S*a*L | 290.1711 | 290.1712 | $C_{12}H_{24}N_3O_5$ | 0.4 |
| **A3_D28** | **51** | L*a*L | 316.2231 | 316.2232 | $C_{15}H_{30}N_3O_4$ | 0.5 |

**Table S8. Comparison of the relative production of the compounds 2 – 7 from the NRPS-5, NRPS-5_mod and NRPS-A1_D3.** The three constructs produce the same NRPS, while NRPS-5 was constructed with Gibson assembly, NRPS-5_mod is NRPS-5 after mutagenesis to contain the DNA changes applied with GGA and NRPS-A1_D3 was constructed using GGA with acceptor and donor plasmids. The titers of compounds **2** – **7** from NRPS-5 (by Gibson assembly) are set to 100 %, to which the titers of the compounds from NRPS-5_mod and NRPS-A1_D3 are compared, respectively.

| Compound | Peptide sequence | MS calculated $[M+H]^+$ | NRPS-5 | NRPS-5_mod | NRPS-A1_D3 |
|---|---|---|---|---|---|
| **2** | C14-*q*YW | 706.4174 | 100% | 105% | 97% |
| **3** | C12-*q*YW | 678.3861 | 100% | 115% | 99% |
| **4** | C14-*q*YF | 667.4066 | 100% | 103% | 80% |
| **5** | C12-*q*YF | 639.3752 | 100% | 95% | 82% |
| **6** | C14-*q*FW | 690.4225 | 100% | 121% | 91% |
| **7** | C14-*q*FF | 651.4116 | 100% | 102% | 82% |

**Table S9. List of all compounds produced by and related to the xenoamicin library.** Expected compounds that were not detected are marked as ND (not detected). The N-terminal linear part C4-P*a*V*I*I is abbreviated as “X”. The ring is always formed between Thr-8 and the C-terminal Val-13. Linear products are marked with an “*”. The relative production is calculated using the EIC’s peak area divided by the area of xenoamicin C (**52**) (NRPS-6; wild type in *E. coli*). Small italic letters indicate D-amino acids for the peptide sequence, and small italic bold letters indicate D-*allo*-amino acids.

| XabAB+NRPS- | Compound | Peptide sequence | MS calculated $[M+2H]^{2+}$ | MS detected $[M+2H]^{2+}$ | Ion Formula | Δppm | Rel. Prod. |
|---|---|---|---|---|---|---|---|
| **6** | **52** | X***t***V***i***VaβAPV | 657.9235 | 657.9229 | $C_{65}H_{113}N_{13}O_{15}$ | 0.7 | 100% |
| | **53** | X***t***V*I*V*a*βAPV | 657.9235 | 657.9231 | $C_{65}H_{113}N_{13}O_{15}$ | 0.5 | 7.0% |
| | **54** | X***t***VvV*a*βAPV | 650.9156 | 650.9154 | $C_{64}H_{111}N_{13}O_{15}$ | 0.3 | 6.3% |
| **7** | **52** | X***t***V***i***V*a*βAPV | 657.9235 | 657.9233 | $C_{65}H_{113}N_{13}O_{15}$ | 0.2 | 111% |
| | **53** | X***t***V*I*V*a*βAPV | 657.9235 | 657.9235 | $C_{65}H_{113}N_{13}O_{15}$ | 0.1 | 13% |
| | **54** | X***t***V*v*V*a*βAPV | 650.9156 | 650.9157 | $C_{64}H_{111}N_{13}O_{15}$ | 0.1 | 6.2% |
| **A4_D29** | **55** | X***t***V***i***V*a*βAV | 609.3971 | 609.397 | $C_{60}H_{106}N_{12}O_{14}$ | 0.1 | 6.9% |
| **A4_D30** | **56** | X***t***V***i***VAPV | 622.4049 | 622.4049 | $C_{62}H_{108}N_{12}O_{14}$ | 0.1 | 19% |
| | **57** | X***t***V***i***VAPV* | 631.4102 | 631.4098 | $C_{62}H_{110}N_{12}O_{15}$ | 0.5 | 10% |
| **A4_D31_D33** | | X***t***V***i***VAAAV | 644.9156 | ND | | | |
| **A4_D31_D34** | **52** | X***t***V***i***VaβAPV | 657.9235 | 657.9236 | $C_{65}H_{113}N_{13}O_{15}$ | 0.2 | 54% |
| | **53** | X***t***V*I*V*a*βAPV | 657.9235 | 657.9241 | $C_{65}H_{113}N_{13}O_{15}$ | 2 | 6.1% |
| | **54** | X***t***VvV*a*βAPV | 650.9156 | 650.9158 | $C_{64}H_{111}N_{13}O_{15}$ | 0.3 | 3.7% |
| **A4_D32_D33** | | X***t***V***i***VA*p*βAV | 657.9235 | ND | | | |
| **A4_D32_D34** | | X***t***V***i***VAPPV | 670.9313 | ND | | | |
| **A4_D1** | **58** | X***t***V***i***V*a*QV | 637.9078 | 637.907 | $C_{62}H_{109}N_{13}O_{15}$ | 0.7 | 0.35% |
| **A4_D2** | **59** | X***t***V***i***V*a*TV | 624.4024 | 624.4021 | $C_{61}H_{108}N_{12}O_{15}$ | 0.3 | 6.4% |

| XabAB+NRPS- | Compound | Peptide sequence | MS calculated $[M+2H]^{2+}$ | MS detected $[M+2H]^{2+}$ | Ion Formula | Δppm | Rel. Prod. |
|---|---|---|---|---|---|---|---|
| | **60** | XtViVaTV* | 633.4077 | 633.4082 | $C_{61}H_{110}N_{12}O_{16}$ | 1 | 1.8% |
| **A4_D3** | **61** | XtViVaYV | 655.4102 | 655.4104 | $C_{66}H_{110}N_{12}O_{15}$ | 0.4 | 15% |
| **A4_D4** | **62** | XtViVAVV | 623.4127 | 623.4126 | $C_{62}H_{110}N_{12}O_{14}$ | 0.2 | 4.1% |
| **A4_D5** | **63** | XtViVALV | 630.4206 | 630.4208 | $C_{63}H_{112}N_{12}O_{14}$ | 0.5 | 17% |
| | **64** | XtViVALV* | 639.4258 | 639.4255 | $C_{63}H_{114}N_{12}O_{15}$ | 0.5 | 0.69% |
| **A4_D6** | **65** | XtViVAAV | 609.3971 | 609.3973 | $C_{60}H_{106}N_{12}O_{14}$ | 0.5 | 49% |
| **A4_D7_D16** | | XtViVaqQV | 701.9371 | ND | | | |
| **A4_D7_D17** | | XtViVaqTV | 688.4317 | ND | | | |
| **A4_D7_D18** | | XtViVaqYV | 719.4395 | ND | | | |
| **A4_D7_D19** | | XtViVaQVV | 687.4420 | ND | | | |
| **A4_D7_D20** | | XtViVaQLV | 694.4498 | ND | | | |
| **A4_D7_D21** | | XtViVaQAV | 673.4264 | ND | | | |
| **A4_D8_D16** | | XtViVatQV | 688.4317 | ND | | | |
| **A4_D8_D17** | | XtViVatTV | 674.9262 | ND | | | |
| | **59** | XtViVaTV | 624.4024 | 624.402 | $C_{61}H_{108}N_{12}O_{15}$ | 0.5 | 2.0% |
| **A4_D8_D18** | | XtViVatYV | 705.9340 | ND | | | |
| | **59** | XtViVaTV | 624.4024 | 624.402 | $C_{61}H_{108}N_{12}O_{15}$ | 0.5 | 1.3% |
| **A4_D8_D19** | **66** | XtViVaTVV | 673.9366 | 673.9364 | $C_{66}H_{117}N_{13}O_{16}$ | 0.2 | 3.2% |
| **A4_D8_D20** | **67** | XtViVaTLV | 680.9444 | ND | | | |
| | **59** | XtViVATV | 624.4024 | 624.4014 | $C_{61}H_{108}N_{12}O_{15}$ | 1.5 | 0.60% |
| **A4_D8_D21** | **68** | XtViVaTAV | 659.9209 | 659.9216 | $C_{64}H_{113}N_{13}O_{16}$ | 1.1 | 1.1% |
| | **59** | XtViVaTV | 624.4024 | 624.403 | $C_{61}H_{108}N_{12}O_{15}$ | 1.2 | 0.62% |
| **A4_D9_D16** | | XtViVayQV | 719.4395 | ND | | | |
| | **58** | XtViVaQV | 637.9078 | 637.9076 | $C_{62}H_{109}N_{13}O_{15}$ | 0.2 | 5.0% |

| XabAB+NRPS- | Compound | Peptide sequence | MS calculated $[M+2H]^{2+}$ | MS detected $[M+2H]^{2+}$ | Ion Formula | Δppm | Rel. Prod. |
|---|---|---|---|---|---|---|---|
| A4_D9_D17 | | X***t***V***i***V*ay*TV | 705.9340 | ND | | | |
| | 60 | X***t***V***i***V*a*TV | 624.4024 | 624.4024 | $C_{61}H_{108}N_{12}O_{15}$ | 0.2 | 6.9% |
| | 69 | X***t***V***i***VAV | 573.8785 | 573.8785 | $C_{57}H_{101}N_{11}O_{13}$ | 0 | 3.7% |
| A4_D9_D18 | | X***t***V***i***V*ay*YV | 736.9419 | ND | | | |
| | 61 | X***t***V***i***V*a*YV | 655.4102 | 655.4102 | $C_{66}H_{110}N_{12}O_{15}$ | 0.1 | 1.1% |
| A4_D9_D19 | 70 | X***t***V***i***VaYVV | 704.9444 | 704.9444 | $C_{71}H_{119}N_{13}O_{16}$ | 0.2 | 1.6% |
| | 62 | X***t***V***i***VAVV | 623.4127 | 623.412 | $C_{62}H_{110}N_{12}O_{14}$ | 1.1 | 2.9% |
| A4_D9_D20 | | X***t***V***i***VaYLV | 711.9522 | ND | | | |
| | 63 | X***t***V***i***VALV | 630.4206 | 630.4209 | $C_{63}H_{112}N_{12}O_{14}$ | 0.6 | 29% |
| A4_D9_D21 | 71 | X***t***V***i***VaYAV | 690.9287 | 690.9288 | $C_{69}H_{115}N_{13}O_{16}$ | 0.2 | 4.6% |
| | 65 | X***t***V***i***VAAV | 609.3971 | 609.3971 | $C_{60}H_{106}N_{12}O_{14}$ | 0.1 | 1.4% |
| | 69 | X***t***V***i***VAV | 573.8785 | 573.8787 | $C_{57}H_{101}N_{11}O_{13}$ | 0.4 | 1.3% |
| A4_D10_D16 | | X***t***V***i***VA*v*QV | 687.4420 | ND | | | |
| A4_D10_D17 | | X***t***V***i***VA*v*TV | 673.9366 | ND | | | |
| A4_D10_D18 | | X***t***V***i***VA*v*YV | 704.9444 | ND | | | |
| A4_D10_D19 | 72 | X***t***V***i***VAVVV | 672.9469 | 672.946 | $C_{67}H_{119}N_{13}O_{15}$ | 1.4 | 0.2% |
| A4_D10_D20 | 73 | X***t***V***i***VAVLV | 679.9548 | 679.9545 | $C_{68}H_{121}N_{13}O_{15}$ | 0.2 | 0.4% |
| A4_D10_D21 | 74 | X***t***V***i***VAVAV | 658.9313 | 658.9313 | $C_{65}H_{115}N_{13}O_{15}$ | 0.2 | 1.5% |
| A4_D11_D16 | | X***t***V***i***VA*l*QV | 694.4498 | ND | | | |
| | 69 | X***t***V***i***VAV | 573.8785 | 573.8787 | $C_{57}H_{101}N_{11}O_{13}$ | 0.5 | 3.9% |
| A4_D11_D17 | | X***t***V***i***VAlTV | 680.9444 | ND | | | |
| | 75 | X***t***V***i***VAaTV | 659.9209 | 659.921 | $C_{64}H_{113}N_{13}O_{16}$ | 0.2 | 50% |
| | 69 | X***t***V***i***VAV | 573.8785 | 573.878 | $C_{57}H_{101}N_{11}O_{13}$ | 0.7 | 4.4% |

| XabAB+NRPS- | Compound | Peptide sequence | MS calculated $[M+2H]^{2+}$ | MS detected $[M+2H]^{2+}$ | Ion Formula | Δppm | Rel. Prod. |
|---|---|---|---|---|---|---|---|
| **A4_D11_D18** | | X***t***V***i***VAIYV | 711.9522 | ND | | | |
| | **69** | X***t***V***i***VAV | 573.8785 | 573.8785 | $C_{57}H_{101}N_{11}O_{13}$ | 0.1 | 6.3% |
| **A4_D11_D19** | | X***t***V***i***VALVV | 679.9548 | ND | | | |
| | **69** | X***t***V***i***VAV | 573.8785 | 573.8785 | $C_{57}H_{101}N_{11}O_{13}$ | 0.1 | 3.0% |
| **A4_D11_D20** | | X***t***V***i***VALLV | 686.9626 | ND | | | |
| | **76** | X***t***-V***i***VALAV | 665.9391 | 665.9391 | $C_{66}H_{117}N_{13}O_{15}$ | 0.1 | 1.3% |
| | **69** | X***t***V***i***VAV | 573.8785 | 573.8786 | $C_{57}H_{101}N_{11}O_{13}$ | 0.2 | 4.1% |
| **A4_D11_D21** | | X***t***V***i***VALAV | 665.9391 | ND | | | |
| | **77** | X***t***V***i***VAAAV | 644.9156 | 644.9157 | $C_{63}H_{111}N_{13}O_{15}$ | 0.2 | 8.6% |
| | **69** | X***t***V***i***VAV | 573.8785 | 573.8787 | $C_{57}H_{101}N_{11}O_{13}$ | 0.4 | 3.7% |
| **A4_D12_D16** | | X***t***V***i***VA*a*QV | 673.4264 | ND | | | |
| | **78** | X***t***V***i***VA*l*QV | 694.4498 | 694.4502 | $C_{68}H_{120}N_{14}O_{16}$ | 0.6 | 0.68% |
| **A4_D12_D17** | | X***t***V***i***VA*a*TV | 659.9209 | ND | | | |
| **A4_D12_D18** | | X***t***V***i***VA*a*YV | 690.9287 | ND | | | |
| **A4_D12_D19** | | X***t***V***i***VAAVV | 658.9313 | ND | | | |
| **A4_D12_D20** | | X***t***V***i***VAALV | 665.9391 | ND | | | |
| | **63** | X***t***V***i***VALV | 630.4206 | 630.4213 | $C_{63}H_{112}N_{12}O_{14}$ | 1.3 | 1.3% |
| **A4_D12_D21** | | X***t***V***i***VAAAV | 644.9156 | ND | | | |
| **A4_D12_D21** | **79** | X***t***V***i***VAALV | 665.9391 | 665.9388 | $C_{66}H_{117}N_{13}O_{15}$ | 0.3 | 1.1% |

### D. Supplementary Figures

**A**

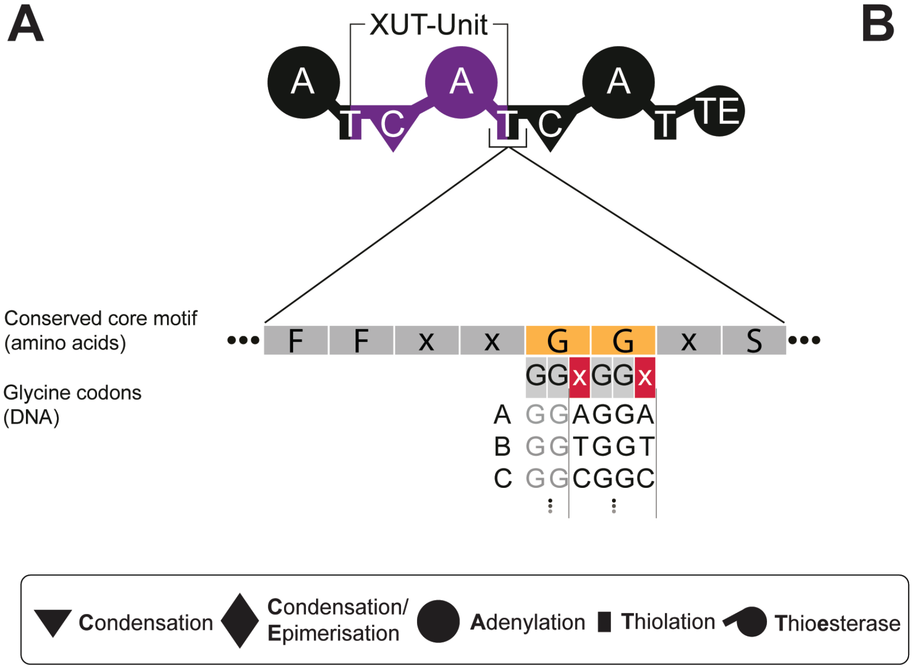


**B**

| Name | Overhang |
|---|---|
| A | AGGA |
| B | TGGT |
| C | CGGC |
| D | GGGG |
| E | AGGT |
| F | TGGC |
| G | CGGG |
| H | AGGC |
| I | TGGG |
| J | GGGA |
| K | AGGG |
| L | CGGA |
| M | GGGT |
| N | TGGA |
| O | CGGT |
| P | GGGC |
| S | CATG |
| T | TGAT |

**Figure S1. Schematic representation of a basic NRPS showing its architecture as well as the conserved motif for the XUT$^{IV}$ fusions side with its possible GGA overhangs. A)** Basic NRPS architecture and the conserved motif for the XUT$^{IV}$ fusion site. Both glycines are marked in orange, providing the codons for the overhangs. The two variable positions for the overhangs are marked in red. **B)** Table for all sixteen possible overhangs, using the flexibility of the third base from the glycine codon. The starter (S) and termination (T) overhangs for the starter and termination modules are also attached.

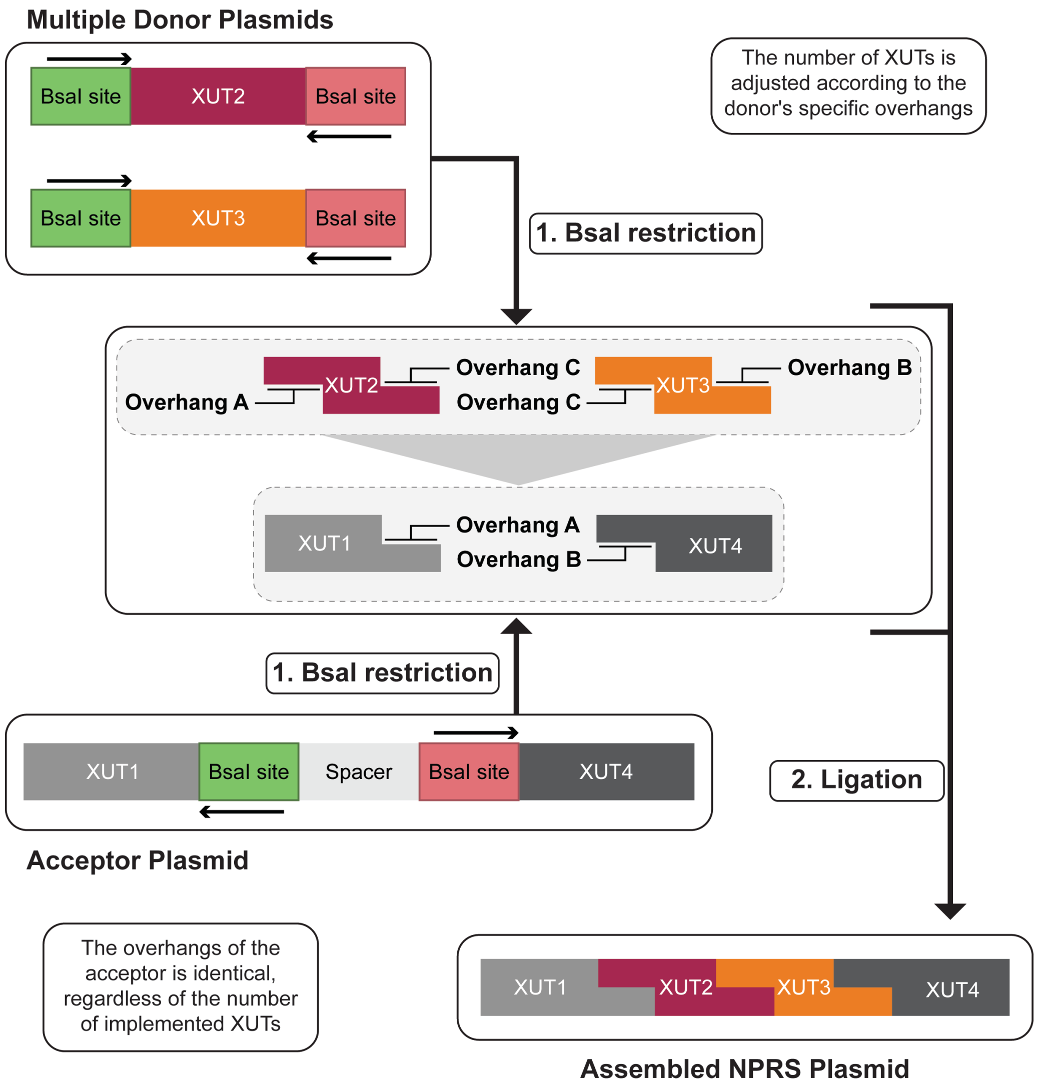


**Figure S2. Schematic GGA acceptor and donor system design for XUT-based NRPS engineering on the DNA level.** The example refers to implementing two XUT modules. The Donor plasmid sequence design with XUT2 & XUT3 on two plasmids contains two flanking BsaI restriction sites (red and green). The acceptor plasmid sequence design contains XUT1 (light grey) and XUT4 (dark grey), with a spacer sequence between both modules. The BsaI sites are highlighted in green (upstream) and red (downstream). The linearised acceptor with overhangs A and B indicated after BsaI restriction, and the cleaved XUT2 & XUT3 DNA fragment with the indicated overhangs A/C and C/B are assembled between XUT1 and XUT4, creating a four-modular NRPS.

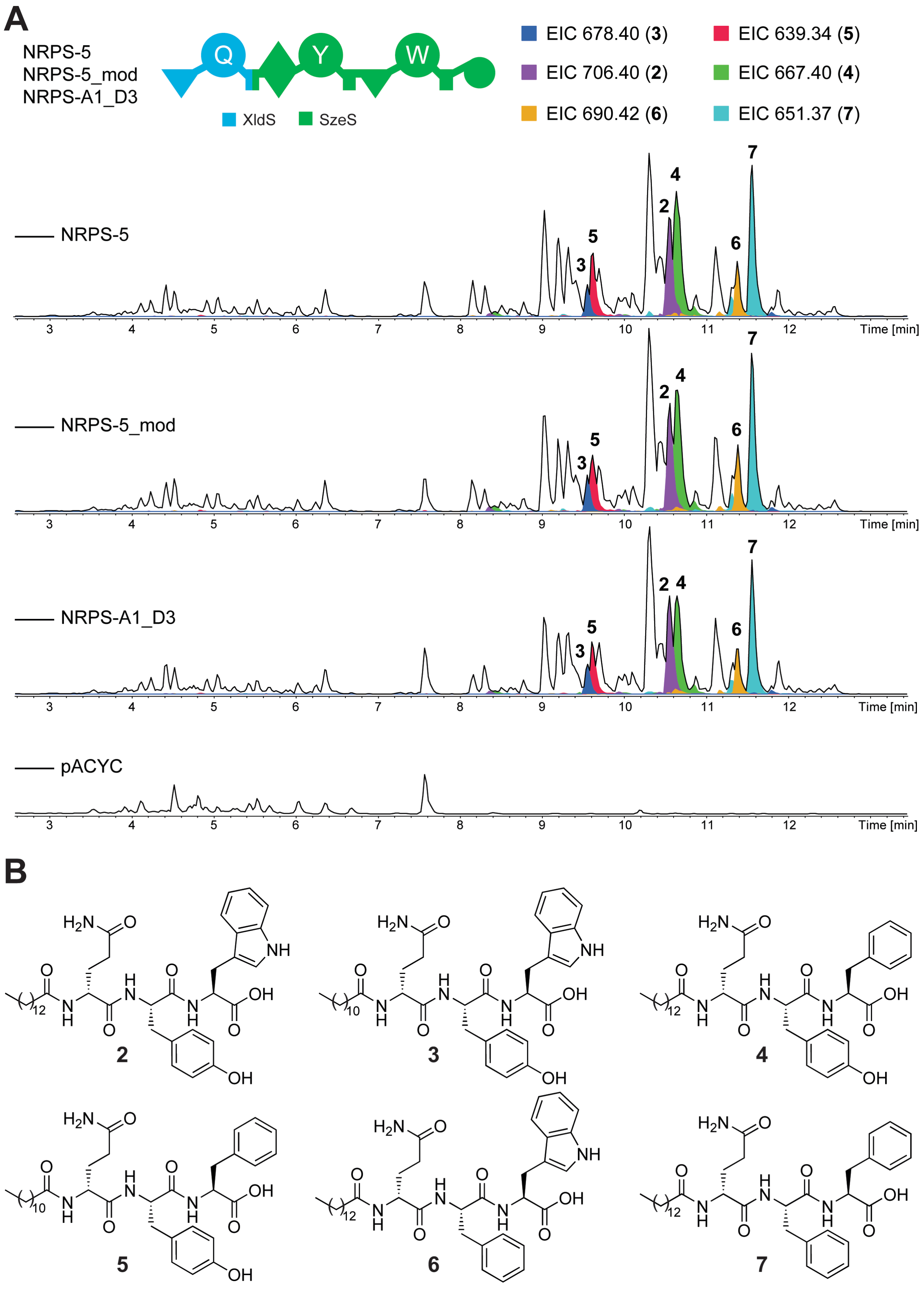
A
NRPS-5
NRPS-5_mod
NRPS-A1_D3
Q
Y
W
XldS
SzeS
EIC 678.40 (3)
EIC 639.34 (5)
EIC 706.40 (2)
EIC 667.40 (4)
EIC 690.42 (6)
EIC 651.37 (7)
NRPS-5
NRPS-5_mod
NRPS-A1_D3
pACYC
Time [min]
B
2
3
4
5
6
7

**Figure S3. HPLC/MS analysis of the produced NRPS-5, NRPS-5 modified and NRPS-A1_D3 in *E. coli* DH10B*::mtaA*. A**) Schematic representation of NRPS-5, NRPS-5 modified and NRPS-A1_D3 with the colour code of the original NRPS. The A domain specificities are indicated with Q = glutamine, Y = tyrosine and W = tryptophan. All three NRPS produce the same compounds. NRPS-5 was assembled using Gibson assembly, while NRPS-A1_D3 was assembled with GGA. NRPS-5 modified is a mutated version of NRPS-5, where the glycine codons for both fusion sides were changed to represent the ones after GGA. HPLC/MS data of compounds **2** – **7** produced in *E. coli* DH10B::*mtaA* expressing NRPS-5, NRPS-5 modified and NRPS-A1_D3. The Base Peak Chromatogram (BPC, black line) and the Extracted Ion Chromatogram (EIC, below with colours according to the depicted legend) of **2** (*m/z* $[M+H]^+$ = 706.40), **3** (*m/z* $[M+H]^+$ = 678.40), **4** (*m/z* $[M+H]^+$ = 667.40), **5** (*m/z* $[M+H]^+$ = 639.34), **6** (*m/z* $[M+H]^+$ = 690.42), and **7** (*m/z* $[M+H]^+$ = 651.37). The pACYC expression was used as a negative control (dotted line). **B**) Chemical structure of the peptides **2** – **7**.

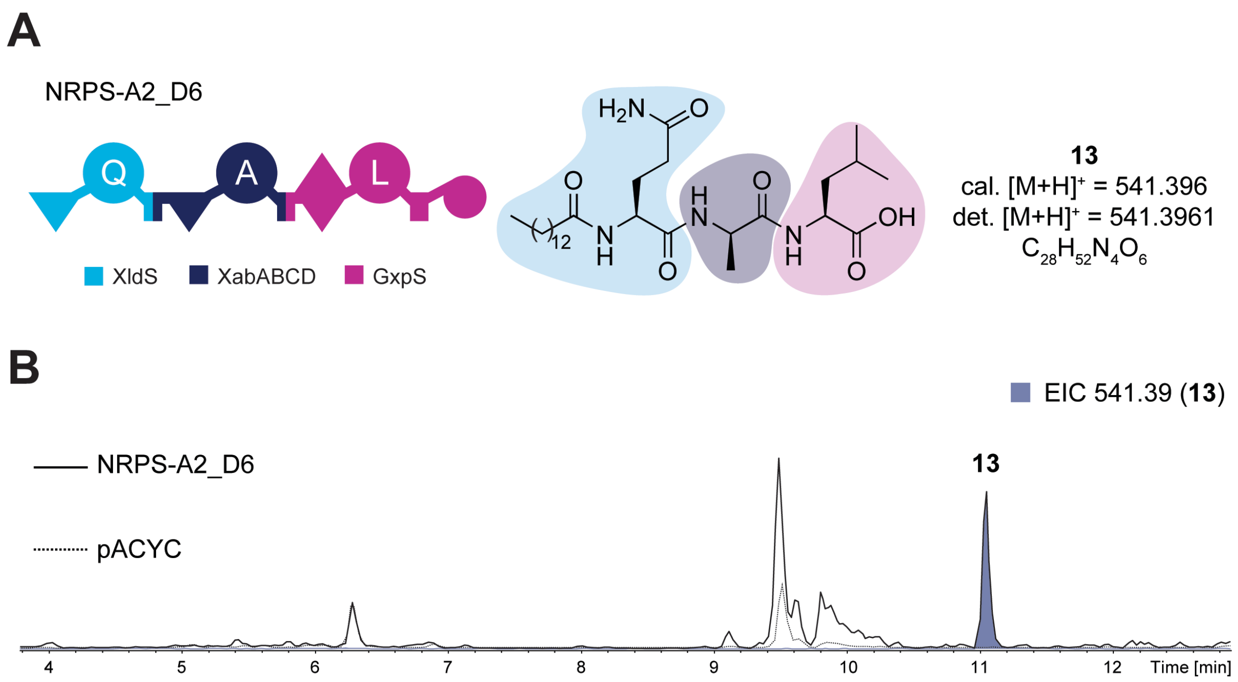


**Figure S4. HPLC/MS analysis of the produced NRPS-A2_D6 in *E. coli* DH10B::*mtaA*.** **A**) Schematic representation of NRPS-A2_D6 with the colour code of the original NRPS. The A domain specificities are indicated with Q = glutamine, A = alanine and L = leucine. Chemical structure of the peptide **13** with its sum formula, the calculated mass and the measured high-resolution mass $[M+H]^+$. **B**) HPLC/MS data of compound **13** produced in *E. coli* DH10B::*mtaA* expressing NRPS-A2_D6. The Base Peak Chromatogram (BPC, black line) and the Extracted Ion Chromatogram (EIC, below with colours according to the depicted legend) of **13** (*m/z* $[M+H]^+$ = 541.39). The pACYC expression was used as a negative control (dotted line).

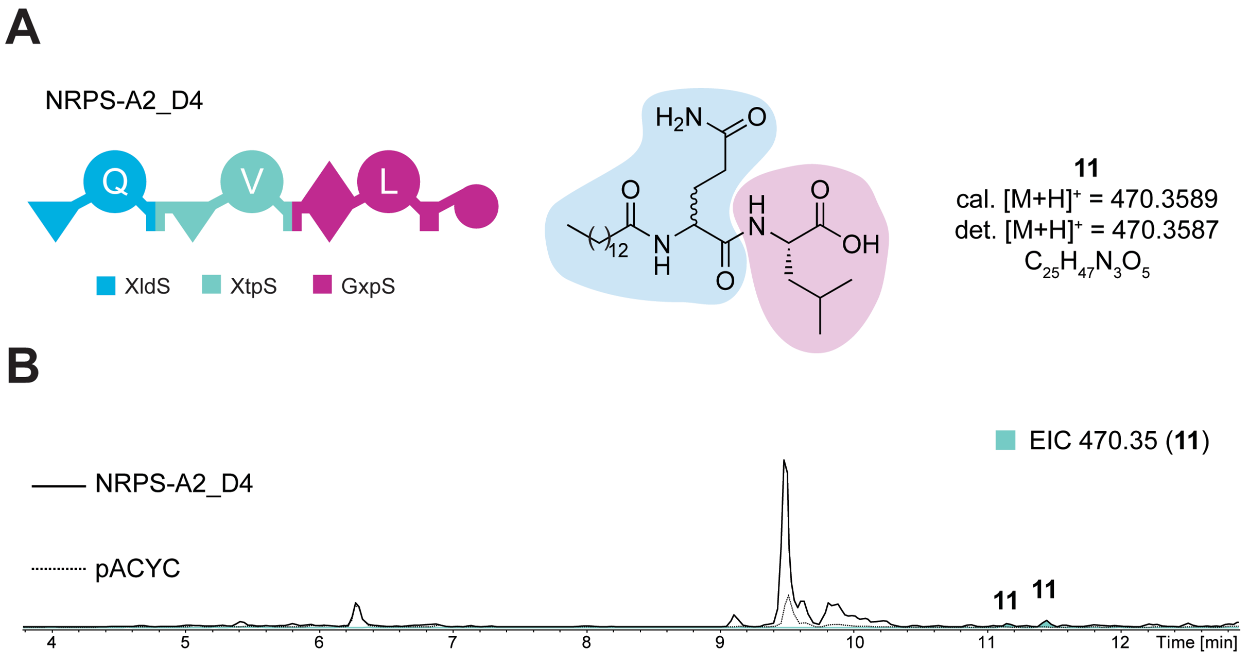


**Figure S5. HPLC/MS analysis of the produced NRPS-A2_D4 in *E. coli* DH10B*::mtaA*.** **A**) Schematic representation of NRPS-A2_D4 with the colour code of the original NRPS. The A domain specificities are indicated with Q = glutamine, V = valine and L = leucine. Chemical structure of the peptide **11** with its sum formula, the calculated mass and the measured high-resolution mass $[M+H]^+$. **B**) HPLC/MS data of compound **11** produced in *E. coli* DH10B::*mtaA* expressing NRPS-A2_D4. The Base Peak Chromatogram (BPC, black line) and the Extracted Ion Chromatogram (EIC, below with colours according to the depicted legend) of **11** (*m/z* $[M+H]^+$ = 470.35). The pACYC expression was used as a negative control (dotted line).

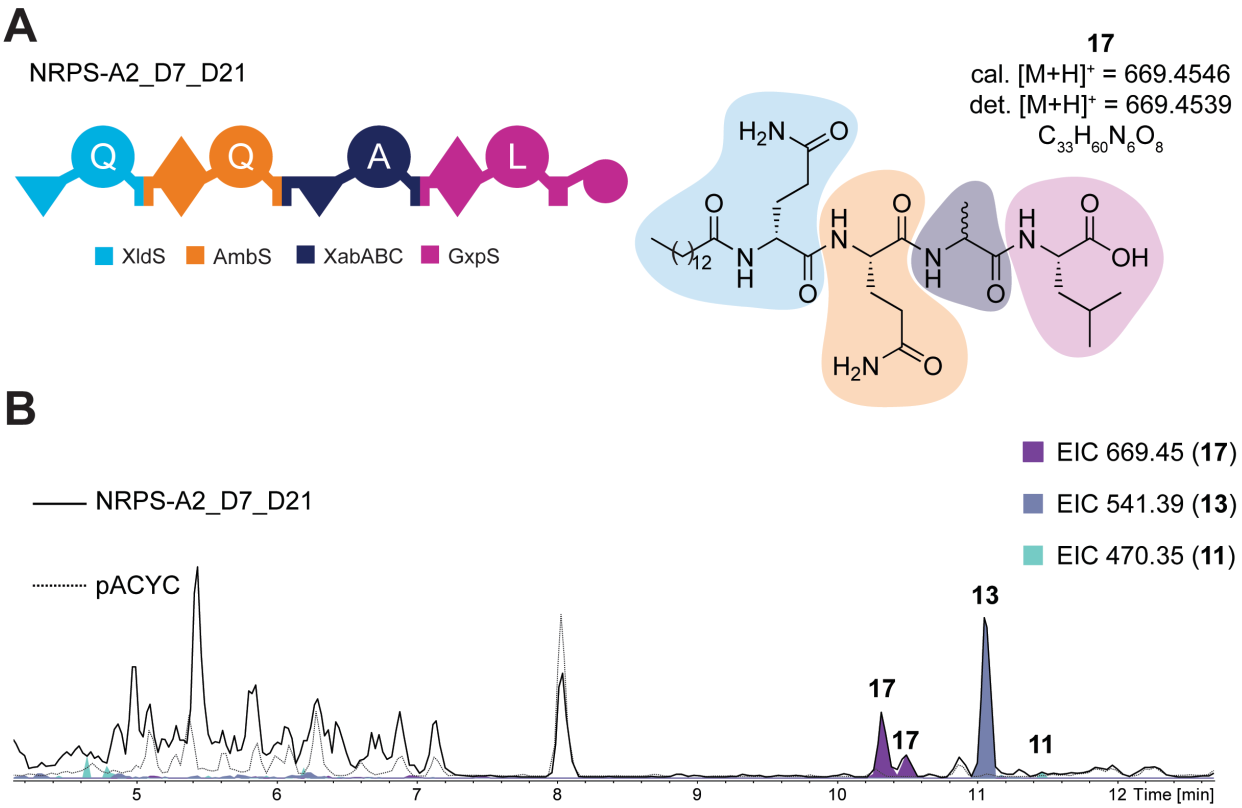


**Figure S6. HPLC/MS analysis of the produced NRPS-A2_D7_D21 in *E. coli* DH10B*::mtaA*. A**) Schematic representation of NRPS-A2_D7_D21 with the colour code of the original NRPS. The A domain specificities are indicated with Q = glutamine, A = alanine and L = leucine. Chemical structure of the peptide **17** with its sum formula, the calculated mass and the measured high-resolution mass $[M+H]^+$. **B**) HPLC/MS data of compounds **17**, **13** (see Fig. S4) and **11** (see Fig. S5) produced in *E. coli* DH10B::*mtaA* expressing NRPS-A2_D7_D21. The Base Peak Chromatogram (BPC, black line) and the Extracted Ion Chromatogram (EIC, below with colours according to the depicted legend) of **17** (*m/z* $[M+H]^+$ = 669.45), **13** (*m/z* $[M+H]^+$ = 541.39) and **11** (*m/z* $[M+H]^+$ = 470.35). The pACYC expression was used as a negative control (dotted line).

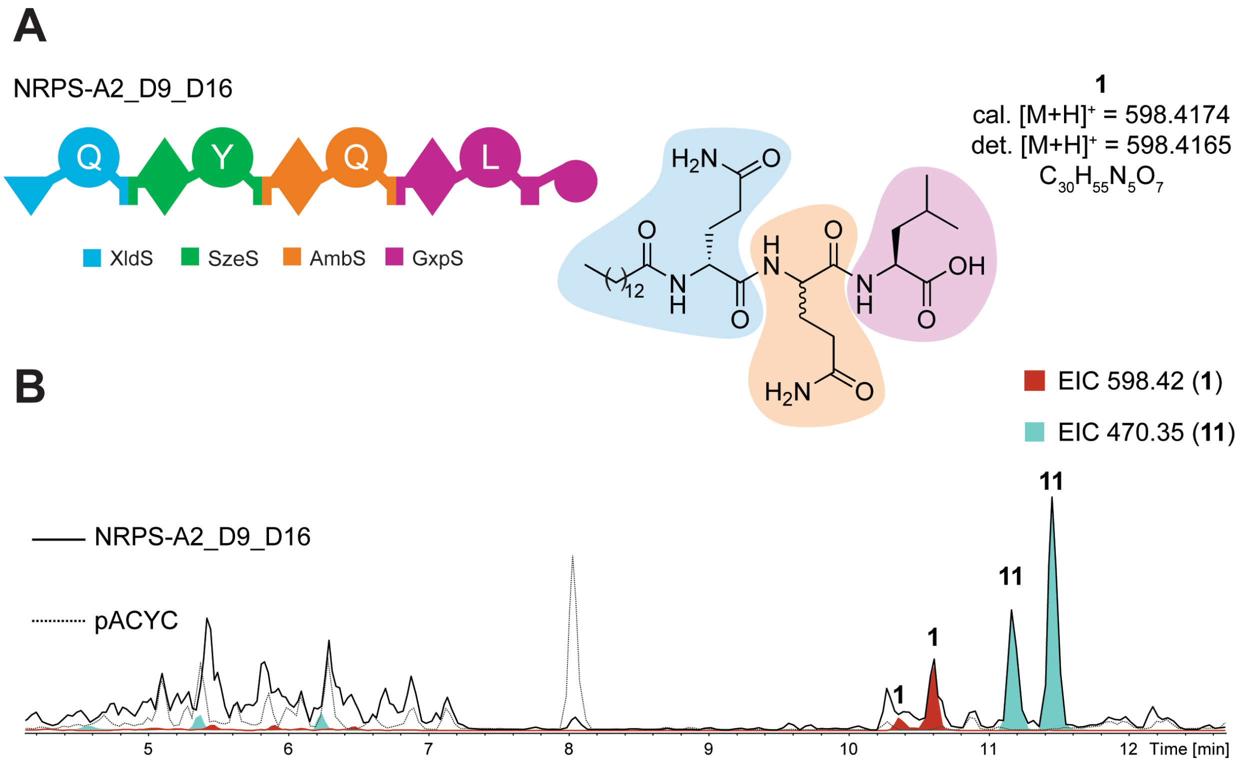


**Figure S7. HPLC/MS analysis of the produced NRPS-A2_D9_D16 in *E. coli* DH10B*::mtaA*. A**) Schematic representation of NRPS-A2_D9_D16 with the colour code of the original NRPS. The A domain specificities are indicated with Q = glutamine, A = alanine and L = leucine. Chemical structure of the peptide **1** with its sum formula, the calculated mass and the measured high-resolution mass $[M+H]^+$. **B**) HPLC/MS data of compounds **1** and **11** (see Fig. S5) produced in *E. coli* DH10B::*mtaA* expressing NRPS-A2_D7_D21. The Base Peak Chromatogram (BPC, black line) and the Extracted Ion Chromatogram (EIC, below with colours according to the depicted legend) of **1** (*m/z* $[M+H]^+$ = 598.42) and **11** (*m/z* $[M+H]^+$ = 470.35). The pACYC expression was used as a negative control (dotted line).

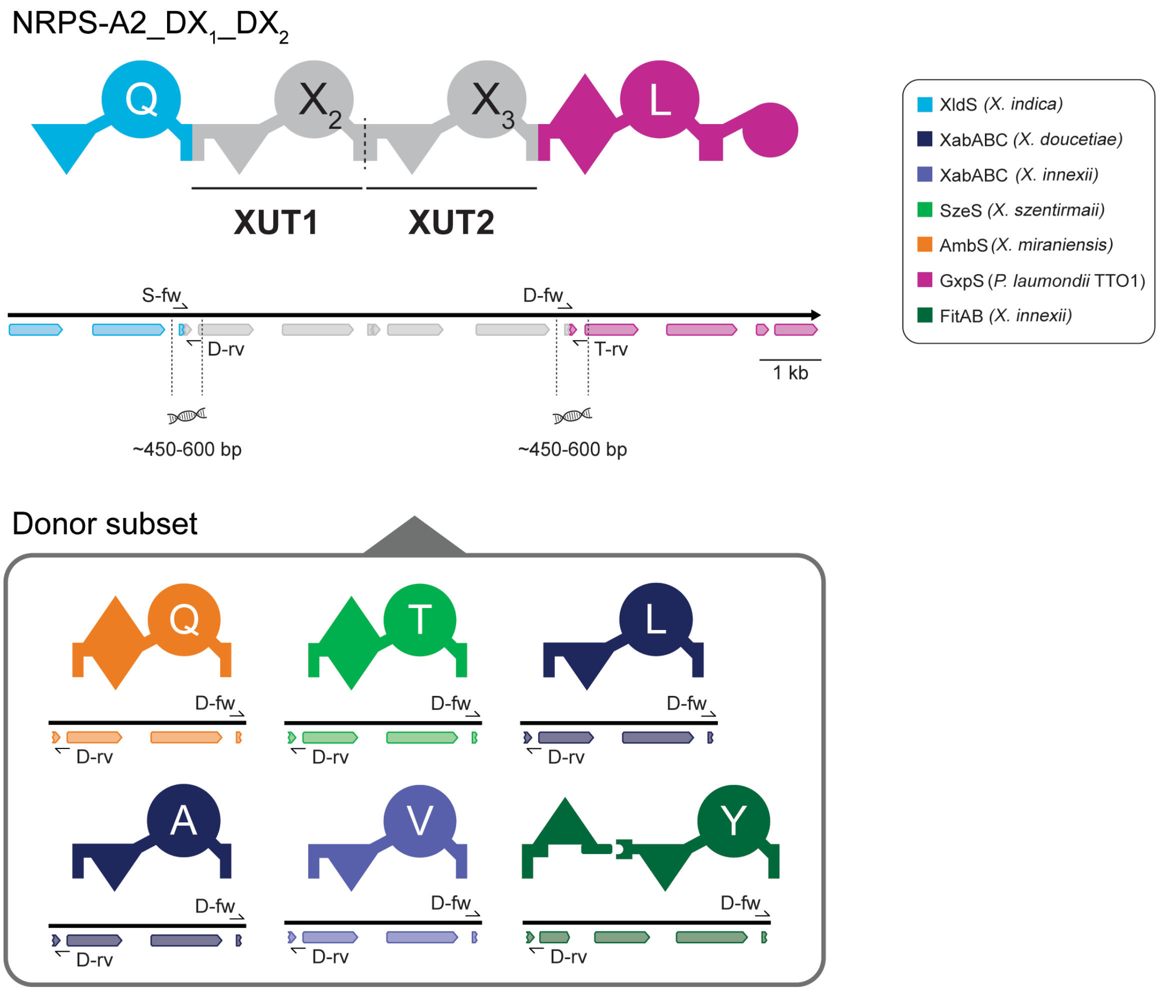


**Figure S8. Schematic overview of the primer allocation of the qPCR analysis for NRPS plasmid validation.** Domain annotations are included below the XUT. Acceptor plasmids contain a forward primer in the coding region of the starter module (S-fw) and a reverse primer in the coding region of the termination module (T-rv). Donors have both primers designed and allocated at the gene regions of the end of the A domain (D-fw) and within the T-C linker or the start of the C domain (D-rv). During qPCR, amplicons of ~450 – 600 bp are created.

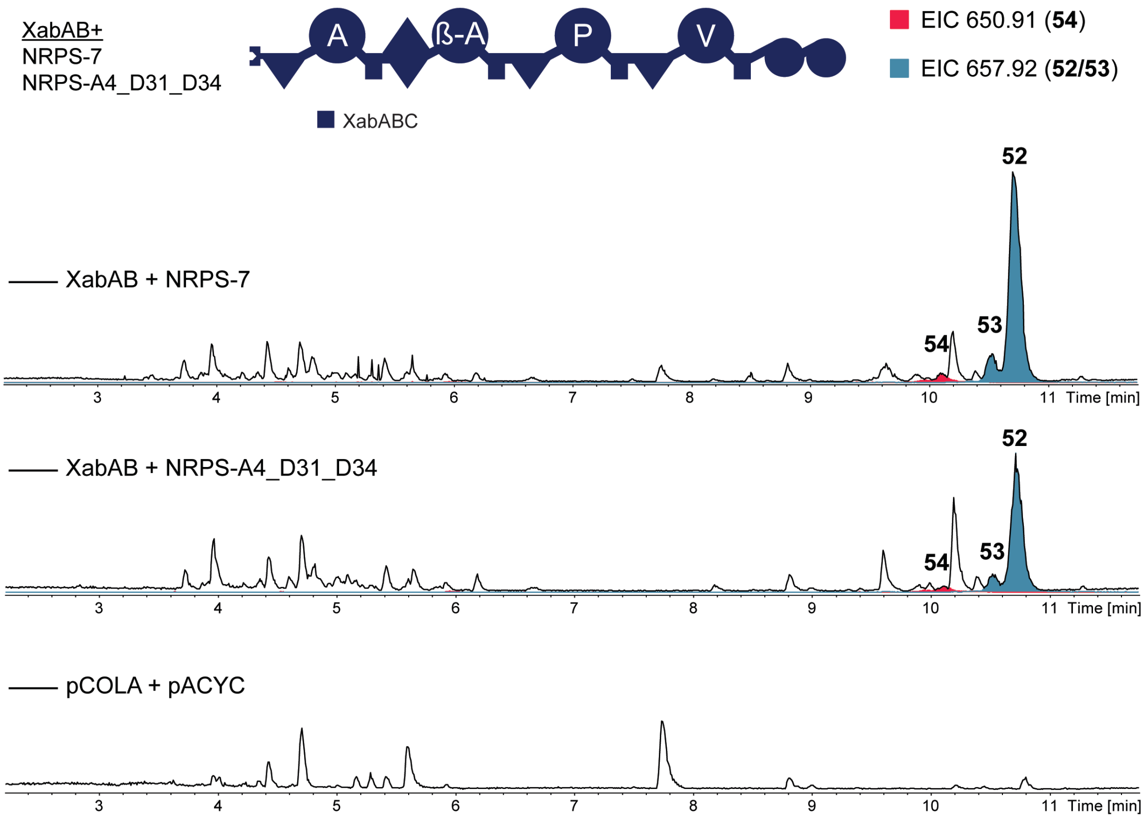


**Figure S9. HPLC/MS analysis of the produced NRPS-7 and NRPS-A4_31_D34 in *E. coli* DH10B*::mtaA*.** Schematic representation of NRPS-7 and NRPS-A4_31_D34 with the colour code of the original NRPS. The A domain specificities are indicated with A = alanine, β-A = beta-alanine, P = proline, and V = valine. HPLC/MS data of compounds **52** – **54** (xenoamicin A, B and C) produced in *E. coli* DH10B::*mtaA* co-expressing XabAB and NRPS-7 or XabAB and NRPS-A4_31_D34. The Base Peak Chromatogram (BPC, black line) and the Extracted Ion Chromatogram (EIC, below with colours according to the depicted legend) of **52** and **53** (*m/z* $[M+2H]^{2+}$ = 657.92) and **54** (*m/z* $[M+2H]^{2+}$ = 650.91). The pCOLA + pACYC expression was used as a negative control.

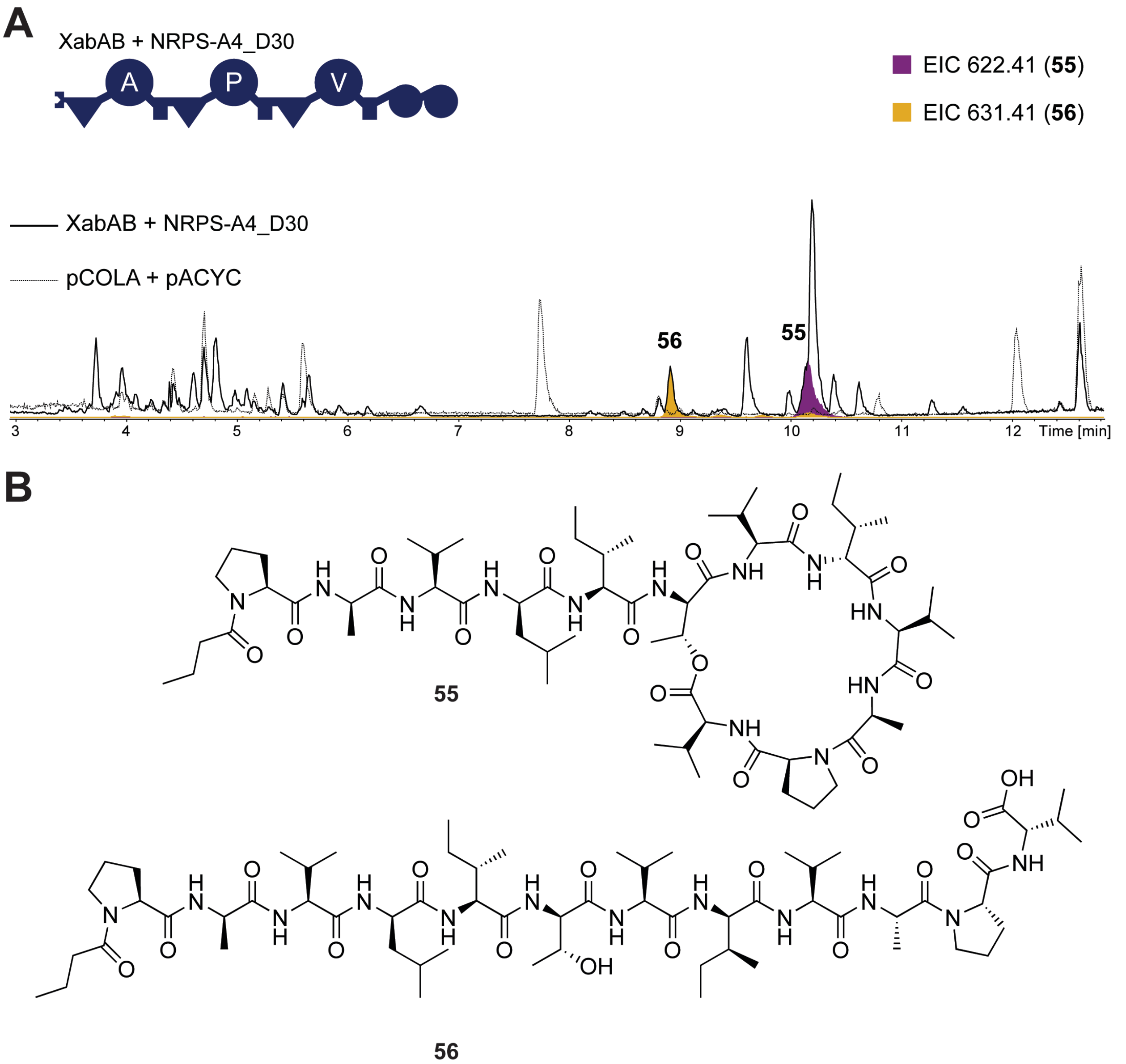


**Figure S3. HPLC/MS analysis of the produced NRPS-A4_D30 in *E. coli* DH10B*::mtaA*. A**) Schematic representation of NRPS-A4_D30 with the colour code of the original NRPS. The A domain specificities are indicated with A = alanine, P = proline, and V = valine. HPLC/MS data of compounds **55** (cyclic) and **56** (linear) produced in *E. coli* DH10B::*mtaA* expressing NRPS-A4_D30. The Base Peak Chromatogram (BPC, black line) and the Extracted Ion Chromatogram (EIC, below with colours according to the depicted legend) of **55** (*m/z* $[M+2H]^{2+}$ = 622.41) and **56** (*m/z* $[M+2H]^{2+}$ = 631.41). The pCOLA + pACYC expression was used as a negative control (dotted line). **B**) Chemical structure of the peptides **55** and **56**.
